# DeepEmbCas9: Cas9 coevolution and sgRNA structural information for CRISPR-Cas9 cleavage activity prediction

**DOI:** 10.1101/2025.10.08.681228

**Authors:** Jeffrey Mak, Peter Minary

## Abstract

The development of CRISPR-Cas9 cleavage activity prediction tools hinges on data produced from high-throughput guide-target lentiviral library screens for different Cas9 variants. However, the majority of such tools remain limited to predictions for one or few Cas9 variants, making it difficult to quantify the effects of Cas9 residues on cleavage activity. To bridge the gap, we introduce 4 interpretable DeepEmbCas9 models for the cleavage activity prediction of 40 type II-A and II-C Cas9 variants — DeepEmbCas9, DeepEmbCas9-MVE, DeepEnsEmbCas9 naive, and DeepEnsEmbCas9 — leveraging protein and RNA language model embeddings to encode Cas9 and sgRNA, respectively. Among the 4 neural network models, DeepEnsEmbCas9 naive performed the best in both in-distribution and out-of-distribution settings, where DeepEnsEmbCas9 naive outperformed individual Cas9 cleavage activity prediction tools on 18 out of 51 and 17 out of 48 benchmark test sets, respectively, and performed comparably otherwise. Concerning uncertainty quantification, DeepEnsEmbCas9 yields quantile-calibrated uncertainty estimates while keeping a minimal performance drop compared to DeepEnsEmbCas9 naive. SHAP importance analysis on DeepEmbCas9 reaffirms the importance of Cas9-target PAM binding as a first step for Cas9 cleavage, and reveals the L2 linker and PLL-WED-PI as important Cas9 domains modulating DeepEmbCas9’s predicted activity change when introducing increased-fidelity and PAM-altering Cas9 mutations, respectively. Our findings demonstrate the usefulness of protein language model embeddings in uncertainty-aware Cas9 cleavage activity prediction. More generally, DeepEmbCas9 models serves as an initial step towards cleavage activity prediction modelling for the whole Cas9 protein family.

## 1 Introduction

The repurposing of *Streptococcus pyogenes* Cas9 (SpCas9) — a programmable RNA-guided DNA endonuclease [1] — and other clustered regularly interspaced short palindromic repeats-associated protein 9 (CRISPR-Cas9, abbreviated Cas9) endonucleases from bacterial adaptive immune systems [2] for genome editing in mammalian cells [3] has revolutionized the field of gene therapy [4, 5], as evidenced by the Nobel Prize in Chemistry 2020 [6] and the U.S. Food and Drug Administration’s approval of Casgevy, the first CRISPR-Cas9 gene therapy for treating sickle cell disease, in 2024 [7, 8]. In essence, by programming the spacer sequence of a single guide RNA (sgRNA) for a user-selected Cas9 nuclease (typically 20-27nt), one can direct the Cas9-sgRNA binary complex to any genomic DNA target site of interest for editing of the (genomic) DNA target site [1, 9].

For the genomic target site to be edited, the CRISPR-Cas9 molecular machinery mechanistically operates in several steps, namely genome interrogation [10] for a protospacer-adjacent motif (PAM)-compatible target site, binding of Cas9’s PAM-interacting (PI) and Wedge (WED) domains to the PAM [11], target dsDNA unwinding stabilized by the phosphate lock loop (PLL) [12], R-loop formation coupled with repositioning of Cas9’s HNH domain next to the target strand (TS), concerted TS and non-target strand (NTS) cleavage (i.e, double-stranded break (DSB)) 3-4bp upstream of PAM by Cas9’s HNH and RuvC domains, and introduction of insertions and deletions (indels) to the target site via non-homologous end joining (NHEJ, i.e., error-prone endogenous DSB repair), and dissociation of Cas9-sgRNA from the edited target site. For SpCas9, the CRISPR-Cas9 complex adopts an inactive checkpoint conformation with rearranged REC2, REC3 and HNH domains upon partial sgRNA-DNA hybridization [13], and adopts a catalytically active conformation with HNH positioned next to the target strand [14, 15] once sgRNA-target hybridization is complete.

Despite SpCas9’s success in genome editing, the genome editor has several major limitations:

1. SpCas9 has variable on-target efficiency for different spacer sequences [16, 17];
2. SpCas9 has a propensity for off-target cleavage, dubbed the off-target effect [16, 18, 19, 20, 21], where SpCas9 may cleave unintended off-target sites due to its tolerance of base pair mismatches between the sgRNA spacer sequence and target dsDNA, especially PAM-distal mismatches [13];
3. SpCas9 cannot access a large portion of the human genome due to its NGG PAM requirement;
4. Because of SpCas9’s large gene size of 4.1kb, an all-in-one SpCas9-sgRNA expression cassette would exceed the ~4.7kb packaging limit [22] of an adeno-associated virus (AAV) [22] — a common approach for delivering Cas9 genome editors into mammalian cells [23].

Given a target site of interest, it also remains challenging to optimize the combination of Cas9 variant and sgRNA for efficient and specific genome editing. To overcome these issues, researchers have pursued four research directions: engineering of the SpCas9 nuclease [24, 25, 26, 27, 28, 29], metagenomic mining [30] and engineering [24, 31, 32, 33, 34, 35, 36] of smaller Cas9 orthologs with alternative PAMs, gRNA scaffold optimization [37], and the use of computational modelling of cleavage activity for optimal sgRNA design [38].

Enhanced efficiency and specificity can be achieved by mutating protein residues in SpCas9, as motivated by the extensive contacts between SpCas9 and the guide-target heteroduplex in the CRISPR-Cas9 complex. Using structure-guided rational engineering, researchers developed high-fidelity variants such as eSpCas9(1.1) [24], SpCas9-HF1 [25], and HypaCas9 [26]. Alternatively, researchers have used yeast-based directed evolution (DE) [39] to develop evoCas9 [27], and *E. coli*-based DE to develop Sniper-Cas9 [28], Sniper2P and Sniper2L [29]. Sc++ [40] was also rationally engineered from *Streptococcus canis* Cas9 (ScCas9) via multiple sequence alignment between ScCas9 and closely related *Streptococcus* orthologs. As for the broadening of SpCas9’s PAM requirement, structure-guided rational engineering gave rise to SpCas9-VQR (VQR onwards) [41], SpCas9-VRER (VRER onwards) [41], SpCas9-VRQR (VRQR onwards) [25], VRQR-HF1 [25], QQR1 [42], SpCas9-NG [43], SpG [44], and SpRY [44], whereas phage-assisted (non-)continuous evolution [45, 46, 47] gave rise to xCas9(3.7) (referred to as xCas9 onwards) [48], SpCas9-NRRH, SpCas9-NRTH, and SpCas9-NRCH [49].

As mentioned earlier, fitting an all-in-one SpCas9-sgRNA expression cassette into an AAV vector is not feasible. Early approaches addressed this issue by identifying smaller Cas9 orthologs that show mammalian genome editing activity, which included St1Cas9 [41, 50, 51, 52, 53], Nm1Cas9 [50, 54, 55, 56], SaCas9 [57, 58, 41, 59, 60, 61], CjCas9 [62, 63], Nm2Cas9 [64], SlugCas9 [36, 33] and SauriCas9 [65]. However, these nucleases lack specificity, so increased fidelity variants such as eSaCas9 [24], efSaCas9 [31], SaCas9-HF [32], SlugCas9-HF [33], and enCjCas9 [34] were developed. The nucleases also have lengthy PAMs, so PAM-relaxed variants such as SaCas9-KKH [35] and SauriCas9-KKH [65]. Further protein engineering gave SaCas9-KKH-HF [32], Sa-SlugCas9 [33], and sRGN3.1 [36], which were developed by stacking SaCas9-KKH and SaCas9-HF mutations, replacing SaCas9’s PI domain with that of SlugCas9, and shuffling of protein fragments from Cas9 orthologs similar to SlugCas9, respectively.

Changes to the gRNA scaffold also boosted on-target activity. For SpCas9, this involved increasing the length of the repeat-antirepeat and a T-to-C mutation that broke the 5’ continuous stretch of thymines which was misinterpreted as a transcription termination signal. Various SaCas9 [58, 60, 35], St1Cas9 [50, 52], NmCas9 [45, 54, 64], and CjCas9 [62, 63] gRNA scaffolds have also been considered for boosting the on-target efficiency of small Cas9 nucleases.

CRISPR-Cas9 cleavage activity prediction models have become cheap *in silico* alternatives to the low-throughput and costly *in vitro* and *in vivo* surrogate reporter [66, 67] and gold-standard indel frequency measurement assays [68, 69, 70] typically required for optimal sgRNA design. Unlike early pioneering work which employed rule-based algorithms [71, 72, 16, 73, 74] and machine learning (ML) [17, 75, 76, 77, 78], modern data-driven approaches use deep learning [79, 80, 81, 82, 83] for activity prediction by leveraging Cas9 (off-)target cleavage indel frequency data from *in vitro*/*vivo* genome-wide off-target cleavage detection [84, 85, 86, 87, 88, 89, 90, 91, 92] and high-throughput guide-target lentiviral library screens [93, 94, 95]. Notably, DL models are favored over ML models due to their ability to perform automatic feature extraction and superior predictive performance [95, 96] over ML models. A variety of neural network architectures have been used, including convolutional neural networks (CNN) [97, 98, 99], recurrent neural networks (CNN) [100], convolutional-recurrent neural networks (C-RNN) [101, 102, 103, 104], and kinetically interpretable neural network (KINN) [105] for Cas9 cleavage activity prediction. Founded on the fact that R-loop formation is the rate-limiting step in Cas9 cleavage, modern approaches also include thermodynamic free energy-based models [106, 107, 108], kinetic models [109, 110], and molecular dynamics (MD)-derived feature-based ML [111, 112] models.

Since the spacer and target sequences form the primary determinants of Cas9 cleavage activity [16], the majority of ML- and DL-based models have relied on representations of the spacer-target interface for input features to the neural network. More concretely, such features include one-hot encoding of the spacer-target interface [113], GC counts, computed DNA melting temperatures and computed minimum free energy of the sgRNA [114]. Given that cleavage activity can be modulated by epigenetics [115], specifically primary chromatin structure [116, 117, 118, 119, 120], some models [121, 101] also incorporate computed and/or epigenetic features as model inputs.

Despite the plethora of ML/DL-based Cas9 cleavage activity tools, the majority of them were trained only on a single Cas9 nuclease (Figures S1 and **??**). As a result, such tools cannot account for the impact of the Cas9 nuclease on cleavage activity. STING CRISPR [112] tried to addresses this by predicting SpCas9 cleavage activity from molecular dynamics (MD)-derived residue-level physicochemical/structural descriptors for 27 SpCas9 spacer-target interfaces, but remains impracticable due to the large computational cost of protein-nucleic acid MD simulations. Uni-deepSG [122] is a DL model trained on SpCas9, eSpCas9(1.1) and SpCas9-HF1, but indirectly captures differences between SpCas9 nucleases in two non-trivial dataset-trained parameters. As part of machine learning-assisted directed evolution (MLDE) for Cas9 nucleases, Thean et al. [123] trained Cas9-activity ML models for multiple SpCas9 or SaCas9-KKH guide-target interface using protein embeddings from Georgiev et al. [124] and Bepler et al. [125], despite these models not being able to generalize to unseen guide-target interfaces. Indirectly related to spacer-target cleavage activity prediction, PAMmla [126] takes a 4nt PAM sequence and a one-hot encoding of 6 PAM-interacting residues and returns the cleavage rate for the specified SpCas9 variant and PAM. Similarly, CICERO [127] leverages ESM2 embeddings for Cas9 protein PAM prediction. Riding the recent success of protein language models (pLM) [128, 129, 130, 131, 132, 133], PLM-CRISPR [96] leverages ESM2 [134] embeddings and DL for SpCas9 variant cleavage activity prediction, but is only limited to NGG PAM on-target interfaces and 7 increased-fidelity SpCas9 variants. Though not related to activity prediction, pLM and genomic language models (gLM) have also been used for CRISPR-Cas protein classification [135] and *de novo* Cas9 protein design [136, 137, 132]. The field has also seen tools, e.g., CCLMoff [138], which use RNA language model (rLM, i.e., RNA foundation models) embeddings [139, 140] for SpCas9 cleavage activity prediction.

In light of the above, we introduce a family of 4 DeepEmbCas9 models — the first set of DL models to featurize all three components (sgRNA, DNA, Cas9) in the CRISPR-Cas9 complex. DeepEmbCas9 models introduce a new guide-target encoding scheme that enables the unification of indel frequency data from various Cas9 orthologs and their variants. The latter enables the utilization of the largest guide-target lentiviral library-based CRISPR-Cas9 indel frequency dataset, which has over 1.75 million datapoints spanning 40 Cas9 variants and 16 gRNA scaffolds. It also makes DeepEmbCas9 models applicable to more Cas9 variants compared to PLM CRISPR. In terms of neural network architecture, DeepEmbCas9 models use inductive biases informed by Cas9’s biophysical mechanism, where sgRNA-DNA interactions motivate the use of early fusion of nucleic acid sequence/energy/structural information, and Cas9-PAM interactions motivate the use of early fusion of Cas9 pLM embedding and PAM sequence. The use of concatenated region-wise pLM mean embeddings for encoding Cas9 variants (further referred to as Cas9 pLM embeddings onwards) enable DeepEmbCas9 models to parallel STING CRISPR’s ability to assess the impact of Cas9 domains on activity. Analogously, concatenated region-wise rLM mean embeddings for encoding sgRNAs (further referred to as sgRNA rLM embeddings) enables one to assess the impact of various parts of the sgRNA (scaffold) on activity. Compared to PLM-CRISPR, DeepEnsEmbCas9 naive – the flagship model among the four models — has comparable test performance when benchmarked against existing single-variant deep learning-based indel frequency prediction tools in both the in-distribution and leave-one-nuclease-out extrapolation settings. Equipped with deep ensembles of mean-variance estimators, DeepEnsEmbCas9 captures both aleatoric and epistemic uncertainty, yielding calibrated uncertainties and good test performance. SHAP importance analysis on DeepEmbCas9 emphasizes on the importance of CRISPR-Cas9 PAM binding on cleavage activity prediction. Comparing between 39 nuclease pairs, DeepEmbCas9 is seen to regard Linker and PLL-WED-PI domains as being important for nuclease pairs containing increased-fidelity and PAM-altering mutations, respectively. Taken together, DeepEmbCas9 models form the first step towards a general CRISPR-Cas9 activity model capable of making accurate prediction for any Cas9 variant available in the literature.

## 2 Methods

Since different studies use different nuclear localization signals (NLS), we distinguish between nucleases and variants, where nucleases refer to the Cas9 domains, and variants refer to the Cas9 domains together with the NLS, FLAG and P2A components. For convenience, the term “variants” also include wild-type nucleases.

### 2.1 Dataset construction

To build a CRISPR-Cas9 activity prediction tool spanning multiple Cas9 variants and gRNA scaffolds, we built a dataset by combining high-throughput guide-target lentiviral library-based indel frequency data from 6 studies [100, 95, 141, 142, 114, 29] (see Table 1). A list of column descriptions for the dataset can be found in Table S1, noting that the primary keys of the dataset are “Spacer sequence”, “Target context sequence”, “Variant”, “gRNA scaffold”, “Day” and “tRNA feature”, the tuple of which forms a unique experimental configuration. Frequency counts on the number of mismatches among guide-target interfaces in the dataset can be found in Table S2. The names of the gRNA scaffold in the dataset are identical to those used in Seo et al. [114], apart from naming of SpCas9 gRNA scaffolds, where “SpCas9 scaffold 1” is the standard unoptimized scaffold with a 6nt poly(U) tail, “SpCas9 scaffold 1 (5T)” is the standard unoptimized scaffold with a 5nt poly(U) tail, and “SpCas9 scaffold 2” is the optimized scaffold described by Dang et al. [37]. To avoid over-representation of experimental configurations with multiple indel frequency measurements within a study and/or across studies, e.g., due to biological replicates, we average indel frequencies within each study and across the whole dataset to build per-study and cross-study ML-ready labels with values ranging up to 100%. Experimental indel frequencies from Wang et al. [100] were also scaled up 100*×* to match the indel frequency ranges used in the other studies.

**Table 1.** List of studies used for curating the Cas9 variant indel frequency dataset consisting of 1.75 million points spanning 40 Cas9 variants and 16 gRNA scaffolds, in addition to corresponding tools used as baselines for this study. Indel frequencies were aggregated across studies such that a datapoint is uniquely identified by its spacer, target, gRNA scaffold, Cas9 variant, number of days post-transduction/transfection, and use of tRNA^Gln^ preprocessing. Numbers enclosed within parentheses in the last column indicate the percentage of unique datapoints which contain spacer-target mismatches, either as a mismatched 5’ guanine or mismatches elsewhere in the heteroduplex.

| Study | Published tools | Cas9 variants | gRNA scaffolds | No. of unique datapoints<br>(% mismatched) |
| --- | --- | --- | --- | --- |
| Kim, Kim et al. (2019) [95] | DeepSpCas9 | SpCas9 | SpCas9 scaffold 1 | 13,359 (72.89%) |
| Wang et al. (2019) [100] | DeepHF | SpCas9, eSpCas9(1.1), SpCas9-HF1 | SpCas9 scaffold 1 (5T) | 171,109 (0%) |
| Kim et al. (2020) [141] | DeepxCas9, DeepSpCas9-NG | SpCas9, xCas9, SpCas9-NG | SpCas9 scaffold 1 | 88,786 (19.04%) |
| Kim, Kim et al. (2020) [142] | DeepSpCas9variants | HypaCas9, Sniper-Cas9, SpCas9-HF1,<br>SpCas9, eSpCas9(1.1), evoCas9,<br>xCas9, SpCas9-NG, VRQR,<br>QQR1, VQR, VRER, VRQR-HF1 | SpCas9 scaffold 2 | 318,136 (51.76%) |
| Seo et al. (2023) [114] | DeepSmallCas9 | SpCas9,<br>SaCas9, SaCas9*, SaCas9-KKH,<br>SaCas9-HF, SaCas9-KKH-HF, eSaCas9,<br>efSaCas9, SauriCas9, SauriCas9-KKH,<br>SlugCas9, SlugCas9-HF,<br>Sa-SlugCas9, sRGN3.1<br>CjCas9, enCjCas9,<br>Nm1Cas9, Nm2Cas9,<br>St1Cas9, | SpCas9 scaffold 2,<br>SaCas9 scaffolds 1-3,<br>CjCas9 scaffolds 1-2,<br>NmCas9 scaffolds 1-3,<br>St1Cas9 scaffolds 1-5 | 713,163 (73.56%) |
| Kim, Kim, Okafor et al. (2023) [29] | DeepSniper | Sniper-Cas9, Sniper2P, Sniper2L | SpCas9 scaffold 1 | 127,034 (51.80%) |
| Kim, Choi et al. (2024) [143] | DeepCas9variants | SpRY, Sc++, SpCas9-NRCH,<br>SpCas9-NRRH, SpCas9-NRTH,<br>SpG, SpCas9, SpCas9-NG,<br>VRQR, xCas9 | SpCas9 scaffold 2 | 205,358 (59.04%) |
| Kim, Kim et al. (2020) [142]<br>and Kim, Choi et al. (2024) [143] | DeepSpCas9variants,<br>DeepCas9variants | SpCas9-NG, SpCas9, VRQR, xCas9 | SpCas9 scaffold 2 | 109,741 (56.06%) |
| Total | 8 tools | 40 Cas9 variants | 16 gRNA scaffolds | 1,746,686 (55.22%) |

Cas9 and guide RNA (gRNA) scaffold sequences were also curated as part of the dataset. The 16 gRNA scaffold sequences were derived from sequences in Supplementary Table 9 of Seo et al. [114], noting that Wang et al. [100] uses “SpCas9 scaffold 1 (5T)”. Protein and codon sequences of the 40 Cas9 variants (including the nuclear localization sequence, FLAG tag and P2A peptide) were obtained from Addgene and supplementary files of the original studies (additional details available in supplementary materials section “Dataset” subsection “Dataset construction”).

### 2.2 Input feature encodings

The dataset was preprocessed in order to construct deep learning-ready data representations of components in the guide-target-Cas9 variant R-loop complex (Figure S4A). Illustrated in Figures 2A and S4B, a one-hot representation of the spacer and target context sequence was built by stacking of one-hot encodings of the padded spacer and target context sequences, where the padded sequences were constructed by flanking the raw spacer and target context sequences with padding (gray), represented by the letter ‘N’ in both sequences, such that:

**Figure 1.**
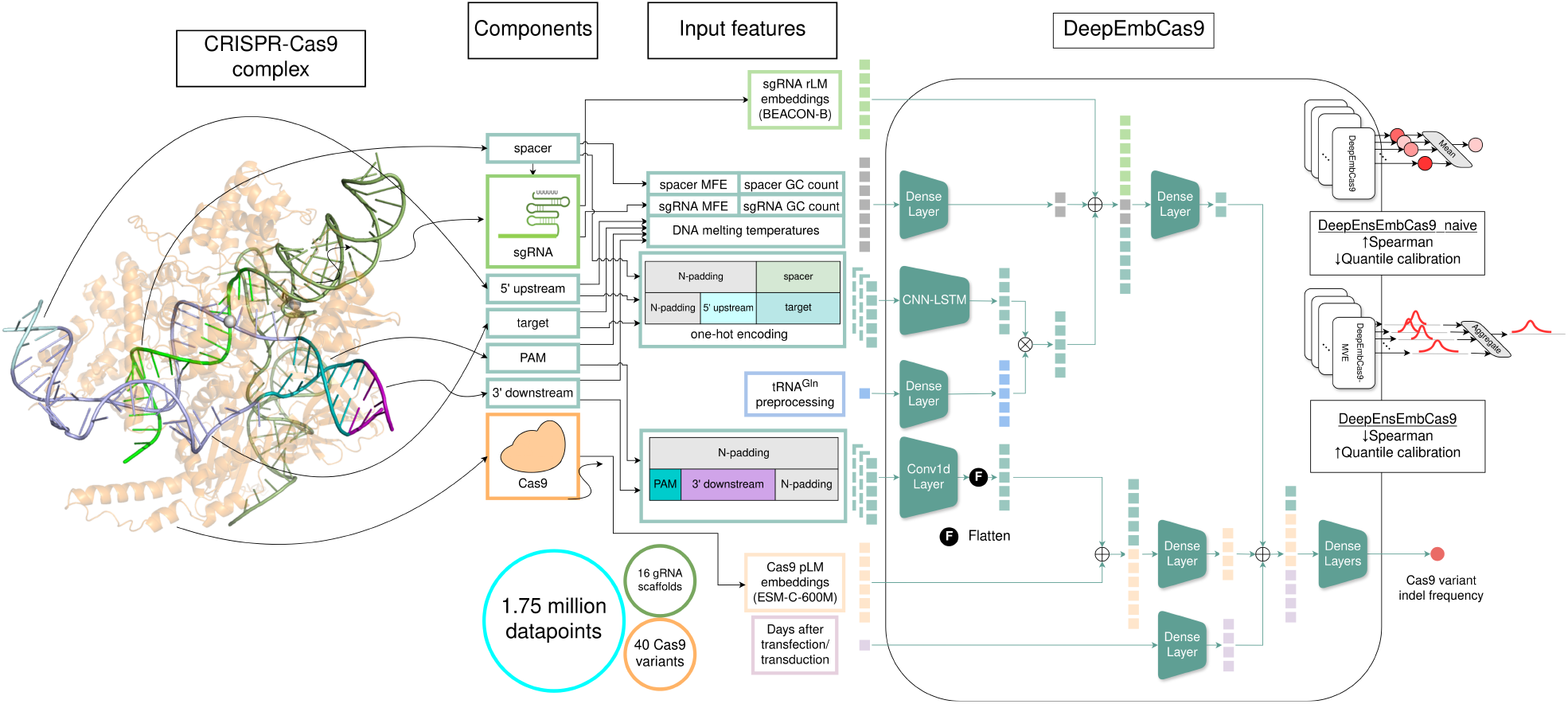
Overview of DeepEmbCas9. All components of the CRISPR-Cas9 complex are featurized and fed as input to DeepEmbCas9. DeepEmbCas9, a biophysically inspired neural network, was developed using a dataset with 1.75 million datapoints spanning 40 Cas9 variants and 16 gRNA scaffolds. Ensembling of neural networks with and without mean-variance estimation yields DeepEnsEmbCas9 and DeepEnsEmbCas9 naive.

**Figure 2.**
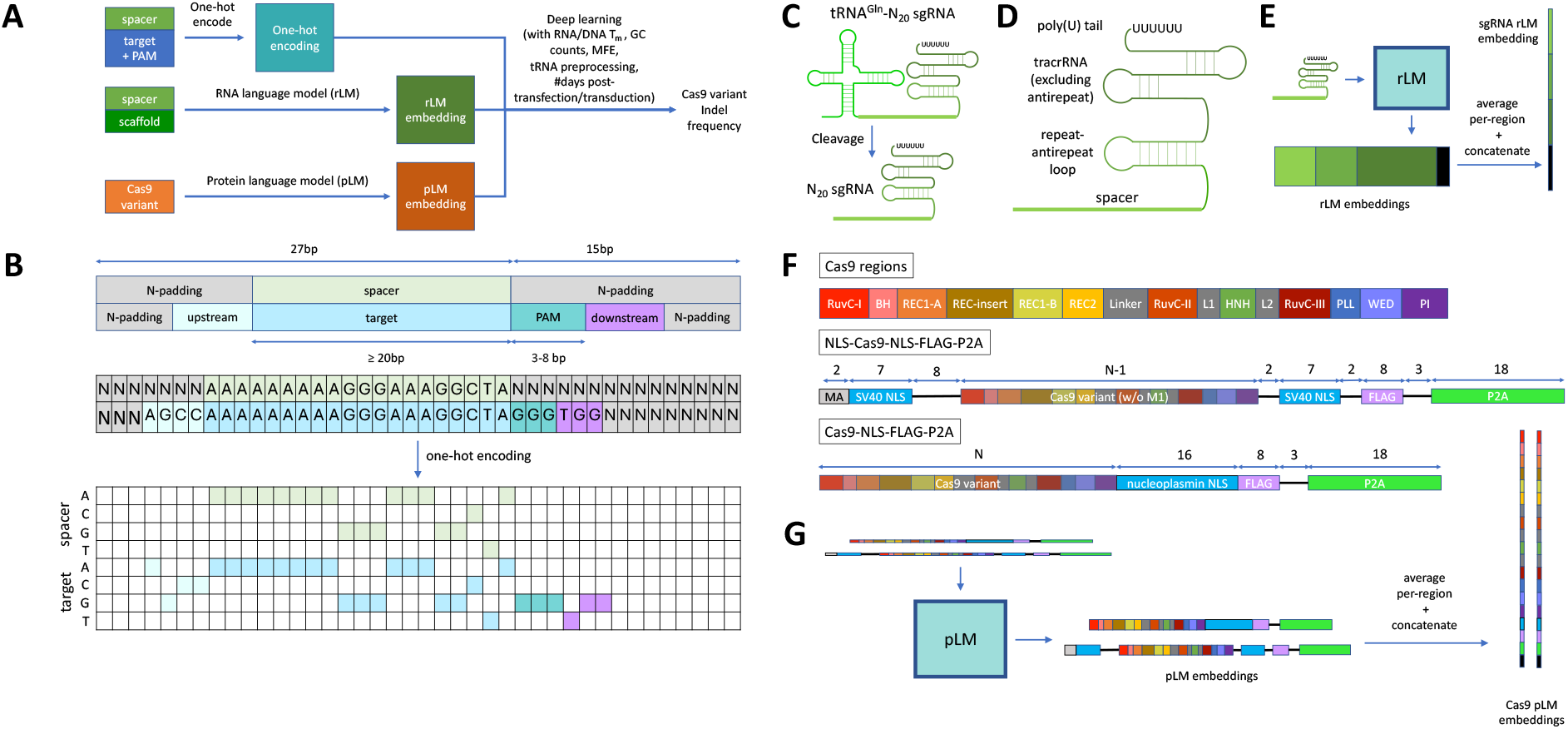
Conceptual overview of DeepEmbCas9 (A), and featurization of the full CRISPR-Cas9 complex (B-G). (B) Unified guide-target interface (top) with an example matched SpCas9 interface (middle) and its one-hot encoding (bottom). (C) Formation of perfectly matching N_20_ sgRNAs via tRNA^Gln^-N_20_ preprocessing (for SpCas9 variants only). (D) The four sgRNA regions used in this study, namely the spacer (light green), repeat-antirepeat loop (green), tracrRNA excluding the antirepeat sequence (dark green) and poly(U) tail (black). (E) concatenated region-wise rLM mean embeddings for encoding sgRNA (sgRNA rLM embedding) are formed by passing sgRNA sequences to RNA language models (rLMs), averaging per-nucleotide embeddings in the sgRNA regions, and concatenation of the averaged embeddings in the feature dimension. (F) Protein regions (color-coded) used for constructing the concatenated region-wise pLM mean embeddings for encoding Cas9 variants (Cas9 pLM embedding). These include regions within a Cas9 variant (top, of length *N*), as well as nuclear localization signals (NLS, sky blue), FLAG-tag (FLAG, light purple), self-cleaving P2A peptide (neon green) and other inter-region residues (black text and lines), as seen in the NLS-Cas9-NLS-FLAG-P2A (middle) and Cas9-NLS-FLAG-P2A (bottom) sequences. (G) Cas9 pLM embeddings are formed by passing Cas9 sequences to protein language models (pLMs), averaging per-residue embeddings in the Cas9 regions, and concatenation of the averaged embeddings in the feature dimension. Inter-region residues are pooled into the “Other” region in black.

- both padded sequences have length 42;
- heteroduplex nucleotides in both padded sequences are positionally aligned; and
- the PAM sequence starts at the 28th nucleotide in the padded target context sequence.

In the one-hot encoding, ‘N’ maps to the zero vector, i.e., ‘N’ becomes zero-padding. Values in the “tRNA feature” and “Day” dataset columns are stored as a binary value and a real number, respectively, where a visual depiction of tRNA^Gln^ preprocessing is shown in Figure 2B.

Similar to Seo et al. [114], melting DNA temperatures, RNA minimum free energy (MFE) and GC counts were calculated for all guide-target interface in the dataset. Tm NN with default parameters from Biopython’s Bio.SeqUtils package [144] (https://biopython.org/docs/dev/api/Bio.SeqUtils.html) was used for computing DNA melting temperatures for 39 subsequences of the target context sequence, namely those with zero-indexed ranges [0, 5), [5, 14), [14, 20), [20, 27), [0, 7), [7, 18), [18, 22), [22, 27), [0, 3), [3, 10), [10, 17), [17, 27), [0, 4), [4, 12), [12, 19), [19, 27), [4, 11), [11, 17), [0, 6), [6, 15) and [15, 20) for the 5’ upstream and protospacer parts of the target context sequence, and [27, 30), [30, 33), [27, 34), [34, 36), [27, 36), [36, 38), [27, 34), [34, 37), [27, 33), [33, 36), [27, 35), [35, 38), [27, 33), [33, 38), [27, 31), [31, 34), [27, 30) and [30, 38) for the PAM and 3’ downstream parts of the target context sequence. DNA melting temperatures for ‘N’-containing subsequences were handled by replacing ‘N’ with ‘G’ (i.e., guanine) prior to MFE calculations. ViennaRNA’s Python API [145] was used for calculating the minimum free energy (MFE) of the spacer and sgRNA (spacer + scaffold) sequences, with T replaced with U before MFE calculations. Bio.SeqUtils’s gc fraction was used for calculating GC counts of the spacer and protospacer sequences.

### 2.3 sgRNA regions

We define 4 non-overlapping sgRNA regions to structurally align the 16 gRNA scaffolds considered in this study (Figure 2C). The four regions are

- spacer sequence;
- repeat-antirepeat loop, consisting of the crRNA repeat sequence, GAAA tetraloop and tracrRNA antirepeat sequence;
- tracrRNA with the antirepeat excluded; and
- poly(U) tail, typically the last 5-7 ribonucleotides in the sgRNA,

Boundaries delineating the regions for the gRNA scaffolds were determined via manual inspection (see Table S3 for sgRNA region lengths for each gRNA scaffold).

### 2.4 sgRNA rLM embeddings

From sgRNA sequences in the dataset, per-nucleotide rLM embeddings were generated from RNAFM [139], RiNALMo [146], BEACON-B [140] and BEACON-B512 [140] using a 32GB V100 NVIDIA GPU. Since genomic language models potentially encode RNA-related information, per-nucleotide genomic language model embeddings were generated from evo-1-8k [137] using a 40GB A100 NVIDIA GPU. rLM/gLM embeddings were then converted into sgRNA rLM embeddings by averaging (ribo)nucleotide embeddings within each sgRNA region, followed by concatenation of the sgRNA region embeddings in the feature dimension (Figure 2D).

### 2.5 Cas9 regions

We use Cas9 regions, i.e., contiguous protein segments in the Cas9 nuclease, to structurally align the 40 Cas9 variants (Figures 2E top and S3B). The term “region” is used to distinguish from “domain”, which is defined as one or more noncontiguous protein segments. To clarify, “REC-insert” (Figure S3, beige-colored) denotes the domain inserted into the REC1 domain, i.e., Wing in St1Cas9, and REC2 domains in SpCas9 and ScCas9 variants. “REC1-A” denotes the first REC domain in the Cas9 nuclease, i.e., REC1-A in SpCas9, St1Cas9 and ScCas9 variants, REC1 in Nm1Cas9 and Nm2Cas9, and REC in the other small Cas9 variants. Boundaries delineating Cas9 regions for the 40 Cas9 variants were gathered from literature and predicted from multiple sequence alignment (MSA) of wild-type Cas9 nucleases. Specifically,

- SpCas9 regions were obtained from Huai et al. [147], with the additional REC3-linker loop boundary at residue 712 from Jiang et al. [148];
- SaCas9* and SaCas9 regions were obtained from Nishimasu et al. [57], with SaCas9 having inserted glycine at position 2 (the same applies for other SaCas9 variants);
- St1Cas9 regions were obtained from Zhang et al. [53];
- SlugCas9, SauriCas9, Sa-SlugCas9 regions were obtained from Hu et al. [33];
- CjCas9 regions were obtained from Yamada et al. [63]; and
- Nm1Cas9 and Nm2Cas9 regions were obtained from Sun et al. [149].

Region boundaries with unknown positions were predicted through MSA of Cas9 nucleases, followed by projection of the region boundary from a similar nuclease to the target nuclease via the MSA.

Specifically, we built an MSA consisting of SpCas9, SaCas9*, SaCas9, St1Cas9, sRGN3.1, SlugCas9, SauriCas9, Sa-SlugCas9, CjCas9, Nm1Cas9, Nm2Cas9, ScCas9 and 8313 other type II CRISPR RNA-guided endonuclease Cas9 of length *>* 800 from UniRef100 [150] using Clustal Omega [151, 152] program. Using the MSA, we projected SpCas9’s WED start position to St1Cas9, Nm1Cas9 and Nm2Cas9, which resulted in predicted WED start site positions at 831, 851, and 850, respectively. ScCas9’s region boundaries were generated by using the same MSA and projecting all SpCas9 region boundaries to ScCas9. sRGN3.1’s region boundaries were generated by aligning ShyCas9, SmiCas9, SpaCas9, SlugCas9 and sRGN3.1 via Clustal Omega, followed by projection all SlugCas9 region boundaries to sRGN3.1. The resulting sets of Cas9 region boundaries are visualized in Figure S3, noting that Cas9 variants sharing the same base nuclease (as indicated in Table S4) share the same region boundaries.

In addition to the contiguous Cas9 regions, we devised four non-contiguous regions — NLS, FLAG, P2A and Other — to account for Cas9 nuclease-flanking residues which are part of the nuclear localization signal (sky blue), FLAG tag (light purple), P2A peptide (neon green) and other residues not part of the aforementioned regions (black horizontal lines and “MA” text at the N-terminal of NLS-Cas9-NLS-FLAG-P2A), respectively (Figure 2E middle and bottom). More concretely, NLS covers the nucleoplasmin NLS and two SV40 NLS sequences in Cas9-NLS-FLAG-P2A and NLS-Cas9-NLS-FLAG-P2A, repectively.

### 2.6 Cas9 pLM embeddings

From the 40 Cas9 sequences in the dataset, per-residue pLM embeddings for each sequence were generated from ProtT5 (specifically Rostlab/prot t5 xl half uniref50-enc) [130], Ankh-large [131], ESM3-1.4B [129, 128], ESM-C-300M [133, 128], ESM-C-600M [133, 128], ESM-C-6B [133, 128] using a 16GB V100 NVIDIA GPU (Figure 2F). Since genomic language models (gLM) potentially encode protein-related information, per-nucleotide genomic language model embeddings were also generated from gLM2 650M [153] for codon sequences corresponding to the 40 Cas9 variants. pLM/gLM embeddings were then processed into Cas9 pLM embbedings by averaging residue/nucleotide embeddings within each Cas9 region, followed by concatenation of the Cas9 region embeddings in the feature dimension. We did not use ESM2 [134] to generate Cas9 pLM embeddings, as the length of SpCas9 exceeds ESM2’s context window size of 1024 (i.e., maximum amino acid sequence length of 1022).

### 2.7 Neural network architecture and training

DeepEmbCas9 is a deep learning model that predicts the indel frequency for a given sgRNA, target context sequence, Cas9 variant with known structural domain annotations, and time since transfection/transduction of the sgRNA and Cas9 plasmids. DeepEmbCas9’s neural network architecture and relevant hyperparameters are detailed in Figure 3. To build DeepEmbCas9, we reused design ideas from existing deep learning-based CRISPR-Cas9 activity prediction models. Notably,

- design of the guide-target branch (consisting of convolutional and bidirectional long short-term memory (biLSTM) layers) and Cas9-PAM branch was inspired by the sequence and epigenetic arms from crispAI [154], respectively;
- element-wise multiplication operation between the “tRNA^Gln^ preprocessing” and spacer-target embeddings for feature fusion was adopted from DeepSpCas9variants [142]; and
- concatenation of the guide embedding, target embedding, DNA melting temperature features, GC counts, MFEs and mismatch encodings before the fully connected layers was adopted from DeepSmallCas9 [114].

**Figure 3.**
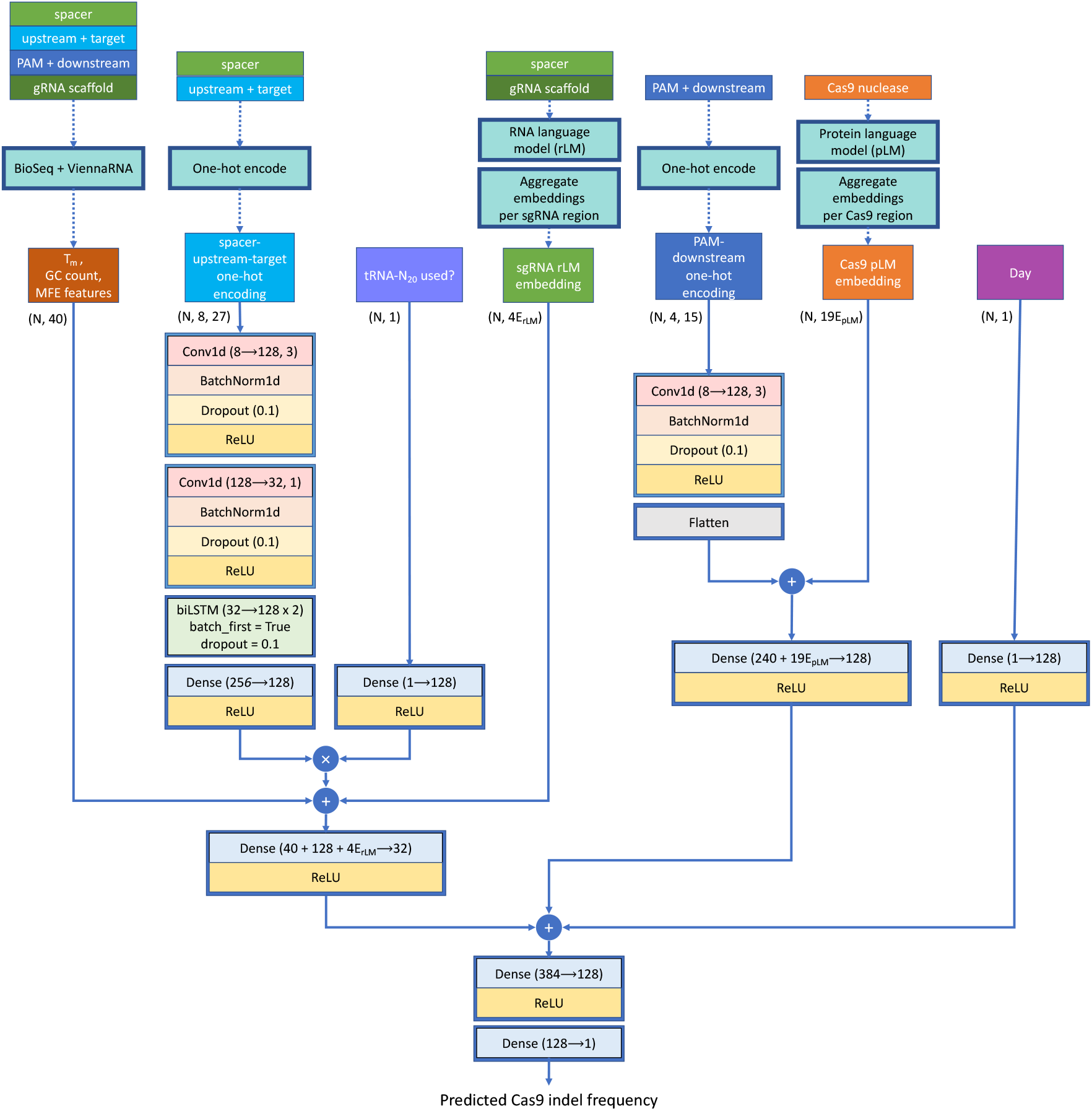
DeepEmbCas9’s neural network architecture. DeepEmbCas9 mainly consists of a spacer-target branch (left side) and a Cas9-PAM branch (right side). The sgRNA’s rLM embedding (green), tRNA^Gln^ preprocessing feature (cornflower blue), and BioSeq/ViennaRNA-derived RNA/DNA melting temperature (*T*_*m*_)/GC count/minimum free energy (MFE) features (brown) are integrated into the spacer-target branch, whereas the Cas9 variant’s pLM embedding (orange) is integrated into the PAM branch. The Day feature (purple) is integrated later in the neural network via the rightmost branch. Different branches in the neural network are fused together via concatenations and dense layers. Dotted arrows indicate the preprocessing steps used for generating DeepEmbCas9’s input feature matrices/tensors. Conv1d(*x* → *y, z*) denotes a 1D convolutional layer with *x* input channels, *y* output channels, *z* kernel size and same padding, whereas Dense(*x* → *y*) denotes a fully connected layer with *x* input and *y* output nodes. Blue circle with + and *×* denotes tensor concatenation and element-wise multiplication, respectively. *N* denotes the number of datapoints used in a forward pass through DeepEmbCas9.

As PAM recognition and binding by Cas9 are required for R-loop formation and double strand DNA cleavage [148], we used separate feature extractors for nucleotides upstream of the PAM (including heteroduplex nucleotides) and nucleotides within or downstream of the PAM into two separate neural network arms, respectively, with the Cas9 pLM embedding as input in the Cas9-PAM branch. We also chose to integrate the “Day” feature in the later layers of the fully connected layers, as the feature is unrelated to the CRISPR-Cas9 complex.

DeepEmbCas9 was implemented in PyTorch [155] and PyTorch Lightning [156]. To train DeepEmbCas9, we used mean squared error (MSE) as the loss objective, Adam [157] with learning rate 1 *×* 10^*−*3^ as the optimizer and `StepLR` with `gamma=0.1` and `step_size=1` as the learning rate scheduler. To avoid overfitting, `EarlyStopping` from PyTorch Lightning with default parameters was used, which halted training after no improved validation loss for three consecutive epochs. The model weights for the training epoch with the lowest validation loss is then saved. Default neural network weight initializations were used in DeepEmbCas9. All DeepEmbCas9 models were trained on a single 16GB V100 NVIDIA GPU for no more than 4 hours.

### 2.8 Benchmark comparisons

#### 2.8.1 Benchmark activity prediction tools

We opt to use deep learning tools published in the six studies listed in Table 1 as baselines. Specifically, DeepHF [100] is a set of 4 bidirectional LSTM models:

- DeepHF T7-SpCas9 and DeepHF SpCas9 predict SpCas9 activity;
- DeepHF eSpCas9(1.1) predict eSpCas9(1.1) activity; and
- DeepHF SpCas9-HF1 predict SpCas9-HF1 activity

for matched A/GN_19_ interfaces. DeepSpCas9 [95] is a convolutional neural network (CNN) model which predicts SpCas9 activity for matched G/gN_19_, i.e., interfaces with a matched 5’ guanine (GN_19_) or mismatched 5’ guanine (gN_19_). Similar to DeepSpCas9, DeepxCas9 and DeepSpCas9-NG [141] are CNN models which predict xCas9 and SpCas9-NG activity, respectively, for matched G/gN_19_ interfaces.

DeepSpCas9variant [142] is a set of 9 CNN models accepting matched G/gN_19_ and tRNA^Gln^-N_20_ interfaces, where each model predicts activity for one of 9 SpCas9 variants: SpCas9, SpCas9-VRQR, SpCas9-NG, xCas9, Sniper-Cas9, eSpCas9(1.1), SpCas9-HF1, HypaCas9 and evoCas9. DeepSmallCas9 [114] is a set of 17 CNN models accepting matched and mismatched (abbreviated (mis)matched) spacer-target interfaces, where matched/mismatched refers to the 19 nucleotides apart from the 5’ guanine. In DeepSmallCas9, each model predicts activity for one of 17 small Cas9 nucleases/variants: sRGN3.1, SlugCas9, Sa-SlugCas9, SlugCas9-HF, SauriCas9, SauriCas9-KKH, SaCas9, eSaCas9, ef-SaCas9, SaCas9-HF, SaCas9-KKH, SaCas9-KKH-HF, St1Cas9, CjCas9, enCjCas9, Nm1Cas9 and Nm2Cas9. DeepSpCas9-v2 [114] is similar to DeepSmallCas9, but predicts (mis)matched G/gN_19_ SpCas9 activity.

DeepCas9variants [143] is a collection of 9 CNN models accepting matched G/gN_19_ interfaces, where each model predicts activity for one of 9 SpCas9/ScCas9 variants: SpCas9, SpCas9-VRQR, SpCas9-NG, SpCas9-NRRH, SpCas9-NRTH, SpCas9-NRTH, SpG, SpRY and Sc++. DeepSniper [29] is a collection of 4 tools which we name DS Sniper1 on, DS Sniper2L on, DS Sniper1 off and DS Sniper1 off. Specifically, DS Sniper1 on and DS Sniper2L on predict Sniper-Cas9 and Sniper2L activity for matched G/gN_19_ and tRNA^Gln^-N_20_ interfaces, respectively, and DS Sniper1 off and DS Sniper2L off predict Sniper-Cas9 and Sniper2L activity for mismatched G/gN_19_ interfaces, respectively. For ease of notation, we use the abbreviation DS Sniper1 for the joint use of DS Sniper1 on and DS Sniper1 off for predicting Sniper-Cas9 activity. Likewise, we use the abbreviation DS Sniper2L for the joint use of DS Sniper2L on and DS Sniper2L off for predicting Sniper2L activity.

To fairly compare between DeepEmbCas9 models and DeepSniper on datasets containing both matched and mismatched interfaces, we refer to DS Sniper1, which abbreviates for using DS Sniper1 on when the input interface is matched, and using DS Sniper1 off when the input interface is mismatched. Similarly, DS Sniper2L refers to the use of DS Sniper2L on for matched interfaces and DS Sniper2L off for mismatched interfaces.

For notational convenience, we abbreviate DeepSpCas9variants, DeepCas9variants and DeepSniper as DSpCv, DCv and DS, respectively in main and supplementary figures.

#### 2.8.2 In-distribution performance

To enable fair performance comparisons between DeepEmbCas9 and existing published cleavage activity prediction tools for CRISPR-Cas9 variants, we trained DeepEmbCas9 on train-valid-test splits com-patible with such tools. To achieve this, we first obtained test partitions used for evaluating DeepHF, DeepSpCas9, DeepxCas9, DeepSpCas9-NG, DeepSpCas9variants, DeepSmallCas9, DeepCas9variants and DeepSniper from their respective source studies [100, 95, 141, 142, 114, 143, 29]. Following GitHub code provided by Wang et al. [100], we also used scikit-learn’s `train_test_split` [158] with `random_state=40` and a 85%-15% train-test ratio to obtain the test partition data from Wang et al. To avoid data leakage and overrepresentation of specific experimental configurations, experimental configurations which have datapoints in both training and test partitions are relabelled to be part of the training partition. These three steps resulted in two non-overlapping partitions: a non-test partition of size 1582129 with 40 Cas9 variants and 16 gRNA scaffolds, and a test partition of size 164557 with 39 Cas9 variants and 7 gRNA scaffolds. Notably, the curated cross-study test set was a strict subset of the union of the test sets from the 6 source studies. We then randomly split the non-test partition into training and validation partitions using a 80%-20% split.

DeepEmbCas9 was trained on the resulting training partition, with the validation partition used for early stopping, where both partitions used the “Mean background subtracted indel frequency (%)” label column. Once trained, DeepEmbCas9 was then evaluated on the test partition, which used labels from the “Mean background subtracted indel frequency (source, %)”, using Spearman rank correlation and root mean squared error (RMSE) as evaluation metrics. DeepEmbCas9-MVE, DeepEnsEmbCas9 naive and DeepEnsEmbCas9 (see subsection “Uncertainty Quantification via Deep Ensembles”) were evaluated in a similar way.

We used DeepHF, DeepSpCas9, DeepxCas9, DeepSpCas9-NG, DeepSpCas9variants, DeepSmallCas9, DeepCas9variants and DeepSniper as baselines to compare with DeepEmbCas9 and DeepEnsEmbCas9. In short, each tool was evaluated on test sets sharing the same nuclease and guide length as the tool (e.g., DeepSpCas9 was evaluated on all test sets with matched G/gN_19_ SpCas9 interfaces) via Spearman correlation and RMSE. In addition to the GitHub model provided in https://github.com/izhangcd/DeepHF, DeepHF models retrained using the GitHub code provided were also used as baselines.

#### 2.8.3 Leave-one-nuclease-out extrapolation

39 leave-one-nuclease-out train-test splits were formed by excluding test Cas9 nuclease training data from the training partition. We then use the same procedures described above to train the neural network, which we name DeepEmbCas9 omit to distinguish from DeepEmbCas9. Existing tools sharing the same spacer length as the test nuclease yet not trained on test nuclease data are used as baselines for DeepEmbCas9 omit. For example, DeepSniper is one of the baselines used for the DeepEmbCas9 omit variant whichI believe that this should be in the previous section, e.g. be section 2.8.4, a similar section to 2.8.3. excluded wild-type SpCas9 training data.

### 2.9 Model interpretation

We interpret DeepEmbCas9 using SHapley Additive exPlanations (SHAP) [159], an additive feature attribution method. Namely, SHAP values were approximated using DeepExplainer and a background dataset consisting of 100 randomly sampled datapoints. Computed SHAP values were then used for deriving SHAP importances for individual input features, where the SHAP importance of the *j*th input feature given by 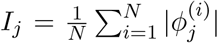 for dataset size *N* and SHAP value 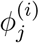 attributed to the *i*th datapoint-*j*th feature pair. Leveraging SHAP’s additivity property, we also computed SHAP importances for set of input features (i.e., feature group) using the formula 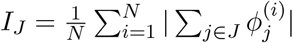, where *J* denotes the set of input features.

To systematically analyze the different parts of the CRISPR-Cas9 complex, we computed SHAP importances for CRISPR-Cas9 complex components. For fine resolution of the components we considered the following 28 feature groups, namely `spacer + spacer MFE + spacer GCcount, upstream + protospacer + protospacer_Tm, PAM + downstream + PAM downstream_Tm, tRNA preprocessing, Day, spacer_scaffold MFE` and 22 Cas9 feature groups — one for each Cas9 region (see Table S7 for a list of features and feature counts for each feature group). For SHAP importance of Cas9 complex components in coarse resolution we considered the following 6 feature groups:

- spacer one-hot encoding, spacer MFE, 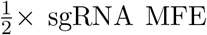, spacer GC count and spacer region part of the sgRNA rLM embedding (labelled as “spacer”);
- target context sequence one-hot encoding, DNA melting temperatures features and protospacer GC count (labelled as non-target strand “NTS”);
- Cas9 pLM embedding features for all Cas9 regions (labelled as “Cas9”);
- rLM embedding features for all regions excluding the spacer and 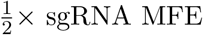 (labelled as “gRNA”); and
- tRNA^Gln^ preprocessing (labelled as “tRNA preprocessing”),

where 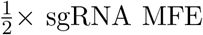 denotes the fact that only half of sgRNA MFE’s SHAP value is used in the SHAP importance calculation (Table S8). As for SHAP importance analysis for each component, in addition to importances for each Cas9 region and sgRNA region, we computed the SHAP importance of each Cas9 domain, i.e.,

- RuvC, by grouping features from RuvC-I, RuvC-II and RuvC-III;
- REC1, by grouping features from REC1-A and REC1-B;
- Linker, by grouping features from the linker loop, L1 and L2;
- PLL-WED-PI, by grouping features from the PLL, WED and PI; and
- NLS-FLAG-P2A, by grouping features from NLS, FLAG, P2A and Other,

in addition to feature groups for BH, REC insert, REC2 and HNH. Since the spacer-target interface plays a primary role in Cas9 cleavage activity prediction, we use heatmaps to visualize the SHAP importance of guide-target positions and nucleotides.

We also devised a framework for assessing the influence of input features on cleavage activity change given specific Cas9 mutation(s). Specifically, given a base Cas9 nuclease *p*_1_ and Cas9 nuclease *p*_2_, with *p*_2_ typically obtained by introducing residue mutations to *p*_1_, we can attribute the difference 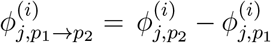 to the *i*th datapoint-*j*th feature pair due to SHAP value’s additivity property. Note that summing the differences over all features yields the activity change. Based on this, we can calculate the *j*th feature’s SHAP importance in predicting activity change, given by 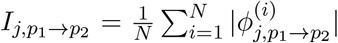 SHAP importances can be extended to groups via the formula 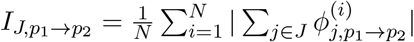 for feature group *J*. We considered the following 39 pairs of Cas9 nucleases in the framework, where A *>* B denotes the mutation from nuclease A to nuclease B: SpCas9 *>* eSpCas9(1.1), SpCas9 *>* SpCas9-HF1, SpCas9 *>* evoCas9, SpCas9 *>* HypaCas9, SpCas9 *>* Sniper-Cas9, SpCas9 *>* xCas9, SpCas9 *>* VRQR-HF1, SpCas9 *>* QQR1, SpCas9 *>* SpCas9-NG, SpCas9 *>* VQR, SpCas9 *>* VRER, SpCas9 *>* VRQR, SpCas9 *>* SpCas9-NRCH, SpCas9 *>* SpCas9-NRRH, SpCas9 *>* SpCas9-NRTH, SpCas9 *>* SpG, SpCas9 *>* SpRY, Sniper-Cas9 *>* Sniper2P, Sniper-Cas9 *>* Sniper2L, VQR *>* VRQR, VQR *>* VRER, VRQR *>* SpG, SpG *>* SpRY, VRQR *>* SpRY, VRQR *>* VRQR-HF1, SpCas9-HF1 *>* VRQR-HF1, NLS-SaCas9 *>* NLS-eSaCas9, NLS-SaCas9 *>* NLS-efSaCas9, NLS-SaCas9 *>* NLS-SaCas9-HF, NLS-SlugCas9 *>* NLS-SlugCas9-HF, NLS-CjCas9 *>* NLS-enCjCas9, NLS-SaCas9 *>* NLS-SaCas9-KKH, NLS-SaCas9 *>* NLS-Sa-SlugCas9, NLS-SauriCas9 *>* NLS-SauriCas9-KKH, NLS-SaCas9 *>* NLS-SaCas9-KKH-HF, NLS-SaCas9-HF *>* NLS-SaCas9-KKH-HF, NLS-SaCas9-KKH *>* NLS-SaCas9-KKH-HF, NLS-SlugCas9 *>* NLS-Sa-SlugCas9, and NLS-SlugCas9 *>* NLS-sRGN3.1.

### 2.10 Uncertainty quantification

We augmented DeepEmbCas9 with uncertainty estimates by using mean-variance estimation [160] and/or deep ensembles [161]. Specifically, we built:

- DeepEmbCas9-MVE by training a single heteroscedastic DeepEmbCas9 model with separate mean and variance-predicting stems (Figure S5) and initialized with different seeds using the Gaussian NLL loss objective. We apply softplus to the variance-predicting stem’s output to ensure non-negativity of the predicted variance. We also add 1 *×* 10^*−*6^ to the variance-predicting stem’s output and clip the global norm of mini-batch gradients to ≤ 5 to maintain numerical stability during training.
- DeepEnsEmbCas9 naive by training 20 (homoscedastic) DeepEmbCas9 models initialized with different seeds using the MSE loss objective; and
- DeepEnsEmbCas9 by training 20 DeepEmbCas9-MVE models.

For DeepEmbCas9-MVE, the predicted mean and variance are given directly by its output heads. For DeepEnsEmbCas9_naive, given point predictions 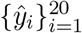 for input **x**_*i*_, the predicted mean and variance is given by 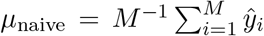 and 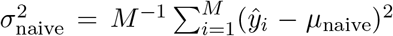, respectively. For DeepEnsEmbCas9, given mean-variance predictions 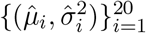 for input **x**_*i*_, the predicted mean and variance is given by 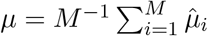 and 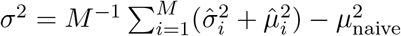, respectively.

### 2.11 Quantile calibration

We generated quantile calibration plots and calculated quantile calibration errors to assess whether the predicted uncertaintes were quantile calibrated. Since quantile calibration was assessed for each test set in the benchmark comparisons, we are technically assessing for group-conditioned quantile calibration, where groups are defined by a specific experimental configuration. To achieve the above, we adopted Kuleshov et al.’s definition of quantile calibration in the regression setting [162], namely that a ML model generating a predictive distribution for datapoint *i* with label *y*_*i*_ and cumulative distribution function (CDF) *F*_*i*_: R → [0, 1] (i.e., quantile function 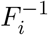) is quantile calibrated if

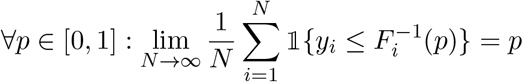

with *N* as the dataset size. We estimate this by selecting confidence levels *p*_*j*_ ∈ *{*0, 0.01, …, 1*}* and plotting 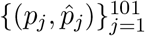 where 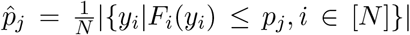 is the empirical frequency. We also adopted Kuleshov et al.’s definition to confidence intervals (CI), i.e., a ML model is calibrated if

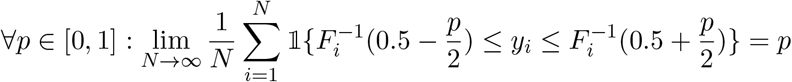

Similarly, we estimate this by selecting confidence intervals *p*_*j*_ = 0, 0.01, …, 0.99, 1 and plotting 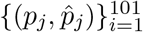 where 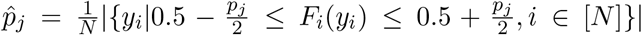 is the CI-based empirical frequency.

We also follow Kuleshov et al.’s approach for computing the quantile calibration error, given by

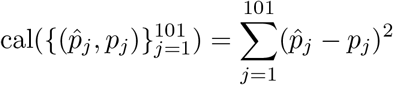

for the two calibration definitions defined above.

## 3 Results

### 3.1 Ranking of pLM-rLM embedding combinations

To determine the best pLM-rLM combination for DeepEmbCas9, we assessed five-fold cross validation Spearman correlations for the 30 pLM-rLM embedding combinations (Table S5). ESM-C-600M with BEACON-B yielded the highest average Spearman correlation of 0.903 *±* 0.002. Averaging pLM-rLM performances for each of the 6 pLM embeddings, we see that ESM-C embeddings ranked the best with ~0.9 Spearman correlation, followed by ProtT5 and Ankh-large embeddings with ~0.89 Spearman correlation. Combinations with gLM2 650M and ESM3 embeddings yielded ~0.8 Spearman correlation. Averaging pLM-rLM performances for each of the 5 rLM embeddings, we see that the rLM embeddings perform similarly at 0.86 *−* 0.88 Spearman correlation, with RNA-FM ranked highest at 0.877 *±* 0.032.

### 3.2 In-distribution performance comparisons

DeepEnsEmbCas9 naive attains higher Spearman correlation than all individual activity prediction tools on 18 out of 51 benchmark test sets (Figure 4, black bars), including 4 mismatched G/gN_19_ interfaces for SpCas9, Sniper-Cas9 and Sniper2L (Figures 4B and 4G), 10 small Cas9 test sets (2 SlugCas9 variants, 2 SauriCas9 variants, 5 SaCas9 increased-fidelity variants and enCjCas9) from Seo et al. [114] (Figure 4I), and 4 other test sets with matched SpCas9, xCas9 and SpCas9-NG interfaces (Figures 4A, C and H). As for the remaining 33 test sets, DeepEnsEmbCas9 naive has an average Spearman performance drop of 3.43 *×* 10^*−*2^ compared to the best-performing individual activity prediction tools, with the test set containing matched A/GN_19_ SpCas9-HF1 interfaces from Wang et al. [100] yielding the largest Spearman drop of 0.137 (DeepHF SpCas9-HF1’s 0.881 vs. DeepEnsEmbCas9 naive’s 0.744). Among test sets which are outside of the 51 benchmark test sets and lack baselines, DeepEnsEmbCas9 naive attains 0.440-0.922 Spearman correlation in 9 out of 10 test sets (Figures S10 rows 2-3, S11, S15, S21 row 3, S22 row 3 and S23 row 3).

**Figure 4.**
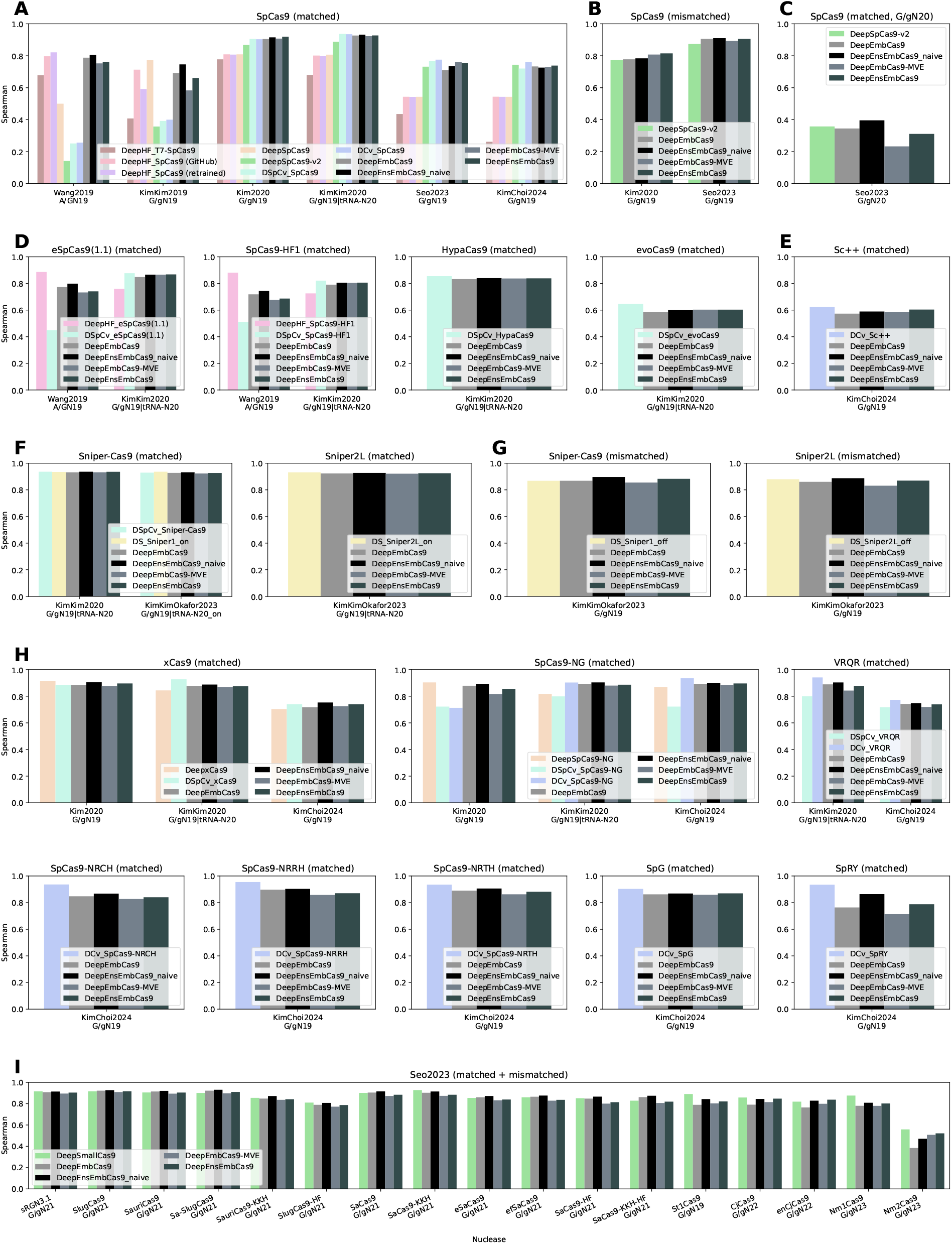
Benchmark test Spearman correlation comparison for DeepEmbCas9, DeepEnsEmb-Cas9 naive, DeepEmbCas9-MVE and DeepEnsEmbCas9 against DeepHF, DeepSpCas9, Deepx-Cas9, DeepSpCas9-NG, DeepSpCas9variants, DeepSmallCas9, DeepSpCas9-v2, DeepCas9variants and DeepSniper across 39 Cas9 nucleases. The test sets consist of (A) matched AN_19_, G/gN_19_ and tRNA^Gln^-N_20_ wild type SpCas9 interfaces; (B) mismatched G/gN_19_ wild type SpCas9 interfaces; (C) matched G/gN_20_ wild type SpCas9 interfaces; (D,E,F) matched AN_19_, G/gN_19_ and tRNA^Gln^-N_20_ interfaces for 4 increased-fidelity SpCas9 variants (D), Sc++ (E), and 2 Sniper variants (F); (G) mismatched G/gN_19_ interfaces for 2 Sniper variants; (H) matched G/gN_19_ and tRNA^Gln^-N_20_ interfaces for 8 PAM-altered SpCas9 variants; and (I) matched and mismatched interfaces for 17 wild type or engineered small Cas9 nucleases.

Analogous test performance comparisons for DeepEmbCas9, DeepEnsEmbCas9 and DeepEmbCas9-MVE can be found in Supplementary Information section “In-distribution performance” subsection “In-distribution performance comparisons for DeepEmbCas9, DeepEnsEmbCas9 and DeepEmbCas9-MVE”. Detailed Spearman correlation comparisons between DeepEmbCas9 and individual activity prediction tools for each benchmark test set is provided in Supplementary Information section “In-distribution performance” subsection “Detailed analysis of DeepEmbCas9’s in-distribution performance”.

### 3.3 Impact of deep ensembles on in-distribution performance

When considering averaged Spearman correlations across the 51 benchmark test sets, DeepEnsEmb-Cas9 naive (0.834) attains slightly higher Spearman correlation compared to DeepEmbCas9 (0.814). Likewise, DeepEnsEmbCas9 (0.817) attains slightly higher Spearman correlation compared to DeepEmbCas9-MVE (0.800). In sum, comparing among the 4 DeepEmbCas9 models with and without mean variance estimation and/or ensembling, DeepEnsEmbCas9 naive, DeepEnsEmbCas9, DeepEmbCas9-MVE attain the highest Spearman correlation in 39, 11 and 1 benchmark test set(s) with baselines out of the 51 in total, respectively (Figure 4). In particular, we observed the ranking DeepEnsEmbCas9 naive *>* DeepEmbCas9 *>* DeepEnsEmbCas9 *>* DeepEmbCas9-MVE for test sets with matched SpCas9 interfaces from Wang et al. [100] (Figures 4A and S7 row 1) and Kim, Kim et al. [95] (Figures 4A and S8), mismatched G/gN_19_ and matched GN_20_ SpCas9 interfaces from Seo et al. [114] (Figures 4B-C and S16 rows 2 and 4), matched eSpCas9(1.1) and SpCas9-HF1 interfaces from Wang et al. [100] (Figures 4D and S7 rows 2 and 3), matched Sniper-Cas9 interfaces from Kim, Kim, Okafor et al. [29] (Figures 4F and S21 row 1), matched SpCas9-NG interfaces from Kim et al. [141] (Figures 4H and S9 row 3), matched xCas9, SpCas9-NG and VRQR interfaces from Kim, Kim et al. [142] (Figures 4H and S14), matched VRQR, SpCas9-NRCH, SpCas9-NRRH and SpCas9-NRTH from Kim, Choi et al. [143] (Figures 4H, S18 row 3, and S19), and (mis)matched G/gN21 interfaces for SaCas9 variants, sRGN3.1, SlugCas9 variants and SauriCas9 variants from Seo et al. [114] (Figures 4I and S24 and S25). In addition, we observed the ranking DeepEnsEmbCas9 naive *>* DeepEnsEmbCas9 *>* DeepEmbCas9 *>* DeepEmbCas9-MVE in test sets for matched SpCas9 interfaces from Kim, Kim et al. [142] (Figures 4A and S12), matched Sniper2L interfaces from Kim, Kim, Okafor et al. [29] (Figures 4F and S21 row 2), mismatched Sniper-Cas9 and Sniper2L from Kim, Kim, Okafor et al. [29] (Figures 4G and S22 rows 1-2), matched xCas9 interfaces from Kim et al. [141] (Figures 4H and S9 row 2), matched SpCas9-NG and SpRY interfaces from Kim, Choi et al. [143] (Figures 4H, S18 row 2 and S20 row 2) and (mis)matched Nm1Cas9 interfaces from Seo et al. [114] (Figures 4I and S26 row 4). We observed the ranking DeepEnsEmbCas9 naive *>* DeepEnsEmbCas9 *>* DeepEmbCas9-MVE *>* DeepEmbCas9 for matched HypaCas9 and Sniper-Cas9 interfaces from Kim, Kim et al. [142] (Figures 4D, 4F and S13 rows 1 and 3), matched xCas9 interfaces from Kim, Choi et al. [143] (Figures 4H and S18 row 1) and (mis)matched St1Cas9 interfaces from Seo et al. [114] (Figures 4I and S26 row 1).

We saw various rankings among the 11 test sets where DeepEnsEmbCas9 performs best within the four DeepEmbCas9 models. Among the four models, DeepEnsEmbCas9 and DeepEmbCas9 have the highest and lowest Spearman correlations in 9 out of 11 test sets. Among the 9 test sets, DeepEnsEm-bCas9 naive ranked higher than DeepEmbCas9-MVE in 6 test sets (i.e., those with matched SpCas9 interfaces from Kim et al. [141] (Figures 4A and S9 row 1), matched eSpCas9(1.1) and SpCas9-HF1 interfaces from Kim, Kim et al. [142] (Figures 4D and S12 rows 2-3), matched Sc++ interfaces from Kim, Choi et al [143] (Figures 4E and S17 row 2), and (mis)matched CjCas9 and enCjCas9 interfaces from Seo et al. [114] (Figures 4I and S26 rows 2-3)), and DeepEmbCas9-MVE ranked higher than DeepEnsEmbCas9 naive in the other 3 test sets (i.e., those with mismatched SpCas9 interfaces from Seo et al. [114] (Figures 4B and S16 row 2), matched evoCas9 interfaces from Kim, Kim et al. [142] (Figures 4D and S13 row 2) and (mis)matched Nm2Cas9 interfaces from Seo et al. [114] (Figures 4I and S26 row 5)). The four models are ranked DeepEnsEmbCas9 *>* DeepEmbCas9 *>* DeepEmbCas9-MVE *>* DeepEnsEmbCas9 naive on the test set with matched SpCas9 interfaces from Kim, Choi et al. [143] (Figures 4A and S17 row 1), and DeepEnsEmbCas9 *>* DeepEnsEmbCas9 naive *>* DeepEmb-Cas9 *>* DeepEmbCas9-MVE for the test set with matched SpG interfaces from Kim, Choi et al. [143] (Figures 4H and S20 row 1). DeepEmbCas9-MVE ranked best among the four models (DeepEmbCas9-MVE *>* DeepEnsEmbCas9 *>* DeepEnsEmbCas9 naive *>* DeepEmbCas9) in the test set with matched SpCas9 interfaces from Seo et al. [114] (Figures 4A and S16 row 1).

### 3.4 Leave-one-nuclease-out performance comparisons

In the leave-one-nuclease-out extrapolation setting, DeepEnsEmbCas9 naive omit (Figures S7-S26, red bars) attains higher Spearman correlation than all individual activity prediction tools on 17 out of 48 test sets (i.e., the 51 benchmark test sets excluding matched G/gN_20_ SpCas9, St1Cas9 and Nm2Cas9 interfaces). These include 2 mismatched G/gN_19_ SpCas9 interface test sets (Figures S10 row 1 and S16 row 2), 1 matched eSpCas9(1.1) interface test set from Kim, Kim et al. [142] (Figure S12 row 2), 2 matched xCas9 interface test sets from Kim et al. [141] and Kim, Kim et al. [142] (Figures S9 row 2 and S14 row 1), 1 matched SpCas9-NG interface test set from Kim et al. [141] (Figure S9 row 3), 3 matched SpCas9-NRCH, SpCas9-NRRH and SpCas9-NRTH interface test sets from Kim, Choi et al. [143] (Figure S19), 2 mismatched Sniper-Cas9 and Sniper2L interface test sets from Kim, Kim, Okafor et al. [29] (Figure S22 rows 1-2), and 6 small Cas9 test sets (3 SaCas9 variants, SauriCas9-KKH, SlugCas9-HF and Nm1Cas9) from Seo et al. [114] (Figures S24-S26). As for the remaining 34 test sets, DeepEnsEmbCas9 naive omit has an average Spearman performance drop of 5.09 *×* 10^*−*2^ compared to the best-performing individual activity prediction tools not trained on the test sets’ nucleases, with the test set containing matched G/gN_19_ Sc++ interfaces from Kim, Choi et al. [143] yielding the largest Spearman drop of 0.266 (DSpCv Sniper-Cas9’s 0.554 vs. DeepEnsEmbCas9 naive omit’s 0.288; Figure S17 row 2).

Among the test sets which have extrapolation baselines and are not in the 51 benchmark test sets (excluding Kim, Kim et al. [142]’s QQR1 test set), DeepEnsEmbCas9 naive omit outperforms all individual activity prediction tools on 7 out of 14 test sets, namely mismatched SpCas9 and xCas9 interface test sets from Kim et al. [141] (Figure S10 rows 1 and 2), (mis)matched SpCas9, xCas9 and SpCas9-NG interface test sets from Kim et al. [141] (Figure S11), a matched VRER test set from Kim, Kim et al. [141] (Figure S15 row 3), and (mis)matched Sniper2L interface test sets from Kim, Kim, Okafor et al. [29] (Figure S23 row 2). As for the remaining 7 test sets, DeepEnsEmbCas9 naive omit has an average Spearman performance drop of 2.24 *×* 10^*−*2^ compared to the best-performing individual activity prediction tools, with the test set containing matched VQR G/gN_19_ interfaces from Kim, Kim et al. [141] yielding the largest Spearman drop of 5.34 *×* 10^*−*2^ (DCv VRQR’s 0.691 vs. DeepEnsEmbCas9 naive omit 0.637; Figure S20 row 2).

Analogous test performance comparisons for DeepEmbCas9, DeepEnsEmbCas9 and DeepEmbCas9-MVE can be found in Supplementary Information section “Leave-one-nuclease-out extrapolation performance” subsection “Leave-one-nuclease-out extrapolation performance comparisons for DeepEmbCas9, DeepEnsEmbCas9 and DeepEmbCas9-MVE”. Detailed Spearman correlation comparisons between DeepEmbCas9 and individual activity prediction tools for each benchmark test set is provided in Supplementary Information section “Leave-one-nuclease-out extrapolation performance” subsection “DeepEmbCas9 extrapolates to unseen Cas9 variants”.

### 3.5 Impact of deep ensembles on leave-one-nuclease-out performance

When considering averaged Spearman correlations across the 51 benchmark test sets in the leave-one-nuclease-out extrapolation setting, DeepEnsEmbCas9 naive (0.786) attains slightly higher Spearman correlation compared to DeepEmbCas9 (0.760). Likewise, DeepEnsEmbCas9 (0.774) attains slightly higher Spearman correlation compared to DeepEmbCas9-MVE (0.756). In sum, comparing among the 4 DeepEmbCas9 models with and without mean variance estimation and/or ensembling, Deep-EnsEmbCas9 naive, DeepEnsEmbCas9, DeepEmbCas9-MVE and DeepEmbCas9 attain the highest Spearman correlation in 40, 15, 7 and 1 benchmark test set(s) with(out) baselines out of the 63 in total, respectively (Figures S7-S26). Specifically, we observed the ranking DeepEnsEmbCas9 naïve DeepEnsEmbCas9 *>* DeepEmbCas9-MVE *>* DeepEmbCas9 for test sets with matched SpCas9-HF1 and eSpCas9(1.1) interfaces from Wang et al. [100], matched SpCas9 interfaces from Kim, Kim et al. [95], matched SpCas9-NG interfaces from Kim et al. [141], matched VQR, VRER and VRQR-HF1 interfaces from Kim, Kim et al. [142], matched SpCas9-NRCH, SpCas9-NRRH, SpCas9-NRTH and SpRY interface from Kim, Choi et al. [143], matched Sniper-Cas9 and Sniper2P interfaces from Kim, Kim, Okafor et al. [143], mismatched Sniper-Cas9 and Sniper2L interfaces from Kim, Kim, Okafor et al. [143], (mis)matched Sniper2P interfaces from Kim, Kim, Okafor et al. [143], and (mis)matched SaCas9, eSaCas9, efSaCas9, SaCas9-HF, SaCas9-KKH, SaCas9-KKH-HF, Sa-SlugCas9 and SlugCas9-HF interfaces from Seo et al. [114]. In addition, we observed the ranking DeepEnsEm-bCas9 naive *>* DeepEnsEmbCas9 *>* DeepEmbCas9 *>* DeepEmbCas9-MVE for test datasets with matched xCas9 interfaces from Kim et al. [141], matched VRQR and QQR1 interfaces from Kim, Kim et al. [142], matched Sniper2L interfaces from Kim, Kim, Okafor et al. [29], (mis)matched Sniper-Cas9 and Sniper2L interfaces from Kim, Kim, Okafor et al. [29], and (mis)matched SlugCas9 interfaces from Seo et al. [114]. We observed DeepEnsEmbCas9 naive *>* DeepEnsEmbCas9 *>* DeepEmbCas9-MVE DeepEmbCas9 for test sets with matched SpCas9 and Sniper-Cas9 interfaces from Kim, Kim et al. [142], mismatched SpCas9 interfaces from Seo et al. [114], matched VRQR interfaces from Kim, Choi et al. [143], and (mis)matched CjCas9 and enCjCas9 interfaces from Seo et al. [114].

We observed various rankings among the 15 test sets where DeepEnsEmbCas9 performs best within the four DeepEmbCas9 models. DeepEmbCas9 has the lowest Spearman correlation among the four models in 12 out of 15 test sets. Among the 12 test sets, DeepEmbCas9-MVE rank higher than DeepEnsEmbCas9 naive for the following 9 test sets (i.e., those with mismatched SpCas9 and xCas9 interfaces from Kim et al. [141], (mis)matched SpCas9 interfaces from Kim et al. [141], matched SpCas9-HF1 interfaces from Kim, Kim et al. [142], matched SpCas9, Sc++ and SpG interfaces from Kim, Choi et al. [143], and matched St1Cas9 and Nm2Cas9 interfaces from Seo et al. [114]), and DeepEnsEmbCas9 naive ranks higher than DeepEmbCas9-MVE for the other 3 test sets (i.e., those with matched SpCas9 interfaces from Kim et al. [141], (mis)matched xCas9 interfaces from Kim et al. [141], and matched eSpCas9(1.1) interfaces from Kim, Kim et al. [142]).

We observed various rankings among the 7 test sets where DeepEmbCas9-MVE performs best within the four DeepEmbCas9 models. DeepEmbCas9 has the lowest Spearman correlations among the four models in 6 out of the 7 test sets. Among the 6 test sets, DeepEnsEmbCas9 ranked higher than DeepEnsEmbCas9 naive on 3 test sets (i.e., those with matched evoCas9 interfaces from Kim, Kim et al. [142], matched SpCas9-NG interfaces from Kim, Choi et al. [143], and (mis)matched Nm1Cas9 interfaces from Seo et al. [114]), and DeepEnsEmbCas9 naive ranked higher than DeepEnsEmbCas9 on the other 3 test sets (i.e., those with matched GN_20_ interfaces from Seo et al. [114], matched xCas9 interfaces from Kim, Choi et al. [143], and (mis)matched SauriCas9 interfaces from Seo et al. [114]).

### 3.6 Uncertainty estimation via mean-variance estimation

In the in-distribution setting on the 51 benchmark test sets, DeepEnsEmbCas9 (Figures 5 and S31) and DeepEmbCas9-MVE (Figures S32 and S33) are more quantile calibrated than DeepEnsEmbCas9 naive (Figures S34 and S35). The left-tailed quantile calibration error of DeepEnsEmbCas9 naive is higher than that of DeepEmbCas9-MVE and DeepEnsEmbCas9 in all 51 benchmark test sets except for those with matched G/gN_19_ SpCas9 interfaces from Seo et al. [114] and matched xCas9 interfaces from Kim et al. [141] (Figure S36). As for CI-based quantile calibration, DeepEnsEmbCas9 naive has higher error than DeepEmbCas9-MVE and DeepEnsEmbCas9 in all 51 benchmark test sets except for those with matched G/gN_19_ and G/gN_20_ SpCas9 interfaces from Seo et al. [114] (Figure S37).

**Figure 5.**
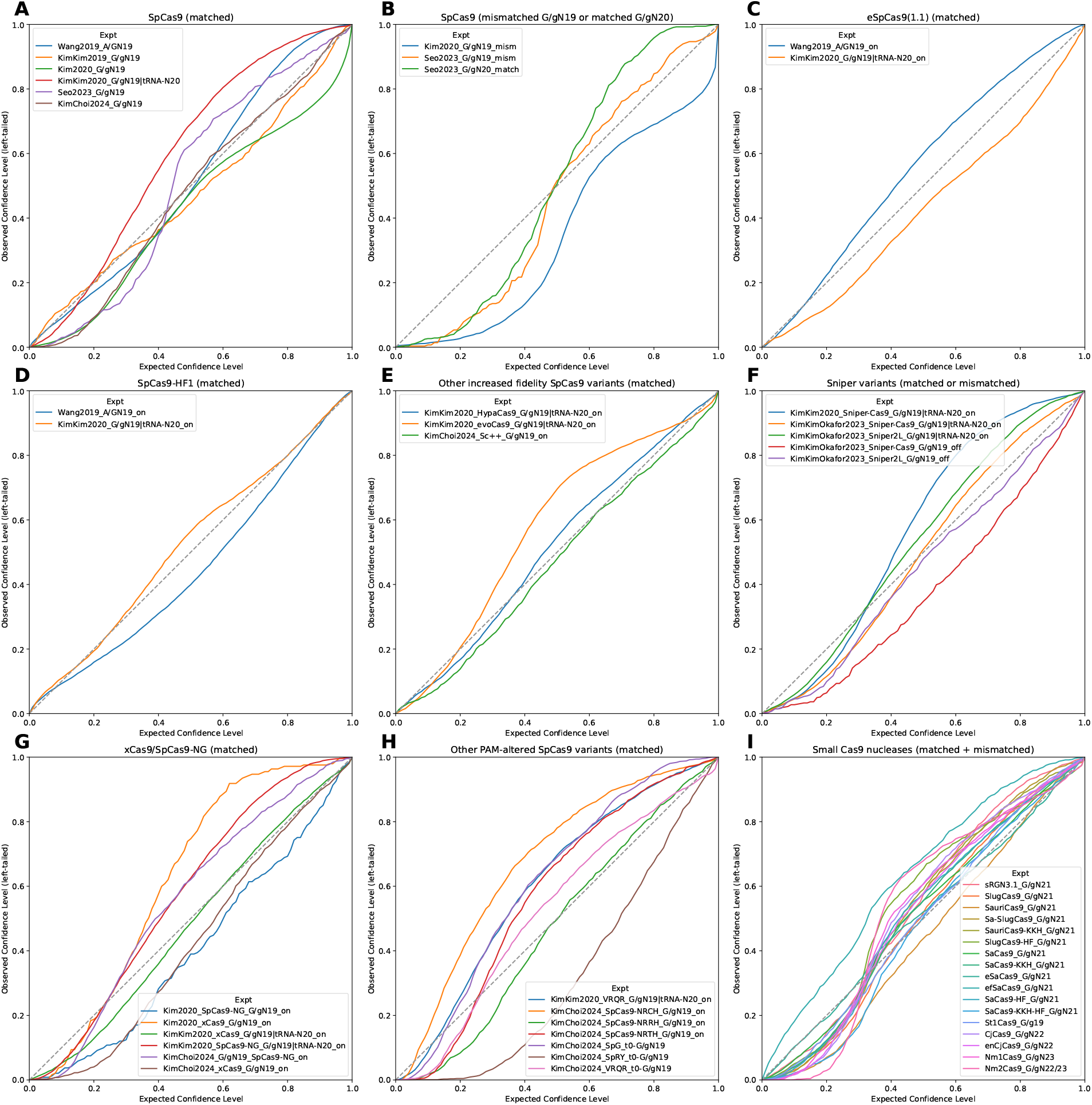
Quantile calibration plots for DeepEnsEmbCas9, conditioned on (A) matched AN_19_, G/gN_19_ and tRNA^Gln^-N_20_ wild type SpCas9 interfaces; (B) mismatched G/gN_19_ and matched G/gN_20_ wild type SpCas9 interfaces; (C) matched AN_19_, G/gN_19_ and tRNA^Gln^-N_20_ eSpCas9(1.1) interfaces; (D) matched AN_19_, G/gN_19_ and tRNA^Gln^-N_20_ SpCas9-HF1 interfaces; for matched G/gN_19_ and tRNA^Gln^-N_20_ HypaCas9/evoCas9 and G/gN_19_ Sc++ interfaces; (F) matched G/gN_19_ and tRNA^Gln^-N_20_ and mismatched G/gN_19_ interfaces for 2 Sniper variants; (G,H) matched G/gN_19_ and tRNA^Gln^-N_20_ interfaces for xCas9/SpCas9-NG (G) and 6 other PAM-altered SpCas9 variants (H); and (I) matched and mismatched interfaces for 17 wild type or engineered small Cas9 nucleases.

In the leave-one-nuclease-out extrapolation setting on the 51 benchmark test sets, DeepEnsEmbCas9 omit (Figures S38 and S39) and DeepEmbCas9-MVE omit (Figures S40 and S41) generally have worse quantile calibration compared to their in-distribution counterparts. Like DeepEnsEmbCas9 naive, DeepEnsEmbCas9 naive omit is not quantile calibrated (Figures S42 and S43). DeepEnsEm-bCas9 naive omit has higher left-tailed quantile calibration error than DeepEnsEmbCas9 omit and DeepEmbCas9-MVE omit for all benchmark test sets except for those with mismatched SpCas9 interfaces from Seo et al. [114], matched SpCas9-NG interfaces from Kim et al. [142], matched SpCas9-NG and SpG interfaces from Kim, Choi et al. [143], and (mis)matched Nm1Cas9 and Nm2Cas9 interfaces from Seo et al. [114] (Figure S44). As for CI-based quantile calibration, DeepEnsEmbCas9 naive omit has higher left-tailed quantile calibration error than DeepEnsEmbCas9 omit and DeepEmbCas9-MVE omit for all benchmark test sets except for those with matched GN_20_ SpCas9 interfaces from Seo et al. [114], matched SpCas9-NG interfaces from Kim et al. [141], and (mis)matched St1Cas9 and Nm2Cas9 interfaces from Seo et al. [114] (Figure S45). Comparing between source studies, we see that DeepEmbCas9-MVE and DeepEnsEmbCas9 have higher left-tailed and CI-based quantile calibration for test sets from Wang et al. [100], Kim, Kim et al. [95] and Kim et al. [141] than in the other 4 studies.

### 3.7 SHAP importance analysis reveals PAM and Cas9 driving DeepEmbCas9 predictions

SHAP importance analysis on the benchmark test sets using different sets of feature groups reveal pertinent feature groups influencing DeepEmbCas9’s predicted Cas9 cleavage activity. When calculating SHAP importance of CRISPR-Cas9 complex components in fine resolution, the top three feature groups with the highest SHAP importance are “PAM + downstream + PAM downstream Tm”, “spacer + spacer MFE + spacer GCcount” and “upstream + protospacer + protospacer Tm” (Figure 6A). When calculating SHAP importance of CRISPR-Cas9 complex componets in coarse resolution, the top three feature groups with the highest SHAP importance are the NTS, Cas9 and the spacer (Figure 6B). As for Cas9 domains, PLL-WED-PI, Linker and REC insert have the highest SHAP importance (Figure 6C). Using Cas9 regions as feature groups, REC insert, L2 and REC1-B have the highest SHAP importance (Figure 6D). sgRNA regions are ranked spacer *>* repeat-anti-repeat *>* polyT *>* trcrRNA rest by SHAP importance (Figure 6E). Positions −2 and −3, specifically −2G and −3G, on the target strand have the highest SHAP importance among PAM and downstream TS nucleotides (Figures 6G and 6I). +1G on the spacer is the most important nucleotide among the upstream and heteroduplex nucleotides, with target positions +22 to +17 and spacer/target positions +3 to +1 being important (Figures 6F and 6H). Importance at positions +1 to +7 is consistent with the seed region present in all 40 Cas9 variants studied (Figure S49-S54).

**Figure 6.**
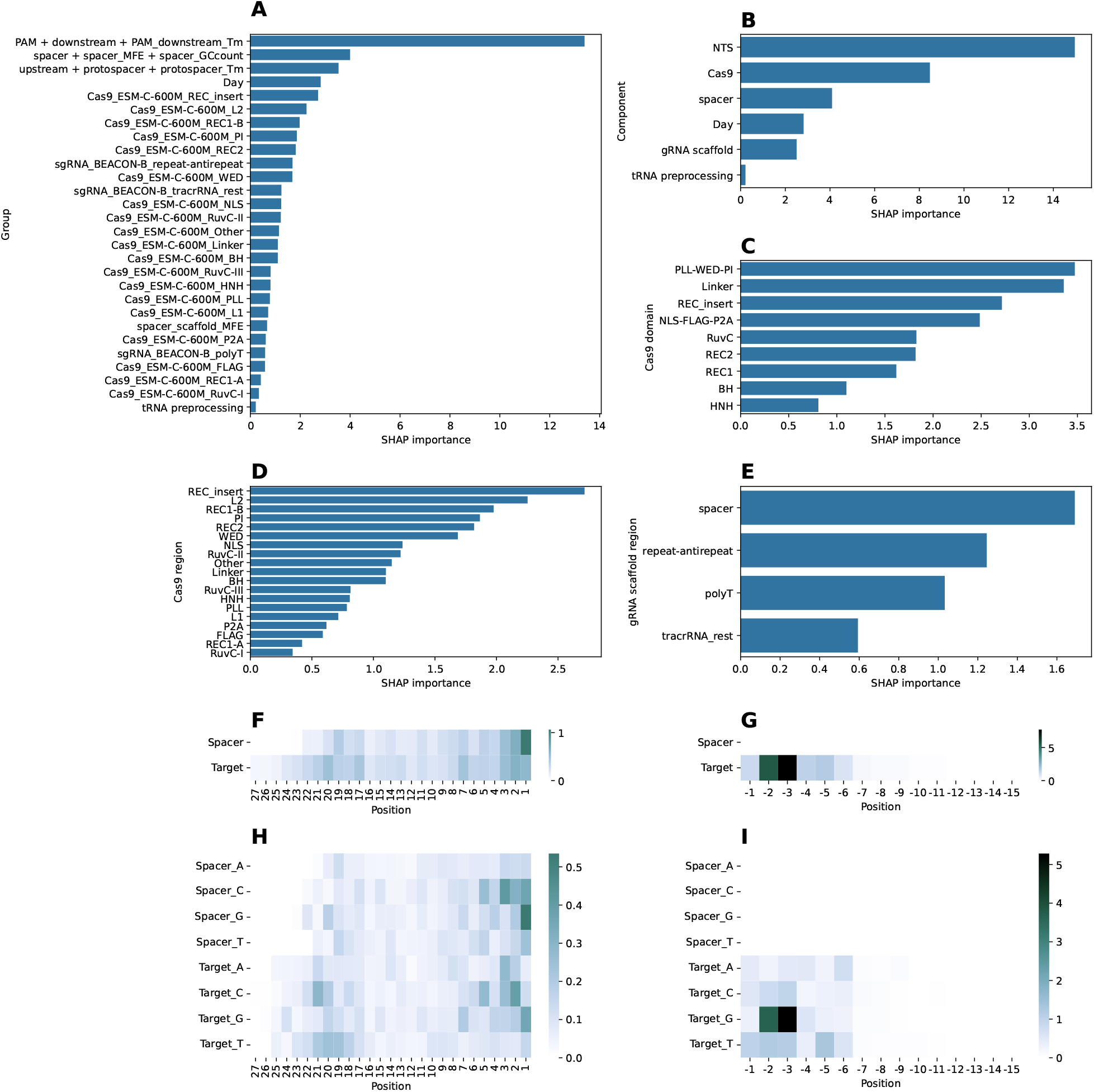
SHAP importance analysis of input features in DeepEmbCas9 (ESM-C-600M-BEACON-B combination) on benchmark test sets. The analysis consists of SHAP importance of Cas9 complex components in fine (A) and coarse (B) resolutions; (coarse-grained) Cas9 domains (C) and (fine-grained) Cas9 regions (D) as defined in Figure 2C; sgRNA regions (E) as defined in Figure 2A; spacer-target nucleotide positions in the upstream-heteroduplex (F) and PAM-downstream regions (G); and spacer-target one-hot encoding features in the upstream-heteroduplex (H) and PAM-downstream regions (I).

### 3.8 DeepEmbCas9’s predicted activity change from Cas9 mutations reflected in Cas9 domain/region SHAP importances

Using the framework for assessing the influence of input features on cleavage activity change given specific Cas9 mutation(s) or domain substitutions, we see that SHAP importances vary only for Cas9 and NTS (Figure 7A). Interestingly, Cas9 SHAP importance positively correlates with NTS SHAP importance (0.593 Spearman correlation; Figure 7B).

**Figure 7.**
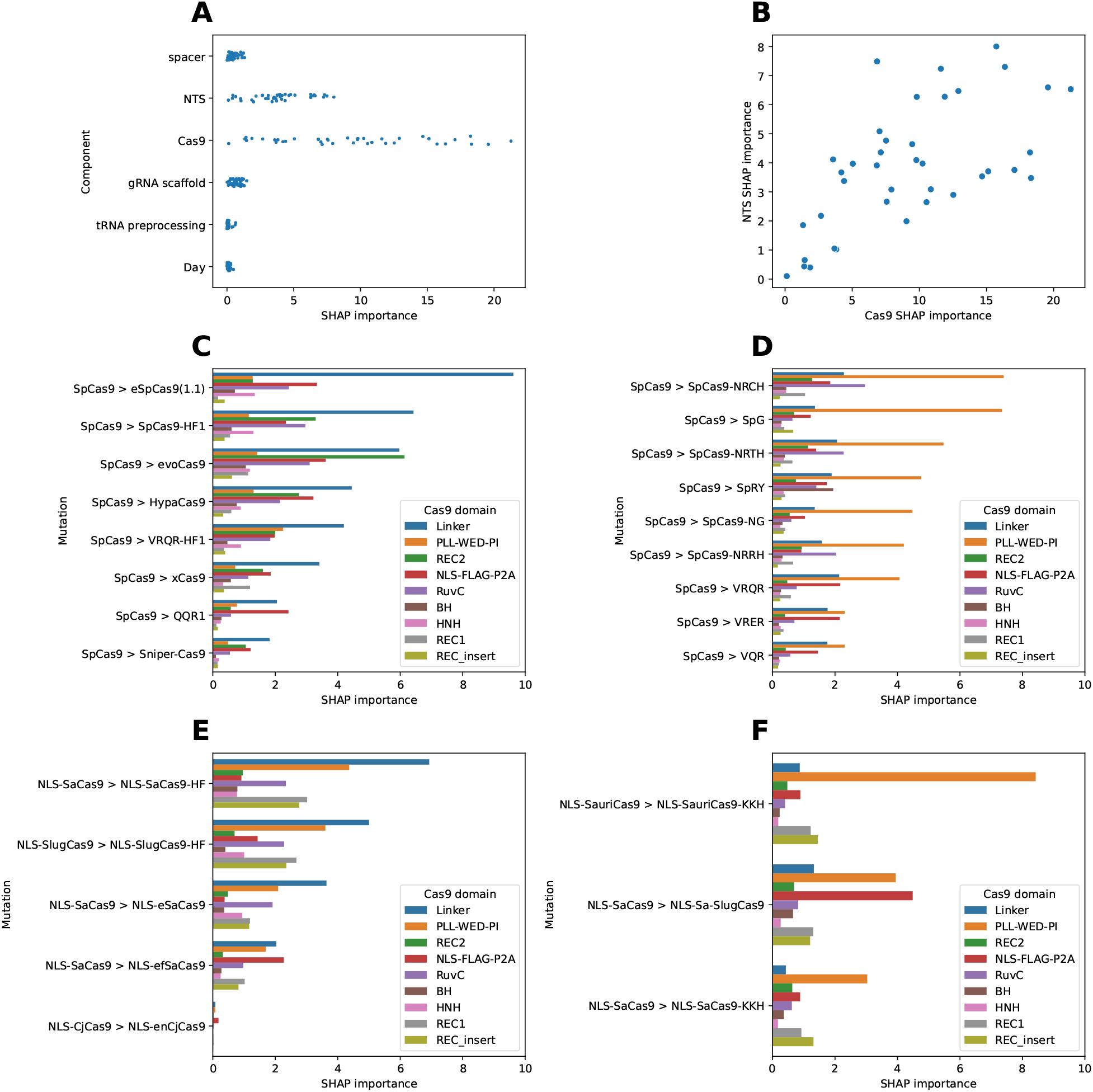
CRISPR-Cas9 complex components and Cas9 domains driving DeepEmbCas9’s change in predicted activity when introducing residue mutations in Cas9. (A) SHAP importance of CRISPR-Cas9 complex components in the 39 nuclease pairs considered. (B) Cas9 vs. NTS SHAP importance in the 39 nuclease pairs considered. (C,D) Cas9 domain importances for SpCas9 variants without (C) and with (D) D1135L/V/N mutations. (E,F) Cas9 domain importances for small Cas9 variants with (E) increased fidelity and (F) PAM-altering variants. Names of Cas9 variants are abbreviated by removing the suffix “-NLS-FLAG-P2A”.

When assessing Cas9 domain importances, the Linker Cas9 domain (specifically L2; Figure S47A) dominates for D1135L/V/N mutation-lacking SpCas9 variants (Figure 7C), all of which are increased fidelity variants apart from xCas9 and QQR1. The phenomenon is most pronounced for nuclease pairs SpCas9 *>* eSpCas9(1.1) and SpCas9 *>* SpCas9-HF1. In contrast, the PLL-WED-PI domain dominates for D1135 mutation-containing SpCas9 variants (Figure 7C), all of which are PAM-altered variants. Specifically, WED among the PLL, WED and PI regions (Figure S47B). Likewise, L2 in Linker dominates in small Cas9 mutations resulting in increased fidelity (Figures 7E and S47C), and PLL in PLL-WED-PI has significant importance in PAM-altering small Cas9 mutations (Figure 7F and Figure S47D).

Other nuclease pairs show similar SHAP importance patterns. Low Cas9 region/domain SHAP im-portances were recorded for nuclease pairs Sniper-Cas9 *>* Sniper2P, Sniper-Cas9 *>* Sniper2L, VQR VRER and VQR *>* VRQR (Figures S46A-B and S48A-B). WED in PLL-WED-PI was found to be important in VRQR *>* SpG, VRQR *>* SpRY and SpCas9-HF1 *>* VRQR-HF1 (Figures S46C-D and S48C-D). Both the bridge helix (BH) from Linker and WED in PLL-WED-PI were important in SpG *>* SpRY(Figures S46C and S48C). In VRQR *>* VRQR-HF1, L2 in Linker and REC2 were found to be important (Figures S46D and S48D). nuclease pairs SaCas9-KKH *>* SaCas9-KKH-HF and SaCas9 *>* SaCas9-KKH-HF showed high SHAP importance for L2 in Linker and PLL-WED-PI(Figures S46E and S48E). PLL in PLL-WED-PI was important in SaCas9-HF *>* SaCas9-KKH-HF, Linker was important in SlugCas9 *>* sRGN3.1, and L2 in Linker and NLS-FLAG-P2A were important in SlugCas9 *>* Sa-SlugCas9 (Figures S46E-F and S48E-F).

## 4 Discussion

### 4.1 Ranking of pLM-rLM embedding combinations

When varying pLM embeddings used in DeepEmbCas9, we saw pLM-rLM combinations with ESM-C embeddings attain higher Spearman correlations compared to combinations with ProtT5, Ankh-large, gLM2 650M or ESM3 (Table S5). This is possibly a result of ESM-C pLMs being trained on datasets with a significant portion of metagenomic sequences, which would include Cas9 sequences [133]. With regards to the impact of ESM-C model size on performance, we observed higher Spearman correlations observed for ESM-C-600M combinations compared to ESM-C-6B combinations, which is consistent with previous claims saying that medium-sized pLMs already perform well on downstream tasks [163]. The poor performance of ESM3-containing combinations likely reflects the fact that ESM3 [129] — a multimodal generative language model — was trained for controllable protein sequence generation rather than for producing high-quality sequence embeddings. Despite gLM2 650M [153] being a mixed-modality gLM with residue-level sequence representation for protein-coding genes, gLM2 650M combinations did not outperform ESM-C, ProtT5 and Ankh-large combinations, perhaps indicating weaker protein coevolutionary signals in gLM2 650M than in ESM-C models. As for RNA embeddings, evo-1-8k combinations performs comparably to RiNALMo combinations, perhaps indicating weaker RNA coevolution signals in evo-1-8k compared to RNA-FM and BEACON-B.

### 4.2 In-distribution and leave-one-nuclease-out performance

We developed DeepEnsEmbCas9 naive, an ensemble model which predicts CRISPR-Cas9 on/off-target cleavage activity prediction for 40 wild-type/increased-fidelity/PAM-altered SpCas9/small Cas9 variants. In the in-distribution setting, DeepEnsEmbCas9 naive attained comparable Spearman performance to individual activity prediction tools on 38 nucleases across 51 benchmark test sets covering (mis)matched spacer-target interfaces with varying spacer lengths (Figure 4). DeepEnsEmbCas9 naive outperformed on several mismatched interface tests, suggesting that pooling of mismatched SpCas9 variant data into one dataset improves model performance. DeepEnsEmbCas9 naive also out-performed DeepSmallCas9 on wild-type and high-fideltiy SaCas9 variants, again suggesting that the pooling of data from similar variants improves model performance. Comparing between models, Deep-EnsEmbCas9 naive has slightly higher Spearman performance than DeepEmbCas9, which is explained by DeepEnsEmbCas9 naive’s ensembling of 20 predictions. Evidenced by DeepEmbCas9-MVE and DeepEnsEmbCas9s’ Spearman performances, use of mean-variance estimation and a Gaussian negative log-likelihood loss objective reduced Spearman performance, suggesting that the point estimates made by DeepEmbCas9 or DeepEnsEmbCas9 naive are overconfident. It is likely that hyperparameter optimization would further boost DeepEnsEmbCas9 naive’s in-distribution performance.

Unlike previous Cas9 cleavage activity models, DeepEnsEmbCas9 naive in theory is able to make indel frequency predictions for any Cas9 nucleases, especially type II-A nucleases given our training dataset’s bias towards SpCas9 variants. We demonstrated this in leave-one-nuclease-out extrapolation tasks, where DeepEnsEmbCas9 naive attained comparable extrapolation performance to the best-performing individual activity prediction models. As expected, extrapolation performance deteriorates when extrapolating to nucleases like St1Cas9, Nm1Cas9 and Nm2Cas9, highlighting the need for more cleavage activity data from diverse Cas9 nucleases and variants.

### 4.3 Uncertainty estimation

Our study adds DeepEnsEmbCas9 to the family of uncertainty-aware CRISPR-Cas9 cleavage activity prediction models, including crispAI [154], CRISPR-DBA [164] and CRISPR DeepEnsemble [165]. Theory-wise, DeepEnsEmbCas9’s built-in uncertainty estimates prevents the model from being over-confident, and allows users to judge the reliability of DeepEnsEmbCas9’s prediction, especially when using DeepEnsEmbCas9 to make extrapolation predictions on type II-B or II-C Cas9 nucleases, both of which are severely underrepresented by the 40 Cas9 variants used in our training dataset.

Analyzing left-tailed and CI-based quantile calibration curves in the in-distribution setting, we see that DeepEmbCas9-MVE (Figures S32 and S33) and DeepEnsEmbCas9 (Figures 5 and S31) are quantile calibrated (i.e., have good quality uncertainty estimates), whereas DeepEnsEmbCas9 naive (Figures S34 and S35) is not quantile calibrated, as corroborated by significantly higher left-tailed and CI-based quantile calibration errors for DeepEnsEmbCas9 naive compared to the low calibration errors for DeepEmbCas9-MVE and DeepEnsEmbCas9 (Figures S36 and S37) on most benchmark test sets. Given that mean-variance estimation and ensembling of output predictions captures aleatoric and epistemic uncertainty [166], respectively, similar quantile calibration errors between DeepEmbCas9-MVE and DeepEnsEmbCas9 suggests that DeepEnsEmbCas9’s aleatoric uncertainty is larger than epistemic uncertainty. Coupled with DeepEnsEmbCas9’s high in-distribution test Spearman correlations, it follows that there is a tradeoff between Spearman performance and quantile correlation among the 3 DL models considered.

As for the leave-one-nuclease-out extrapolation setting, we see that DeepEmbCas9-MVE (Figure S40 and S41) and DeepEnsEmbCas9 (Figure S38 and S39) exhibit mixed level of quantile calibration dependent on the nuclease and study, whereas DeepEnsEmbCas9 naive (Figures S42 and S43) is not quantile calibrated. Interestingly, DeepEmbCas9-MVE and DeepEnsEmbCas9 have higher left-tailed and CI-based quantile calibration errors for Wang et al. [100], Kim, Kim et al. [95] and Kim et al. [141], which is likely a result of training data imbalance arising from discrepancies in genome editing experimental protocols among the different studies.

### 2.2 SHAP importance analysis

SHAP analysis on DeepEmbCas9 primarily reflects the importance of the PAM sequence and Cas9’s PAM-interacting (PI) domain. Specifically, −2G and −3G PAM features are shown to be very important (Figure 6G and 6I), as corroborated by high SHAP importance of feature group “PAM + downstream + PAM downstream Tm” in Figure 6A and feature group “NTS” in Figure 6B. This is consistent with SpCas9 and its increased fidelity variants recognizing NGG PAM. SHAP importances in the NTS-downstream region are observed to taper off beyond PAM position −6 (Figure 6G and 6I), a result of 36 out of 40 Cas9 variants having ≤4 nt PAM. Cas9’s PI domain also has high importance, as shown by feature groups “Cas9 ESM-C-600M REC insert”, “PLL-WED-PI” and”PI” in Figure 6A, 6C and 6D, respectively. This is consistent with Cas9’s PI domain participation in PAM binding — the first step towards Cas9 cleavage [148].

SHAP analysis also shows the importance of the PAM-proximal heteroduplex region on CRISPR-Cas9 cleavage activity. In particular, the spacer sequence is shown to be important, as demonstrated by the high SHAP importance of “spacer + spacer MFE + spacer GCcount” and “spacer” feature groups in Figure 6A, 6B and 6E, respectively. Heteroduplex positions +1 to +7 have high SHAP importance (Figure 6F), relative to other heteroduplex nucleotide positions, corroborating with the low spacer-target mismatch tolerance in the PAM-proximal seed region during R-loop formation [1]. The +1G spacer nucleotide is consistent with feature importance analysis of other prediction tools [72, 17, 77, 100, 104], with the literature reporting said nucleotide being associated with improved SpCas9 cleavage activity possibly due to its importance during the SpCas9 loading of sgRNA [167].

Cas9 domains apart from Cas9’s PI domain are also influential in DeepEmbCas9’s cleavage activity prediction. Surprisingly, we see REC insert, L2, REC1-B and REC2 listed among the top 5 important Cas9 regions (Figure 6D). Given that:

- the REC_insert embedding is a non-zero vector only for SpCas9/Sc++ (encoding the REC2 domain) and St1Cas9 (encoding the Wing domain);
- the L2 embedding is a non-zero vector for all Cas9 nucleases except for CjCas9 and enCjCas9;
- the REC1-B embedding is a non-zero vector for only SpCas9, St1Cas9 and Sc++; and
- the REC2 embedding is a non-zero vector for only SpCas9 (encoding the REC3 domain) and St1Cas9/CjCas9/Nm1Cas9/Nm2Cas9/Sc++ (encoding the REC2 domain),

it is possible that DeepEmbCas9 took advantage of zero-valued features from these Cas9 regions — where Cas9 protein architectures differ — in order to distinguish between different Cas9 nucleases.

### 4.5 SHAP importance analysis of nuclease pairs

When using the framework used for assessing the impact of input features on DeepEmbCas9’s predicted activity change in 39 Cas9 nuclease pairs, we see that SHAP importance varies substantially only for Cas9 and NTS (Figure 7A). Together with Cas9 SHAP importance being positively correlated with NTS SHAP importance (Figure 7B), the two observations highlight the presence of feature interactions between the Cas9 and NTS components. Plotting SHAP domain and region importances for the 39 Cas9 nuclease pairs, we see that L2 region in the Linker domain (abbreviated as Linker/L2 onwards) dominates in SHAP importance when mutating from SpCas9 to increased-fidelity SpCas9 variants (i.e., eSpCas9(1.1), SpCas9-HF1, evoCas9, HypaCas9, Sniper-Cas9 and xCas9; Figure 7C). Interestingly, xCas9 does not have high PLL-WED-PI importance, possibly hinting at the conflation between SpCas9 and xCas9’s PAM preferences by DeepEmbCas9. VRQR-HF1 has higher PLL-WED-PI importance in addition to Linker/L2 importance, which is consistent with the Cas9 PI domain VRQR mutations in VRQR-HF1. evoCas9 has high REC2 (i.e., SpCas9 REC3) importance in addition to high Linker/L2 importance, which is consistent with the 4 SpCas9 REC3 mutations possessed by evoCas9. QQR1’s low Linker/L2 and PLL-WED-PI/WED importance is likely due to QQR1’s overall low cleavage activity. We observe similar importance patterns for SaCas9 and SlugCas9 high-fidelity nuclease pairs (Figure 7E).

In contrast, we see that the WED region in the PLL-WED-PI domain (abbreviated as PLL-WED-PI/WED onwards) dominates in SHAP importance for all PAM-altered SpCas9 variants except xCas9 and QQR1. Moreover, only xCas9 and QQR1 lack mutations at WED residue D1135 among the PAM-altered SpCas9 variants. Combined, this suggests the use of D1135-related features in the WED part of the Cas9 pLM embedding by DeepEmbCas9 for Cas9 cleavage activity prediction (Figure 7E). Indeed, D1135 is a residue which interacts with the minor groove of the PAM duplex via electrostatic repulsion between the negatively charged aspartate and sugar-phosphate backbone, whereas:

- D1135V in VQR, VRER and VRQR stabilizes the PAM duplex by replacing the electrostatic repulsion with van der Waals forces between valine and the sugar-phosphate backbone;
- D1135L in SpG and SpRY acts similarly to D1135V, but introduces a hydrophobic bulky leucine instead, which together with S1136V sterically pushes the third PAM base towards Q1335; and
- D1135N is a consensus mutation found during directed evolution when developing SpCas9-NRRH, SpCas9-NRTH and SpCas9-NRCH.

We also observe similar importance patterns for PAM-altered SaCas9 and SauriCas9 mutants (Figure 7F).

### 4.6 Limitations

We acknowledge several limitations in this study. Regarding data, there is an uneven amount of Cas9 nucleases and gRNA scaffolds in our training and test datasets, so DeepEmbCas9 may overfit certain experimental configuration types from specific studies. Due to limited DNA/RNA bulge data in existing indel frequency library screens, e.g., in Seo et al. [114], this study does not consider guide-target interfaces with DNA/RNA bulges. Library data used in this study also does not contain interfaces with 4-6 mismatches, limiting DeepEmbCas9’s predictive accuracy on such interfaces.

As for the embeddings, the Cas9 pLM and sgRNA rLM embeddings used in DeepEmbCas9 do not account for Cas9-gRNA scaffold interactions, so the effect of such interactions would need to be learned from the activity labels. DeepEmbCas9 also only accept Cas9 proteins with known domain boundaries as input, since this information is required for generating Cas9 pLM embeddings from the per-residue pLM embedding matrix. For analogous reasons, DeepEmbCas9 can only work with sgRNA scaffolds that have similar structure to those in our dataset.

With respect to SHAP interpretation, the low number of Cas9 variants limits the accuracy of the SHAP importances generated for the protein domains. The low count is also why we did not consider per-Cas9-residue protein representations and by extension per-Cas9-residue SHAP importances. There are also limitations to using SHAP feature importance scores for ML model interpretability, as suggested by Kumar et al. [168].

### Conclusion

In this study, we developed a family of 4 DeepEmbCas9 models — DeepEmbCas9, DeepEmbCas9-MVE, DeepEnsEmbCas9 naive and DeepEnsEmbCas9 — for cleavage activity prediction of Cas9 variants by:

1. representing all three components of the CRISPR-Cas9 complex, i.e., sgRNA, DNA and Cas9, as input features;
2. using a unified guide-target interface to align spacer and target sequences from different Cas9 nucleases;
3. adopting inductive biases compatible with Cas9’s biophysical mechanism; and
4. training on a curated dataset with *>*1.75 million datapoints spanning 40 Cas9 variants and 16 gRNA scaffolds.

Obtained by ensembling predictions, DeepEnsEmbCas9 naive attains comparable performance in both in-distribution and leave-one-nuclease-out extrapolation settings when compared to suitable individual cleavage activity models. We also built DeepEnsEmbCas9, trading off a small Spearman performance drop with well-calibrated uncertainty estimates. SHAP importance analysis of DeepEmbCas9 on all benchmark test set datapoints reaffirms the structural and functional importance of Cas9’s PLL-WED-PI domain and the PAM sequence for Cas9 binding — a prerequisite for Cas9 cleavage. Furthermore, SHAP importance analysis of DeepEmbCas9 on 39 nuclease pairs show that Linker and PLL-WED-PI features contribute significantly to predicted activity change for increased-fidelity and PAM-altering Cas9 mutations, respectively.

DeepEmbCas9 models confer advantages over existing Cas9 activity models leveraging Cas9 protein information. Compared to PLM CRISPR [96], DeepEmbCas9 trains on 33 more Cas9 variants spanning beyond increased-fidelty SpCas9 variants. Unlike STING CRISPR [112], DeepEmbCas9 scales better with increasing guide-target training data while maintaining the ability to assess Cas9 domain importances. In contrast to PAMmla [126] and CICERO [127], DeepEmbCas9 models directly address the problem of cleavage activity prediction rather than the subproblem of Cas9-target PAM binding prediction. Altogether, DeepEmbCas9 models serve as the first step towards generalistic interpretable DL-based models capable of predicting cleavage activity for diverse combinations of wild-type/engineered/pLM-generated nucleases and guide-target interfaces in the Cas9 protein family.

## 6 Acknowledgements

We acknowledge the use of the University of Oxford Advanced Research Computing (ARC) facility in carrying out this work (http://dx.doi.org/10.5281/zenodo.22558).

## Author contributions

Conceptualization: JM, PM

Methodology: JM, PM

Investigation: JM, PM

Visualization: JM, PM

Funding Acquisition: JM, PM

Project administration: JM, PM

Supervision: PM

Writing - original draft: JM, PM

Writing - review & editing: JM, PM

## 8 Data availability

RNA-FM model weights are available at https://github.com/ml4bio/RNA-FM. RiNALmo model weights are available at https://github.com/lbcb-sci/RiNALMo. BEACON-B and BEACON-B512 model weights are available at https://github.com/terry-r123/RNABenchmark. Model weights for ProtT5 are available at https://huggingface.co/Rostlab/prot_t5_xl_half_uniref50-enc. Model weights for Ankh-Large are available at https://huggingface.co/ElnaggarLab/ankh-large. Model weights for ESM3, ESM-C-300M, ESM-C-600M and ESM-C-6B are available at https://github.com/evolutionaryscale/esm. Inference code and training data for DeepEmbCas9 models will be made available upon publication.

