## Supplementary Information for "DeepEmbCas9: Cas9 coevolution and sgRNA structural information for CRISPR-Cas9 cleavage activity prediction"

### 1 Supplementary methods

#### 1.1 Model comparisons

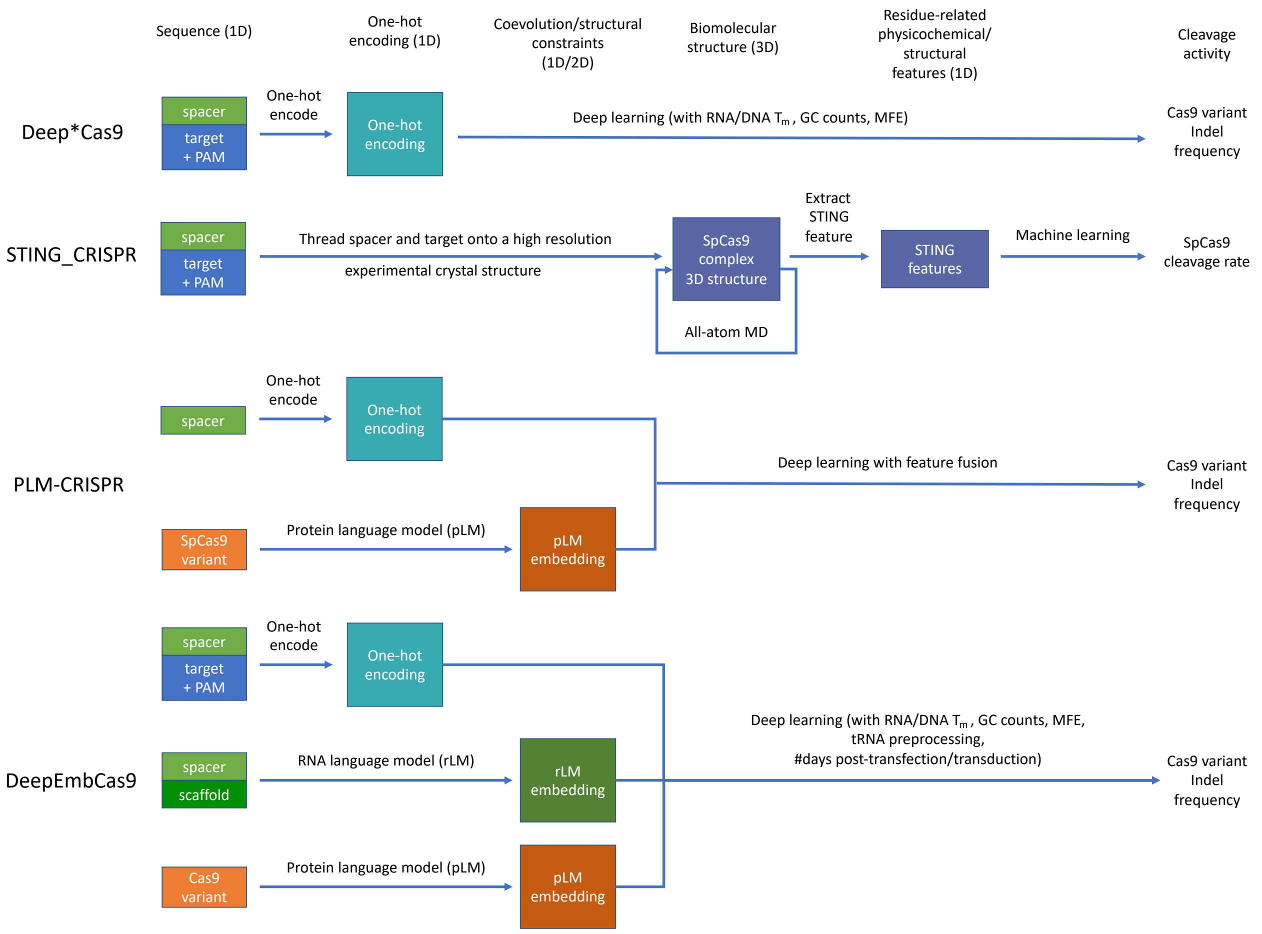


Figure S1: ML/DL model comparison between individual Cas9 cleavage activity tools (i.e., DeepSpCas9 [1], DeepHF [2], DeepxCas9 [3], DeepSpCas9-NG [3], DeepSpCas9variants [4], DeepSmallCas9 [5], DeepSniper [6], DeepCas9variants [7]), STING_CRISPR, PLM-CRISPR and DeepEmbCas9. A single ML/DL model (DeepEmbCas9) is built for 40 Cas9 variants, while ML/DL-based individual Cas9 cleavage activity models (Deep*Cas9) are built for each. PLM_CRISPR only considers 7 SpCas9 variants (including wild type SpCas9). Owing to limited comptuational resources, STING_CRISPR can only predict wild type SpCas9 cleavage activity for an extremely limited set of guide-target interfaces.

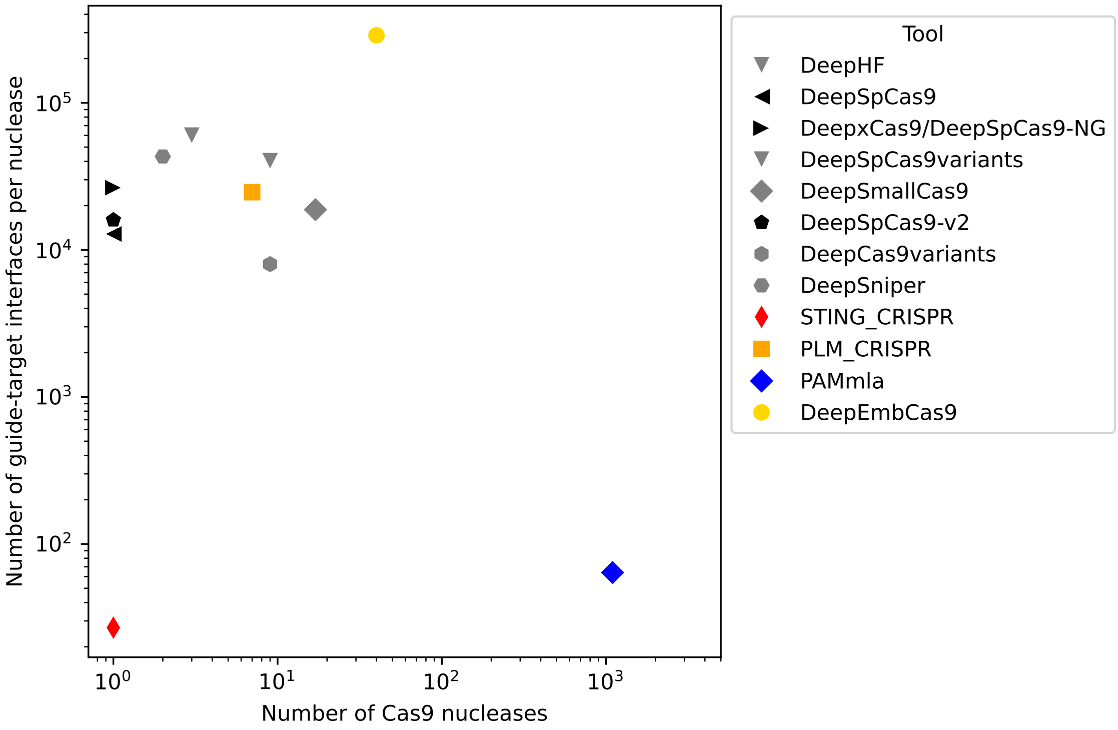


Figure S2: Number of Cas9 nucleases and average number of guide-target interfaces per nuclease used for training DeepHF, DeepSpCas9, DeepxCas9, DeepSpCas9-NG, DeepSPCas9variants, DeepSmall-Cas9, DeepSpCas9-v2, DeepCas9variants, DeepSniper, STING_CRISPR, PLM_CRISPR, PAMmla and DeepEmbCas9.

#### 1.2 Dataset

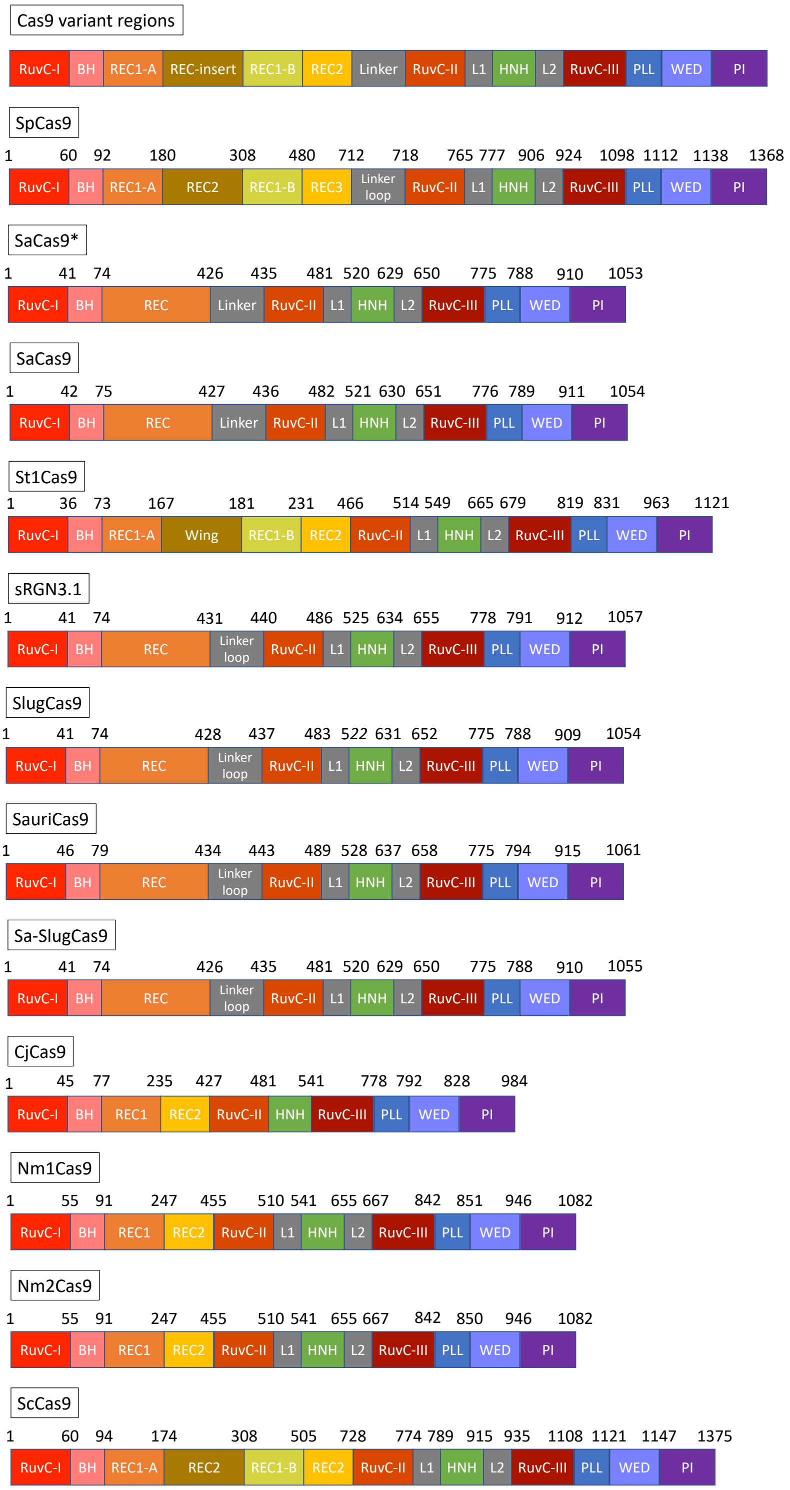


Figure S3: Cas9 regions used for partitioning Cas9 variants. Top row shows the list of Cas9 regions, and the subsequent rows show the partitioning for the base Cas9 nucleases SpCas9, SaCas9*, SaCas9, St1Cas9, sRGN3.1, SlugCas9, SauriCas9, Sa-SlugCas9, CjCas9, Nm1Cas9, Nm2Cas9 and ScCas9. Identical partitionings are used for Cas9 variants with the same base nuclease.

| Dataset column | Description |
| --- | --- |
| Study | Publication(s) where the datapoint(s) were collected |
| Library | The oligonucleotide pool containing the datapoint(s) |
| Table | Supplementary table/data in the study containing the datapoint(s) |
| Sheet | Datasheet name within the Excel file containing the datapoint(s) |
| src_idx | Row number within the Excel datasheet |
| n_data | Total number of biological/technical replicates across all studies |
| Partition | Training or test datapoint(s) |
| Partition (source) | Data partition(s) for each indel frequency label listed in "Background subtracted indel frequencies (%)" |
| Barcode | Barcode(s) for each experiment in each study |
| Spacer sequence (raw) | The spacer sequence without N-padding (typically $\geq$ 20nt) |
| Target context sequence (raw) | The target sequence with 5' upstream and 3' downstream context without N-padding (typically $\geq$ 30nt) |
| Spacer sequence | The spacer sequence with N-padding (40nt) |
| Target context sequence | The target sequence with 5' upstream and 3' downstream context, with N-padding (40nt) |
| Variant | Name of the Cas9 nuclease with nuclear localization signal (NLS), FLAG tag and P2A peptide |
| Nuclease | Name of the Cas9 nuclease only |
| gRNA scaffold | Name of the guide RNA scaffold |
| Day | The number of days post-transfection/transduction prior to genomic DNA isolation from edited cells |
| tRNA feature | Binary feature indicating the use of a $tRNA^{Gln}$-N20 sgRNA |
| Background subtracted indel frequencies (%) | Sets of indel frequencies for each study listed in "Study" |
| Mean background subtracted indel frequency (source, %) | Replicate-weighted mean of indel frequencies for each study |
| Mean background subtracted indel frequency (%) | Replicate-weighted mean indel frequency |

Table S1: List of column names and descriptions for the curated CRISPR-Cas9 indel frequency dataset.

| No. of mismatches | No. of datapoints |
|---|---|
| 0 | 782201 |
| 1 | 858316 |
| 2 | 71367 |
| 3 | 33802 |
| 4 | 1000 |

Table S2: Number of guide-target mismatches in the Cas9 variant indel frequency dataset.

| gRNA scaffold | Repeat-antirepeat length | tracrRNA (excluding antirepeat) length | poly(U) tail length |
|---|---|---|---|
| SpCas9 scaffold 1 | 30 | 46 | 6 |
| SpCas9 scaffold 1 (5T) | 30 | 46 | 5 |
| SpCas9 scaffold 2 | 40 | 46 | 6 |
| SaCas9 scaffold 1 | 34 | 42 | 6 |
| SaCas9 scaffold 2 | 42 | 42 | 6 |
| SaCas9 scaffold 3 | 34 | 42 | 6 |
| NmCas9 scaffold 1 | 52 | 69 | 6 |
| NmCas9 scaffold 2 | 40 | 61 | 6 |
| NmCas9 scaffold 3 | 36 | 61 | 6 |
| CjCas9 scaffold 1 | 28 | 45 | 6 |
| CjCas9 scaffold 2 | 26 | 45 | 6 |
| St1Cas9 scaffold 1 | 79 | 46 | 6 |
| St1Cas9 scaffold 2 | 79 | 46 | 6 |
| St1Cas9 scaffold 3 | 37 | 46 | 6 |
| St1Cas9 scaffold 4 | 32 | 46 | 6 |
| St1Cas9 scaffold 5 | 34 | 46 | 6 |

Table S3: Length of sgRNA regions for the 16 gRNA scaffolds in this study.

##### 1.3 Protein sequences

Codon sequences for SpCas9-NLS-FLAG-P2A (Addgene, #52962), eSpCas9(1.1)-NLS-FLAG-P2A (Addgene, #138555), SpCas9-HF1-NLS-FLAG-P2A (Addgene, #138556), HypaCas9-NLS-FLAG-P2A (Addgene, #138557), evoCas9-NLS-FLAG-P2A (Addgene, #138558), xCas9-NLS-FLAG-P2A (Addgene, #138565), Sniper-Cas9-NLS-FLAG-P2A (Addgene, #138559), VQR-NLS-FLAG-P2A (Addgene, #138560), VRER-NLS-FLAG-P2A (Addgene, #138561), VRQR-NLS-FLAG-P2A (Addgene, #138562), VRQR-HF1-NLS-FLAG-P2A (Addgene, #138563), QQR1-NLS-FLAG-P2A (Addgene, #138564), SpCas9-NG-NLS-FLAG-P2A (Addgene, #138566), Sniper2L-NLS-FLAG-P2A (Addgene, #193856), Sniper2P-NLS-FLAG-P2A (Addgene, #193857) were obtained from their respective Addgene plasmids. Codon sequences for the small Cas9 variants (including NLS-SpCas9-NLS-FLAG-P2A) in Seo et al. [5] were obtained from Supplementary Note 1 of the study, and codon sequences from nucleases in Kim, Choi et al. [7] were computationally derived from the gBlocks Gene Fragments and PCR primers listed in Supplementary Table 9 of the study. Biopython [8] was then used to computationally derive protein sequences from codon sequences. The protein sequence of NLS-SaCas9*-NLS-FLAG was derived from NLS-SaCas9-NLS-FLAG by removing the glycine residue located at the start of SaCas9's RuvC-I subdomain.

| Nuclease | Base nuclease | Mutations | Primary PAM |
|---|---|---|---|
| SpCas9 | SpCas9 | WT | NGG |
| eSpCas9(1.1) | SpCas9 | K848A/K1003A/R1060A | NGG |
| SpCas9-HF1 | SpCas9 | N497A/R661A/Q695A/Q926A | NGG |
| HypaCas9 | SpCas9 | N692A/M694A/Q695A/H698A | NGG |
| evoCas9 | SpCas9 | M495V/Y515N/K526E/R661Q | NGG |
| xCas9 | SpCas9 | A262T/R324L/S409I/E480K/E543D/M694I/E1219V | NG, GAA, GAT |
| Sniper-Cas9 | SpCas9 | F539S/M763I/K890N | NGG |
| VQR | SpCas9 | D1135V/R1335Q/T1337R | NGAN, NGCG |
| VRER | SpCas9 | D1135V/G1218R/R1335E/T1337R | NGCG |
| VRQR | SpCas9 | D1135V/G1218R/R1335Q/T1337R | NGAH |
| VRQR-HF1 | SpCas9 | N497A/R661A/Q695A/Q926A/D1135V/G1218R/R1335Q/T1337R | NGAH |
| QQR1 | SpCas9 | G1218R/N1286Q/I1331F/D1332K/R1333Q/R1335Q/T1337R | NAAG |
| SpCas9-NG | SpCas9 | L1111R/D1135V/G1218R/E1219F/A1322R/R1335V/T1337R | NG |
| sRGN3.1 | sRGN3.1 | WT | NNGG |
| SlugCas9 | SlugCas9 | WT | NNGG |
| SaCas9 | SaCas9 | WT | NNGRRT |
| SauriCas9 | SauriCas9 | WT | NNGG |
| Sa-SlugCas9 | Sa-SlugCas9 | WT | NNGG |
| SaCas9* | SaCas9* | WT | NNGRRT |
| SaCas9-KKH | SaCas9 | E799K/N985K/R1032H | NNNRRT |
| eSaCas9 | SaCas9 | R516A/Q517A/R671A/G672A | NNGRRT |
| efSaCas9 | SaCas9 | N277D | NNGRRT |
| SauriCas9-KKH | SauriCas9 | Q804K/Y989K/R1036H | NNRG |
| SlugCas9-HF | SlugCas9 | R263A/N431A/T437A/R672A | NNGG |
| SaCas9-HF | SaCas9 | R262A/N430A/N436A/R671A | NNGRRT |
| SaCas9-KKH-HF | SaCas9 | R262A/N430A/N436A/R671A/E799K/N985K/R1032H | NNNRRT |
| St1Cas9 | St1Cas9 | WT | NNRGAA |
| Nm1Cas9 | Nm1Cas9 | WT | NNNNGATT |
| enCjCas9 | CjCas9 | L74Y/D916K | NNNVRYAC |
| CjCas9 | CjCas9 | WT | NNNVRYAC |
| Nm2Cas9 | Nm2Cas9 | WT | NNNNCCA |
| SpCas9-NRRH | SpCas9 | I322V/S409I/E427G/R654L/R753G/R1114G/D1135N/V1139A/D1180G/E1219V/Q1221H/A1320V/R1333K | NRRH |
| SpCas9-NRTH | SpCas9 | I322V/S409I/E427G/R654L/R753G/R1114G/D1135N/D1180G/G1218S/E1219V/Q1221H/P1249S/E1253K/P1321S/D1332G/R1335L | NRTH |
| SpCas9-NRCH | SpCas9 | I322V/S409I/E427G/R654L/R753G/R1114G/D1135N/E1219V/D1332N/R1335Q/T1337N/S1338T/H1349R | NRCH |
| SpG | SpCas9 | D1135L/S1136W/G1218K/E1219Q/R1335Q/T1337R | NGN |
| SpRY | SpCas9 | A61R/L1111R/D1135L/S1136W/G1218K/E1219Q/N1317R/A1322R/R1333P/R1335Q/T1337R | NRN > NYN |
| Sc++ | Sc++ | I365A/G366D/I367K/H369L/T373S/T374G/Q379E/T1227K | NNG |
| Sniper2L | SpCas9 | F539S/M763I/K890N/E1007L | NGG |
| Sniper2P | SpCas9 | F539S/M763I/K890N/E1007P | NGG |

Table S4: List of 39 Cas9 nucleases considered in this study, with WT denoting wild type. All SpCas9 variants and Sc++ appear as Cas9-NLS-FLAG-P2A, and all Small Cas9s appear as NLS-Cas9-NLS-FLAG-P2A in the dataset, except for SpCas9, which appears as NLS-Cas9-NLS-FLAG-P2A in [5] and as Cas9-NLS-FLAG-P2A in all other studies, thus resulting in 40 Cas9 proteins observed in the dataset.

#### 1.4 Input feature encodings

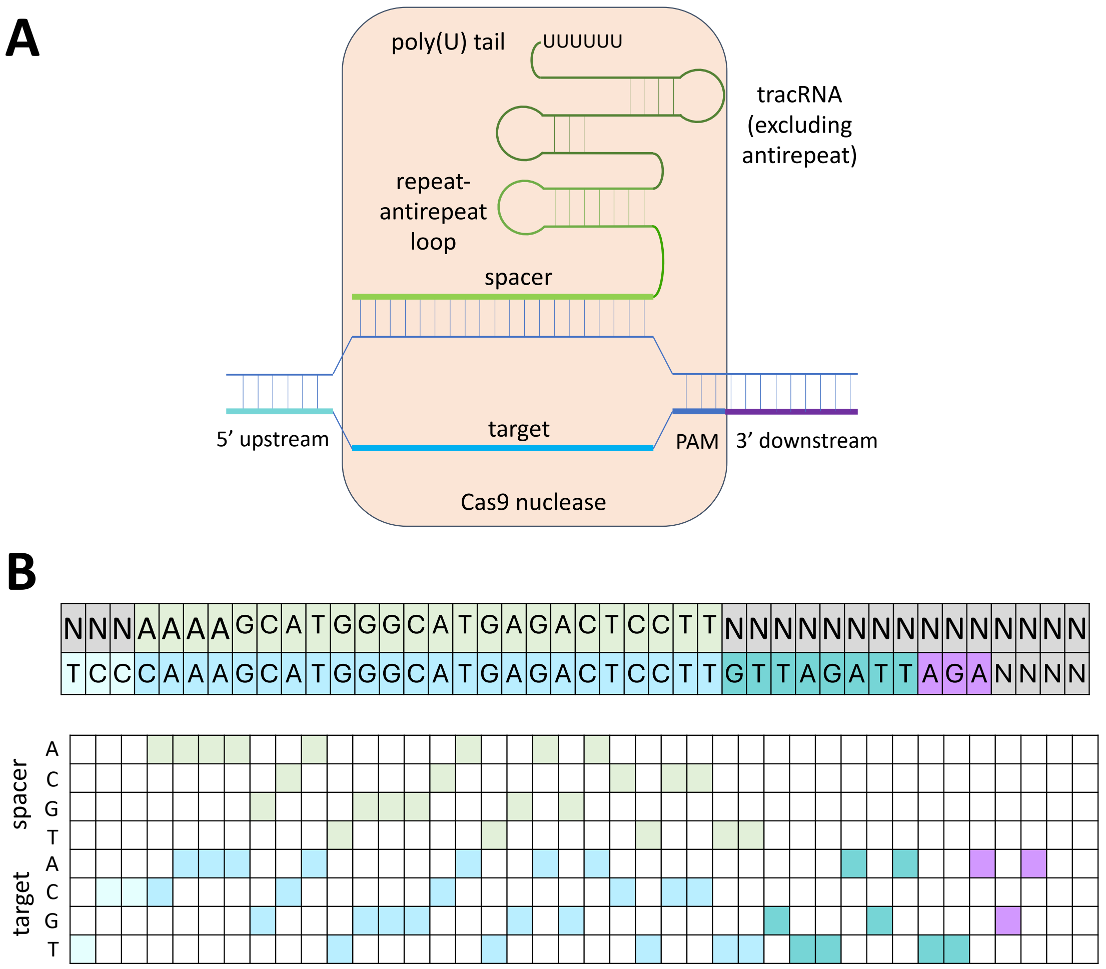


Figure S4: (A) Schematic representation of the guide-target-Cas9 variant R-loop complex. (B) Unified guide-target interface for an example mismatched Nm1Cas9 interface (top) and its one-hot encoding (bottom).

#### 1.5 Mean-variance estimation

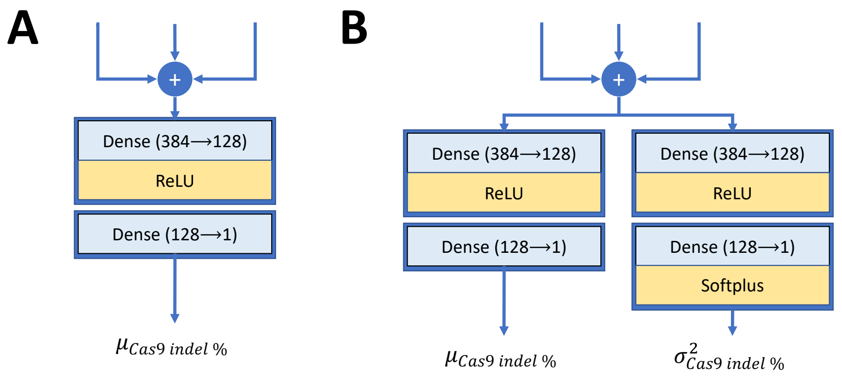


Figure S5: (A) Deterministic (mean) output head used in DeepEmbCas9 and DeepEmbCas9_naive (B) Mean and variance output heads used in DeepEmbCas9-MVE and DeepEnsEmbCas9.

### 2 Supplementary results

#### 2.1 In-distribution performance

| pLM (↓), rLM(→) | RNA-FM | BEACON-B512 | BEACON-B | evo-1-8k | RiNALMo | Average pLM performance |
|---|---|---|---|---|---|---|
| ESM-C-600M | 0.901 ± 0.002 | 0.900 ± 0.002 | **0.903 ± 0.002** | 0.895 ± 0.003 | 0.899 ± 0.001 | 0.900 ± 0.003 |
| ESM-C-300M | 0.898 ± 0.002 | 0.896 ± 0.002 | 0.898 ± 0.001 | 0.896 ± 0.002 | 0.898 ± 0.002 | 0.897 ± 0.002 |
| ESM-C-6B | 0.898 ± 0.004 | 0.894 ± 0.002 | 0.899 ± 0.002 | 0.889 ± 0.004 | 0.896 ± 0.004 | 0.895 ± 0.005 |
| ProtT5 | 0.895 ± 0.002 | 0.885 ± 0.006 | 0.890 ± 0.004 | 0.889 ± 0.003 | 0.894 ± 0.002 | 0.890 ± 0.005 |
| Ankh-large | 0.892 ± 0.002 | 0.886 ± 0.001 | 0.889 ± 0.004 | 0.889 ± 0.002 | 0.890 ± 0.003 | 0.889 ± 0.003 |
| gLM2_650M | 0.826 ± 0.003 | 0.823 ± 0.005 | 0.799 ± 0.064 | 0.820 ± 0.006 | 0.824 ± 0.007 | 0.818 ± 0.028 |
| ESM3 | 0.828 ± 0.001 | 0.795 ± 0.065 | 0.799 ± 0.068 | 0.792 ± 0.070 | 0.752 ± 0.100 | 0.793 ± 0.068 |
| Average rLM performance | 0.877 ± 0.032 | 0.868 ± 0.045 | 0.868 ± 0.055 | 0.867 ± 0.047 | 0.865 ± 0.063 | N/A |

Table S5: DeepEmbCas9's performance on the validation sets during five-fold cross validation, with results sorted in descending order of averaged pLM and rLM performances.

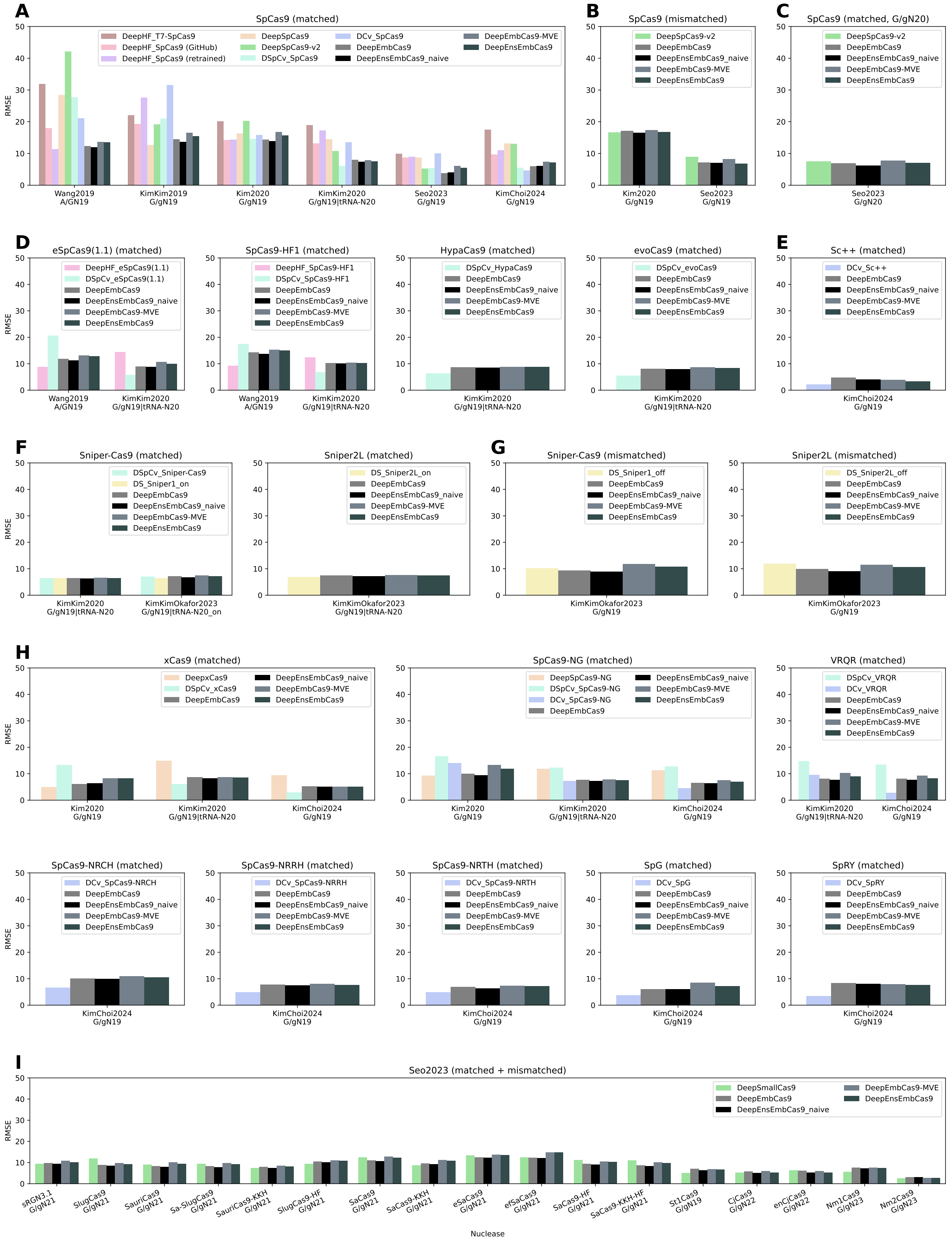


Figure S6: Benchmark test RMSE correlation comparison for DeepEmbCas9, DeepEnsEmbCas9_naive, DeepEmbCas9-MVE and DeepEnsEmbCas9 against DeepHF, DeepSpCas9, DeepxCas9, DeepSpCas9-NG, DeepSpCas9variants, DeepSmallCas9, DeepSpCas9-v2, DeepCas9variants and DeepSniper across 39 Cas9 nucleases. The test sets consist of (A) matched $AN_{19}$, $G/gN_{19}$ and $tRNA^{Gln}$-$N_{20}$ wild type SpCas9 interfaces; (B) mismatched $G/gN_{19}$ wild type SpCas9 interfaces; (C) matched $G/gN_{20}$ wild type SpCas9 interfaces; (D,E,F) matched $AN_{19}$, $G/gN_{19}$ and $tRNA^{Gln}$-$N_{20}$ interfaces for 6 increased-fidelity SpCas9 variants and Sc++; (G) mismatched $G/gN_{19}$ interfaces for Sniper variants; (H) matched $G/gN_{19}$ and $tRNA^{Gln}$-$N_{20}$ interfaces for 8 PAM-altered SpCas9 variants; and (I) (mis)matched interfaces for 17 wild type or engineered small Cas9 nucleases.

##### 2.1.1 In-distribution performance comparisons of DeepEmbCas9, DeepEnsEmbCas9 and DeepEmbCas9-MVE

Among the 51 benchmark test sets, DeepEmbCas9 attains higher Spearman correlation than all individual activity prediction tools on 10 test sets(Figure 4, gray bars), namely 3 mismatched G/g$N_{19}$ interface test sets for SpCas9 and Sniper-Cas9 (Figures 4B and G) and 7 small Cas9 test sets (2 SlugCas9 variants, 4 SaCas9 variants and SauriCas9) from Seo et al. [5] (Figure 4I). As for the remaining 41 test sets, DeepEmbCas9 has an average Spearman correlation drop of $4.80 \times 10^{-2}$ compared to the best-performing individual activity prediction tools, with the test set containing (mis)matched G/g$N_{23}$ Nm2Cas9 interfaces from Seo et al. [5] yielding the largest Spearman drop of 0.175 (DeepSmallCas9's 0.559 vs. DeepEmbCas9 0.383). Among test sets outside of the 51 benchmark test sets which lack proper baselines, DeepEmbCas9 attains 0.451-0.917 Spearman correlation in 9 out of 10 test sets (Figures S10 rows 2-3, S11, S15, S21 row 3, S22 row 3 and S23 row 3).

Among the 51 benchmark test sets, DeepEnsEmbCas9 attains higher Spearman correlation than all individual activity prediction tools on 8 test sets (Figure 4, dark slate gray bars), which consist of the test set containing matched G/g$N_{19}$ SpCas9 interfaces from Kim et al. [3] (Figure 4A), 3 mismatched G/g$N_{19}$ interface test sets for SpCas9 and Sniper-Cas9 (Figures 4B and G), and 3 small Cas9 test sets (SlugCas9, Sa-SlugCas9 and enCjCas9) from Seo et al. [5] (Figure 4I). As for the remaining 43 test sets, DeepEnsEmbCas9 has an average Spearman correlation drop of $4.13 \times 10^{-2}$ compared to the best-performing individual activity prediction tools, with the test set containing matched A/G$N_{19}$ SpCas9-HF1 interfaces from Wang et al. [2] yielding the largest Spearman drop of 0.194 (DeepHF_SpCas9-HF1's 0.881 and DeepEnsEmbCas9's 0.687). Among test sets outside of the 51 benchmark test sets which lack proper baselines, DeepEnsEmbCas9 attains 0.414-0.916 Spearman correlation in 9 out of 10 test sets (Figures S10 rows 2-3, S11, S15, S21 row 3, S22 row 3 and S23 row 3).

Among the 51 benchmark test sets, DeepEmbCas9-MVE attains higher Spearman correlation than all individual activity prediction tools on 3 test sets (Figure 4, slate gray bars), specifically the test set with matched G/g$N_{19}$ SpCas9 interfaces from Kim et al. [3] and 2 test sets with mismatched G/g$N_{19}$ SpCas9 interfaces (Figure 4A-B). As for the remaining 48 test sets, DeepEmbCas9-MVE has an average Spearman correlation drop of $5.40 \times 10^{-2}$ compared to the best-performing individual activity prediction tools, with the test set containing matched G/g$N_{19}$ SpRY interfaces from Kim, Choi et al. [7] yielding the largest Spearman drop of 0.221 (DCv_SpRY's 0.934 vs. DeepEmbCas9-MVE 0.713). Among test sets outside of the 51 benchmark test sets which lack proper baselines, DeepEmbCas9-MVE attains 0.412-0.911 Spearman correlation in 9 out of 10 test sets (Figures S10 rows 2-3, S11, S15, S21 row 3, S22 row 3 and S23 row 3).

##### 2.1.2 Detailed analysis of DeepEmbCas9's in-distribution performance

DeepEmbCas9 trained using ESM-C-600M and BEACON-B embeddings yields Spearman and RMSE metrics comparable to those of individual activity prediction tools corresponding to the test set's study on 51 benchmark test sets from the 6 studies considered (Figures 4 and S6). Regarding matched SpCas9 interfaces (Figures 4A) with A/G$N_{19}$ sgRNAs from Wang et al. [2], DeepEmbCas9 (0.788) attain higher Spearman correlation for all SpCas9 activity prediction tools except DeepHF_SpCas9 (retrained, 0.821; GitHub, 0.797) (Figure S7 row 1). On matched G/$g_{19}$ SpCas9 test interfaces from Kim Kim et al. [1], DeepEmbCas9 (0.692) attains higher Spearman correlation for all SpCas9 activity prediction tools except DeepSpCas9 (0.773) and DeepHF (GitHub, 0.713; Figure S8). On matched G/$g_{19}$ SpCas9 test interfaces from Kim et al. [3], DeepEmbCas9 (0.905) attains similar Spearman correlation to DeepSpCas9variants (abbreviated DSpCv_SpCas9), and higher Spearman correlation for all other SpCas9 activity prediction tools (Figure S9 row 1). On matched G/$g_{19}$ and $tRNA^{Gln}$-$N_{20}$ SpCas9 test interfaces from Kim, Kim et al. [4], DeepEmbCas9 (0.927) attains higher Spearman correlation for all SpCas9 activity prediction tools except DeepSpCas9variants (0.937; Figure S12 row 1). On matched G/$g_{19}$ SpCas9 test interfaces from Seo et al. [5], DeepEmbCas9 (0.710) attains higher Spearman correlation for all SpCas9 activity prediction tools except DeepCas9variants (abbreviated

DCv_SpCas9, 0.776), DeepSpCas9variants (0.766), and DeepSpCas9-v2 (0.732; Figure S17 row 1). On matched G/$g_{19}$ SpCas9 test interfaces from Kim, Choi et al. [7], DeepEmbCas9 (0.733) attains higher Spearman correlation for all SpCas9 activity prediction tools except DeepCas9variants (0.762) and DeepSpCas9-v2 (0.744; Figure S17 row 1).

Regarding mismatched G/g$N_{19}$ SpCas9 test interfaces, DeepEmbCas9 (0.778 and 0.906) surpasses DeepSpCas9-v2 (0.773 and 0.874) in Spearman correlation for test datasets from Kim et al. [3] and Seo et al. [5](Figures 4B, S10 row 1 and S16 row 2). When combining matched and mismatched G/g$N_{19}$ SpCas9 interfaces in Kim et al. [3], DeepEmbCas9 (0.889) has higher Spearman correlation than DeepSpCas9-v2 (0.862; Figure S11 row 1). When combining matched and mismatched G/g/$N_{19}$ SpCas9 interfaces in Seo et al. [5], DeepEmbCas9 (0.808) attains lower SPearman correlation than DeepSpCas9-v2 (0.823; Figure S16 row 3). On matched G$N_{20}$ SpCas9 interfaces from Seo et al. [5], DeepEmbCas9 (0.345) has lower Spearman correlation compared to DeepSpCas9-v2 (0.358; Figures 4C and S16 row 4).

Next, we assess performances for 4 increased-fidelity SpCas9 variants and Sc++ (Figure 4D-E). Regarding eSpCas9(1.1) and SpCas9-HF1 on matched A/G$N_{19}$ interfaces from Wang et al. [2], DeepEmbCas9 (0.849 and 0.718) attains higher Spearman correlation than DeepSpCas9variants (0.449 and 0.511) but not DeepHF (0.886 and 0.881; Figure S7 rows 2-3). On matched G/g$N_{19}$ and $tRNA^{Gln}$-$N_{20}$ eSpCas9(1.1) and SpCas9-HF1 interfaces from Kim, Kim et al. [4], DeepEmbCas9 (0.849 and 0.790) attains higher Spearman correlation than DeepHF (0.759 and 0.725) but not DeepSpCas9variants (0.877 and 0.821; Figure S12 rows 2-3). For HypaCas9 and evoCas9 on matched G/$g_{19}$ interfaces from Kim, Kim et al. [4], DeepEmbCas9 attains lower Spearman correlation than DeepSpCas9variants (0.833 vs. 0.855 for HypaCas9; 0.587 vs. 0.647 for evoCas9; Figures S13 rows 1-2). For Sc++ on matched G/$g_{19}$ interfaces from Kim, Choi et al. [7], DeepEmbCas9 (0.573) attains lower Spearman correlation than DeepCas9variants (0.624; Figures 4E and S17 row 2).

We then examine performances for Sniper-Cas9 and Sniper2L on matched (Figure 4F) and mismatched (Figure 4G) interfaces. For Sniper-Cas9, on matched G/g$N_{19}$ and $tRNA^{Gln}$-$N_{20}$ interfaces from Kim, Kim et al. [4], DeepEmbCas9 (0.931) attains similar Spearman correlation to DeepSpCas9variants (0.936) and DeepSniper (0.935; Figure S13 row 3). On matched G/g$N_{19}$ and $tRNA^{Gln}$-$N_{20}$ Sniper-Cas9 interfaces from Kim, Kim, Okafor et al. [6], DeepEmbCas9 (0.926) also has similar Spearman correlation to DeepSpCas9variants (0.929) and DeepSniper (0.936; Figure S21 row 1). For Sniper2L, on matched G/g$N_{19}$ and $tRNA^{Gln}$-$N_{20}$ interfaces from Kim, Kim, Okafor et al. [6], DeepEmbCas9 attains similiar Spearman correlation (0.923) to DeepSniper (0.931; Figure S21 row 2). Combining matched and mismatched interfaces from Kim, Kim, Okafor et al. [6], DeepEmbCas9 (0.925 and 0.922) attains similar Spearman correlation to DeepSniper (0.929 and 0.924) for Sniper-Cas9 and Sniper2L, respectively (Figure S23 rows 1-2).

We subsequently examine performances for xCas9, SpCas9-NG, and 6 other PAM-altered SpCas9 variants (Figure 4H). For xCas9, on matched G/g$N_{19}$ interfaces from Kim et al. [3], DeepEmbCas9 (0.884) attains lower Spearman correlation than DeepxCas9 (0.913) and DeepSpCas9variants (0.886; Figure S9 row 2). On matched xCas9 interfaces from Kim, Kim et al. [4] and Kim, Choi et al. [7], DeepEmbCas9 (0.877 and 0.717) attains higher Spearman correlation than DeepxCas9 (0.844 and 0.704), but not for DeepSpCas9variants (0.927 and 0.740; Figure S14 row 1 and S18 row 1).

For SpCas9-NG, on matched G/g$N_{19}$ interfaces from Kim et al. [3], DeepEmbCas9 (0.879) attains higher Spearman correlation than DeepSpCas9variants (0.722) and DeepCas9variants (0.713), but not for DeepSpCas9-NG (0.904; Figure S9 row 3). On matched SpCas9-NG interfaces for test datasets from Kim, Kim et al. [4] and Kim, Choi et al. [7], DeepEmbCas9 (0.890 and 0.891) attains higher Spearman correlation than DeepSpCas9-NG (0.817 and 0.868) and DeepSpCas9variants (0.799 and 0.722), but not for DeepCas9variants (0.903 and 0.935; Figure S14 row 2 and S18 row 2).

Moving to SpCas9-VRQR (abbreviated as VRQR), on matched VRQR interfaces for test datasets from Kim, Kim et al. [4] and Kim, Choi et al. [7], DeepEmbCas9 (0.889 and 0.742) attains higher Spearman correlation than DeepCas9variants (0.941 and 0.772), but not for DeepSpCas9variants (0.799 and

0.717; Figures S13 row 3 and S18 row 3). As for matched interfaces for the other 5 SpCas9 variants, DeepEmbCas9 attains lower Spearman correlation than DeepCas9variants (0.848 vs. 0.936 for SpCas9-NRCH, 0.897 vs. 0.955 for SpCas9-NRRH, 0.890 vs. 0.935 for SpCas9-NRTH, 0.862 vs. 0.903 for SpG, and 0.764 vs. 0.934 for SpRY; Figures S19 and S20).

As for the (mis)matched small Cas9 variant interfaces, DeepEmbCas9 attains higher Spearman correlation than DeepSmallCas9 for SlugCas9 (0.922 vs. 0.916), SauriCas9 (0.914 vs. 0.905), Sa-SlugCas9 (0.922 vs. 0.901), SaCas9 (0.904 vs. 0.901), eSaCas9 (0.860 vs. 0.854), efSaCas9 (0.865 vs. 0.860), and SaCas9-KKH-HF (0.862 vs. 0.828), but not for sRGN3.1 (0.908 vs. 0.916), SauriCas9-KKH (0.848 vs. 0.855), SlugCas9-HF (0.789 vs. 0.810), SaCas9-KKH (0.902 vs. 0.928), SaCas9-HF (0.847 vs. 0.851), St1Cas9 (0.789 vs. 0.890), CjCas9 (0.791 vs. 0.858), enCjCas9 (0.765 vs. 0.819), Nm1Cas9 (0.780 vs. 0.876), and Nm2Cas9 (0.384 vs. 0.559; Figures S24, S25 and S26).

DeepEmbCas9 also attains high generalization performance on test sets without baselines. On mismatched G/g$N_{19}$ interfaces from Kim et al. [3], DeepEmbCas9 attains 0.538 and 0.779 Spearman correlation for xCas9 and SpCas9-NG, respectively (Figure S10 rows 2-3). Combining matched and mismatched G/g$N_{19}$ interfaces from Kim et al. [3], DeepEmbCas9 attains 0.576 and 0.797 Spearman correlation for xCas9 and SpCas9-NG, respectivley (Figure S11). On matched G/g$N_{19}$ and tRNA$^{Gln}$-$N_{20}$ interfaces from Kim, Kim et al. [4], DeepEmbCas9 has Spearman correlations 0.643, 0.522, 0.451 and 0.200 for VQR, VRER, VRQR-HF1 and QQR1, respectively (Figure S15). On matched G/g$N_{19}$ and tRNA$^{Gln}$-$N_{20}$ and mismatched G/g$N_{19}$ Sniper2P interfaces from Kim, Kim, Okafor et al. [6], DeepEmbCas9 attains 0.917 and 0.833 Spearman correlation performance, respectively (Figures S21 row 3 and S22 row 3). Combining matched and mismatched Sniper2P interfaces, DeepEmbCas9 attains 0.912 Spearman correlation (Figures S23 row 3).

#### 2.2 Leave-one-nuclease-out extrapolation performance

##### 2.2.1 Further leave-one-nuclease-out extrapolation performance

DeepEmbCas9_omit attains higher Spearman correlation than all individual activity prediction tools on 11 out of 48 benchmark test sets, namely 1 mismatched SpCas9 interface test set from Kim et al. [3] (Figure S9 row 1), 2 matched xCas9 interface test sets from Kim et al. [3] and Kim, Kim et al. [4] (Figures S9 row 2 and S14 row 1), 2 matched SpCas9-NRCH and SpCas9-NRTH interface test sets from Kim, Choi et al. [7] (Figure S19 rows 1 and 3), 1 mismatched Sniper2L interface test set from Kim, Kim, Okafor et al. [6], and and 5 small Cas9 test sets (3 SaCas9 variants, SauriCas9-KKH, and SlugCas9-HF) from Seo et al. [5] (Figures S24-S26). As for the remaining 37 test sets, DeepEmbCas9_omit has an average Spearman performance drop of $6.65 \times 10^{-2}$ compared to the best-performing individual activity prediction tools not trained on the test sets' nucleases, with the test set containing matched G/g$N_{19}$ Sc++ interfaces from Kim, Choi et al. [7] yielding the largest Spearman drop of 0.275 (DSpCv_Sniper-Cas9's 0.554 vs. DeepEmbCas9_omit's 0.279; Figure S17 row 2).

Among the test sets with extrapolation baselines and not in the 51 benchmark test sets (excluding Kim, Kim et al. [4]'s QQR1 test set), DeepEmbCas9_omit outperforms all individual activity prediction tools on 4 out of 14 test sets, namely 2 mismatched xCas9 and SpCas9-NG interface test sets from Kim et al. [3] (Figure S10 rows 2-3) and 2 (mis)matched xCas9 and SpCas9-NG interface test sets from Kim et al. [3] (Figure S11 rows 2-3). As for the remaining 10 test sets, DeepEmbCas9_omit has an average Spearman performance drop of $1.78 \times 10^{-2}$ compared to the best-performing individual activity prediction tools, with the test set containing matched VQR G/g$N_{19}$ interfaces from Kim, Kim et al. [3] yielding the largest Spearman drop of $6.09 \times 10^{-2}$ (DCv_VRQR's 0.691 vs. DeepEnsEmbCas9_naive_omit 0.630).

DeepEnsEmbCas9_omit attains higher Spearman correlation than all individual activity prediction tools on 9 out of 48 benchmark test sets, namely 2 mismatched G/g$N_{19}$ SpCas9 interface test sets (Figures S10 row 1 and S16 row 2), 1 matched eSpCas9(1.1) interface test set from Kim, Kim et al. [4] (Figure S12 row 2), 2 matched xCas9 interface test sets from Kim et al. [3] and Kim, Kim et al. [4] (Figures S9 row 2 and S14 row 1), 1 mismatched Sniper2L interface test sets from Kim, Kim,

Okafor et al. [6] (Figure S22 rows 2), and 3 small Cas9 test sets (SaCas9, SlugCas9-HF and Nm1Cas9) from Seo et al. [5] (Figures S24-S26). As for the remaining 39 test sets, DeepEnsEmbCas9_omit has an average Spearman performance drop of $5.39 \times 10^{-2}$ compared to the best-performing individual activity prediction tools not trained on the test sets' nucleases, with the test set containing (mis)matched G/gN$_{21}$ Sa-SlugCas9 interfaces from Seo et al. [5] yielding the largest Spearman drop of 0.206 (DeepSmallCas9_SauriCas9's 0.884 vs. DeepEnsEmbCas9_omit's 0.678; Figure S25 row 4).

Among the test sets with extrapolation baselines and not in the 51 benchmark test sets (excluding Kim, Kim et al. [4]'s QQR1 test set), DeepEnsEmbCas9_omit outperforms all individual activity prediction tools on 3 out of 14 test sets, namely the mismatched xCas9, (mis)matched SpCas9 and (mis)matched xCas9 interface test sets from Kim et al. [3]. As for the remaining 11 test sets, DeepEnsEmbCas9_omit has an average Spearman performance drop of 0.120 compared to the best-performing individual activity prediction tools, with the test set containing mismatched G/gN$_{19}$ SpCas9-NG interfaces from Kim et al. [3] yielding the largest Spearman drop of 0.576 (DeepSpCas9-v2's 0.603 vs. DeepEnsEmbCas9_omit's 0.0274; Figure S10 row 3).

DeepEmbCas9-MVE_omit attains higher Spearman correlation than all individual activity prediction tools on 6 out of 48 benchmark test sets, namely 1 mismatched G/gN$_{19}$ SpCas9 interface test sets from Kim et al. [3] (Figures S10 row 1), 3 matched xCas9 interface test sets (Figures S9 row 2, S14 row 1 and S18 row 1), and 2 small Cas9 test sets (SaCas9 and Nm1Cas9) from Seo et al. [5] (Figures S24 and S26). As for the remaining 42 test sets, DeepEmbCas9-MVE_omit has an average Spearman performance drop of $7.11 \times 10^{-2}$ compared to the best-performing individual activity prediction tools not trained on the test sets' nucleases, with the test set containing matched G/gN$_{19}$ SpCas9-NG interfaces from Kim et al. [3] yielding the largest Spearman drop of 0.506 (DCv_SpG's 0.786 vs. DeepEmbCas9-MVE_omit's 0.279; Figure S9 row 3).

Among the test sets with extrapolation baselines and not in the 51 benchmark test sets (excluding Kim, Kim et al. [4]'s QQR1 test set), DeepEmbCas9-MVE_omit outperforms all individual activity prediction tools on 3 out of 14 test sets, namely the mismatched xCas9, (mis)matched SpCas9 and (mis)matched xCas9 interface test sets from Kim et al. [3]. As for the remaining 11 test sets, DeepEnsEmbCas9_omit has an average Spearman performance drop of 0.120 compared to the best-performing individual activity prediction tools, with the test set containing mismatched G/gN$_{19}$ SpCas9-NG interfaces from Kim et al. [3] yielding the largest Spearman drop of 0.995 (DeepSpCas9-v2's 0.603 vs. DeepEnsEmbCas9_omit's -0.392; Figure S10 row 3).

##### 2.2.2 DeepEmbCas9 extrapolates to unseen Cas9 variants

Deep(Ens)EmbCas9_omit has decent leave-Cas9-nuclease-out performance compared to benchmark test sets (Figure S7-S26). Similar trends are observed when using the entire dataset for train-test splits (Figure S27 for Spearman, Figure S28 for Pearson).

On matched A/GN$_{19}$ SpCas9 interfaces from Wang et al. [2], DeepEmbCas9_omit (0.677) attains higher Spearman correlation than all non-SpCas9 activity tools except for DeepHF_eSpCas9(1.1) (0.707) and DeepHF_SpCas9-HF1 (0.690; Figure S7 row 1). On matched A/GN$_{19}$ SpCas9-HF1 interfaces from Wang et al. [2], DeepEmbCas9_omit (0.671) attains higher Spearman correlation than all non-SpCas9 activity tools except for DeepHF_eSpCas9(1.1) (0.769) and DeepHF_SpCas9 (GitHub) (0.680; Figure S7 row 2). On matched A/GN$_{19}$ eSpCas9(1.1) interfaces from Wang et al. [2], DeepEmbCas9_omit (0.725) attains higher Spearman correlation than all non-SpCas9 activity tools except for DeepHF_SpCas9-HF1 (0.777) and DeepHF_SpCas9 (GitHub) (0.730; Figure S7 row 3).

On matched G/gN$_{19}$ SpCas9 interfaces from Kim et al. [1], DeepEmbCas9_omit (0.601) attains higher Spearman correlation than all non-SpCas9 activity tools apart from DeepHF_eSpCas9(1.1) (0.693), DeepHF_SpCas9-HF1 (0.674) and DeepxCas9 (0.611; Figure S8).

On matched G/gN$_{19}$ SpCas9 interfaces from Kim et al. [3], DeepEmbCas9_omit (0.859) attains higher Spearman correlation than all non-SpCas9 activity tools apart from DeepSniper's Sniper1_on (0.909), DeepSniper's Sniper2L_on (0.905), and DeepSpCas9variants's Sniper-Cas9 model (0.900; Figure S9 row 1). On matched G/gN$_{19}$ xCas9 interfaces from Kim et al. [3], DeepEmbCas9_omit (0.850) attains higher Spearman correlation than all non-xCas9 activity tools (Figure S9 row 1). On matched G/gN$_{19}$ SpCas9-NG interfaces from Kim et al. [3], DeepEmbCas9_omit (0.765) attains higher Spearman correlation than all non-SpCas9-NG activity tools except for DeepCas9variants's SpG model (0.786) and DeepxCas9 (0.772; Figure S9 row 3).

On mismatched G/gN$_{19}$ SpCas9 interfaces from Kim et al. [3], DeepEmbCas9_omit (0.528) attains higher Spearman correlation than DeepSniper's Sniper1 (0.295) and Sniper2L (0.250) models (Figure S10 row 1). On mismatched G/gN$_{19}$ xCas9 interfaces from Kim et al. [3], DeepEmbCas9_omit (0.558) attains higher Spearman correlation than DeepSpCas9-v2 (0.525), DeepSniper's Sniper1 model (0.368) and DeepSniper's Sniper2L model (0.335; Figure S10 row 2). On mismatched G/gN$_{19}$ SpCas9-NG interfaces from Kim et al. [3], DeepEmbCas9_omit (0.745) attains higher Spearman correlation than DeepSpCas9-v2 (0.603), DeepSniper's Sniper1 model (0.461) and DeepSniper's Sniper2L model (0.413; Figure S10 row 3).

Combining matched and mismatched SpCas9 interfaces from Kim et al. [3], DeepEmbCas9_omit (0.823) has lower Spearman correlation than DeepSniper's Sniper1 (0.846) and Sniper2L (0.837) models (Figure S11 row 1). As for (mis)matched xCas9 interfaces from Kim et al.[3], DeepEmbCas9_omit (0.594) attain higher Spearman correlation than DeepSpCas9-v2 (0.550), DeepSniper's Sniper1 model (0.404) and DeepSniper's Sniper2L model (0.379; Figure S11 row 2). Such is also the case for SpCas9-NG interfaces from Kim et al.[3], with DeepEmbCas9_omit (0.7580) surpassing DeepSpCas9-v2 (0.612), DeepSniper's Sniper1 model (0.479) and DeepSniper's Sniper2L model (0.441; Figure S11 row 3).

On matched G/gN$_{19}$ and tRNA$^{Gln}$-N$_{20}$ SpCas9 interfaces from Kim, Kim et al. [4], DeepEmbCas9_omit (0.903) attains higher Spearman correlation than all non-SpCas9 models except for DeepSniper's Sniper1_on model (0.935), DeepSpCas9variants's Sniper-Cas9 model (0.931) and DeepSniper's Sniper2L_on model (0.931; Figure S12 row 1). On matched G/gN$_{19}$ and tRNA$^{Gln}$-N$_{20}$ eSpCas9(1.1) interfaces from Kim, Kim et al. [4], DeepEmbCas9_omit (0.850) attains higher Spearman correlation than all non-eSpCas9(1.1) models except for DeepSniper's Sniper2L_on model (0.858) and DeepSpCas9variants's Sniper-Cas9 model (0.852; Figure S12 row 2). On matched G/gN$_{19}$ and tRNA$^{Gln}$-N$_{20}$ SpCas9-HF1 interfaces from Kim, Kim et al. [4], DeepEmbCas9_omit (0.767) attains higher Spearman correlation than all non-SpCas9-HF1 models except for DSpCv_eSpCas9(1.1) (0.829), DSpCv_HypaCas9 (0.814), DS_Sniper2L_on (0.798), DSpCv_Sniper-Cas9 (0.788), DS_Sniper1_on (0.780) and DCv_SpCas9 (0.776; Figure S12 row 3).

On matched G/gN$_{19}$ and tRNA$^{Gln}$-N$_{20}$ HypaCas9 interfaces from Kim, Kim et al. [4], DeepEmbCas9_omit (0.833) attains higher Spearman correlation than all non-HypaCas9 models except for DeepSniper's Sniper2L_on model (0.839; Figure S13 row 1). On matched G/gN$_{19}$ and tRNA$^{Gln}$-N$_{20}$ evoCas9 interfaces from Kim, Kim et al. [4], DeepEmbCas9_omit (0.576) attains higher Spearman correlation than all non-evoCas9 models except DSpCv_eSpCas9(1.1) (0.628), DSpCv_SpCas9-HF1 (0.606), DSpCv_HypaCas9 (0.590), DeepHF_eSpCas9(1.1) (0.582), DeepHF_SpCas9-HF1 (0.581) and DS_Sniper2L_on (0.576; Figure S13 row 2). On matched G/gN$_{19}$ and tRNA$^{Gln}$-N$_{20}$ Sniper-Cas9 interfaces from Kim, Kim et al. [4], DeepEmbCas9_omit (0.920) attains higher Spearman correlation than all non-Sniper-Cas9 models except for DCv_SpCas9 (0.935), DSpCv_SpCas9 (0.935) and DS_Sniper2L_on (0.934; Figure S13 row 3).

On matched G/gN$_{19}$ and tRNA$^{Gln}$-N$_{20}$ xCas9 interfaces from Kim, Kim et al. [4], DeepEmbCas9_omit (0.829) surpasses all non-xCas9 models in Spearman correlation (Figure S14 row 1). On matched G/gN$_{19}$ and tRNA$^{Gln}$-N$_{20}$ SpCas9-NG interfaces from Kim, Kim et al. [4], DeepEmbCas9_omit (0.805) attains higher Spearman correlation than all non-SpCas9-NG models apart from DSpCv_VRQR (0.919), DCv_SpG (0.889) and DCv_VRQR (0.810; Figure S14 row 2). On matched G/gN$_{19}$ and tRNA$^{Gln}$-N$_{20}$ VRQR interfaces from Kim, Kim et al. [4], DeepEmbCas9_omit (0.812) surpasses all non-VRQR models in Spearman correlation, except for DSpCv_SpCas9-NG (0.928) and DCv_SpG (0.861; Figure S14 row 3).

On matched G/gN$_{19}$ and tRNA$^{Gln}$-N$_{20}$ QQR1 interfaces from Kim, Kim et al. [4], DeepEmbCas9_omit attains 0.160 Spearman correlation (Figure S15 row 1). On matched G/g$N_{19}$ and $tRNA^{Gln}$-$N_{20}$ VQR interfaces from Kim, Kim et al. [4], DeepEmbCas9_omit (0.630) attains higher Spearman correlation for all non-VQR models except for DCv_VRQR (0.691), DSpCv_SpCas9-NG (0.655), DCv_SpCas9-NG (0.637) and DCv_SpG (0.635; Figure S15 row 2). On matched G/g$N_{19}$ and $tRNA^{Gln}$-$N_{20}$ VRER interfaces from Kim, Kim et al. [4], DeepEmbCas9_omit (0.512) attains higher Spearman correlation for all non-VRER model except for DCv_SpG (0.523) and DSpCv_SpCas9-NG (0.515; Figure S15 row 3). On matched G/g$N_{19}$ and $tRNA^{Gln}$-$N_{20}$ VRQR-HF1 interfaces from Kim, Kim et al. [4], DeepEmbCas9_omit (0.421) surpasses all non-VRQR-HF1 models in Spearman correlation, apart from DSpCv_SpCas9-NG (0.455), DCv_VRQR (0.455), DCv_SpG (0.442), DSpCv_HypaCas9 (0.439), DSpCv_evoCas9 (0.435) and DCv_SpCas9-NG (0.428; Figure S15 row 4).

On matched G/g$N_{19}$ SpCas9 interfaces from Seo et al. [5], DeepEmbCas9_omit (0.734) attains higher Spearman correlation than all non-SpCas9 models apart from DS_Sniper1_on (0.748), DS_Sniper2L_on (0.747) and DSpCv_Sniper-Cas9 (0.747; Figure S16 row 1). On mismatched G/g$N_{19}$ SpCas9 interfaces from Seo et al. [5], DeepEmbCas9_omit (0.891) attains higher Spearman correlation than DS_Sniper2L_off (0.875), but not DS_Sniper1_off (0.892; Figure S16 row 1). Combining matched and mismatched G/g$N_{19}$ SpCas9 interfaces from Seo et al. [5], DeepEmbCas9_omit (0.812) attains lower Spearman correlation than DS_Sniper1 (0.822) and DS_Sniper2L (0.822; Figure S16 row 3). On matched G/g$N_{20}$ SpCas9 interfaces, DeepEmbCas9_omit has 0.184 Spearman correlation (Figure S16 row 4).

Looking at test datasets from Kim, Choi et al. [7] with matched G/g$N_{19}$ interfaces, for SpCas9, DeepEmbCas9_omit (0.670) attains higher Spearman correlation for all non-SpCas9 models except for DSpCv_Sniper-Cas9 (0.728) and DS_Sniper2L_on (0.722; Figure S17 row 1). For Sc++, DeepEmbCas9_omit attains 0.279 Spearman correlation (Figure S17 row 2).

For xCas9, DeepEmbCas9_omit (0.656) attains higher Spearman correlation than all non-xCas9 models apart from DCv_SpCas9-NRRH (0.699), DCv_SpCas9-NRCH (0.696), DeepSpCas9-v2 (0.686), DCv_SpCas9-NRTH (0.670), DCv_SpCas9-NG (0.665), DCv_SpG (0.664), DSpCv_VRQR (0.663) and DCv_SpCas9 (0.659; Figure S18 row 1). For SpCas9-NG, DeepEmbCas9_omit (0.801) attains higher Spearman correlation than all non-SpCas9-NG models apart from DSpCv_VRQR (0.927) and DCv_SpG (0.867; Figure S18 row 2). For VRQR, DeepEmbCas9_omit (0.668) attains higher Spearman correlation than all non-VRQR models apart from DSpCv_SpCas9-NG (0.758), DCv_SpCas9-NG (0.735), DCv_SpG (0.733) and DeepSpCas9-NG (0.687; Figure S18 row 3).

For SpCas9-NRCH/SpCas9-NRTH, DeepEmbCas9_omit (0.801/0.854) attains higher Spearman correlation than all non-SpCas9-NRCH/non-SpCas9-NRTH models (Figure S19 rows 1 and 3). For SpCas9-NRRH, DeepEmbCas9_omit (0.840) attains higher Spearman correlation than all non-SpCas9-NRRH models except for DCv_SpCas9-NRTH (0.849; Figure S19 row 2).

For SpG, DeepEmbCas9_omit (0.736) attains higher Spearman correlation than all non-SpG models except for DCv_SpCas9-NG (0.870), DSpCv_VRQR (0.852), DeepSpCas9-NG (0.820), DCv_VRQR (0.779), DeepxCas9 (0.741; Figure S20 row 1). For SpRY, DeepEmbCas9_omit (0.729) attains higher Spearman correlation than all non-SpRY models except for DCv_SpCas9-NG (0.770) and DCv_SpG (0.747; Figure S20 row 2).

We next look at test datasets from Kim, Kim, Okafor et al. [6]. On matched G/g$N_{19}$ and $tRNA^{Gln}$-$N_{20}$ interfaces, for Sniper-Cas9, DeepEmbCas9_omit (0.918) attains higher Spearman correlation than all non-Sniper-Cas9 models except for DSpCv_SpCas9 (0.933), DS_Sniper2L_on (0.933) and DCv_SpCas9 (0.930; Figure S21 row 1). For Sniper2L, DeepEmbCas9_omit (0.917) attains higher Spearman correlation than all non-Sniper2L models except for DS_Sniper1_on (0.928), DSpCv_Sniper-Cas9 (0.924), DSpCv_SpCas9 (0.923) and DCv_SpCas9 (0.922; Figure S21 row 2). For Sniper2P, DeepEmbCas9_omit (0.914) attains higher Spearman correlation than all non-Sniper2P models except for DS_Sniper1_on (0.929), DS_Sniper2L_on (0.925), DSpCv_SpCas9 (0.925), DSpCv_Sniper-Cas9 (0.917), and DCv_SpCas9 (0.916; Figure S21 row 3).

Regarding mismatched G/g$N_{19}$ interfaces, for Sniper-Cas9, DeepEmbCas9_omit (0.876) attains higher

Spearman correlation than DeepSpCas9-v2 (0.866), but not DS_Sniper2L_off (0.881; Figure S22 row 1). For Sniper2L, DeepEmbCas9_omit (0.872) attains higher Spearman correlation than DS_Sniper1_off (0.880) and DeepSpCas9-v2 (0.848; Figure S22 row 2). For Sniper2P, DeepEmbCas9_omit (0.862) attains higher Spearman correlation than DS_Sniper1_off (0.840) and DeepSpCas9-v2 (0.838), but not for DS_Sniper2L_off (0.865; Figure S22 row 3).

Combining matched and mismatched interfaces from Kim, Kim, Okafor et al. [6], for Sniper-Cas9, DeepEmbCas9_omit (0.917) attains higher Spearman correlation than DeepSpCas9-v2 (0.875) but not for DS_Sniper2L (0.925; Figure S23 row 1). For Sniper2L, DeepEmbCas9_omit (0.918) attains higher Spearman correlation than DeepSpCas9-v2 (0.868) but not for DS_Sniper1 (0.922; Figure S23 row 2). For Sniper2P, DeepEmbCas9_omit (0.918) attains higher Spearman correlation than DeepSpCas9-v2 (0.868) but not for DS_Sniper2L (0.924) and DS_Sniper1 (0.922; Figure S23 row 3).

We next examine small Cas9 nuclease test datasets from Seo et al. [5]. Looking at test sets with G/g$N_{21}$ interfaces, for SaCas9/efSaCas9/SaCas9-HF, DeepEmbCas9_omit (0.815/0.855/0.831) attains higher Spearman correlation than all non-SaCas9/non-efSaCas9/non-SaCas9-HF models, respectively (Figure S24 rows 1, 3 and 4). For eSaCas9, DeepEmbCas9_omit (0.852) attains higher Spearman correlation than all non-eSaCas9 models except for DeepSmallCas9_efSaCas9 (0.858; Figure S24 row 2). For SaCas9-KKH, DeepEmbCas9_omit (0.789) attains higher Spearman correlation than all non-SaCas9-KKH models except for DeepSmallCas9_SaCas9-KKH-HF (0.831; Figure S24 row 5). For SaCas9-KKH-HF, DeepEmbCas9_omit (0.833) attains higher Spearman correlation than all non-SaCas9-KKH-HF models except for DeepSmallCas9_SaCas9-KKH (0.883; Figure S24 row 6).

For sRGN3.1, DeepEmbCas9_omit (0.870) attains higher Spearman correlation than all non-sRGN3.1 models except for DeepSmallCas9_SlugCas9 (0.894) and DeepSmallCas9_SauriCas9 (0.874; Figure S25 row 1). For SlugCas9, DeepEmbCas9_omit (0.901) attains higher Spearman correlation than all non-SlugCas9 models except for DeepSmallCas9_sRGN3.1 (0.915; Figure S25 row 2). For SauriCas9, DeepEmbCas9_omit (0.846) attains higher Spearman correlation than all non-SauriCas9 models except for DeepSmallCas9_Sa-SlugCas9 (0.893), DeepSmallCas9_SauriCas9-KKH (0.862) and DeepSmallCas9_sRGN3.1 (0.848; Figure S25 row 3). For Sa-SlugCas9, DeepEmbCas9_omit (0.688) attains lower Spearman correlation than all non-Sa-SlugCas9 models (Figure S25 rows 4-6). For SauriCas9-KKH/SlugCas9-HF, DeepEmbCas9_omit (0.817/0.783) attains higher Spearman correlation than all non-SauriCas9-KKH/non-SlugCas9-HF models (Figure S25 rows 4-6).

Looking at non-G/g$N_{21}$ nucleases, for G/g$N_{19}$ St1Cas9 interfaces, DeepEmbCas9_omit attains 0.068 Spearman correlation (Figure S26 row 1). For G/g$N_{22}$ CjCas9 interfaces, DeepEmbCas9_omit (0.708) attains lower Spearman correlation than DeepSmallCas9_enCjCas9 (0.820; Figure S26 row 2). For G/g$N_{22}$ enCjCas9 interfaces, DeepEmbCas9_omit (0.741) attains lower Spearman correlation than DeepSmallCas9_CjCas9 (0.887; Figure S26 row 3). For G/g$N_{23}$ Nm1Cas9 interfaces, DeepEmbCas9_omit (-0.049) attains lower Spearman correlation than DeepSmallCas9_Nm2Cas9 (0.193; Figure S26 row 4). For G/g$N_{22}$ and G/g$N_{23}$ Nm2Cas9 interfaces, DeepEmbCas9_omit attains -0.007 Spearman correlation (Figure S26 row 5).

Performance is varied when leaving one gRNA scaffold out for testing (Figures S29 and Figure S30 for Spearman and Pearson correlations, respectively).

#### 2.3 Per-nuclease in-distribution and extrapolation plots

##### 2.3.1 SpCas9 variants and Sc++

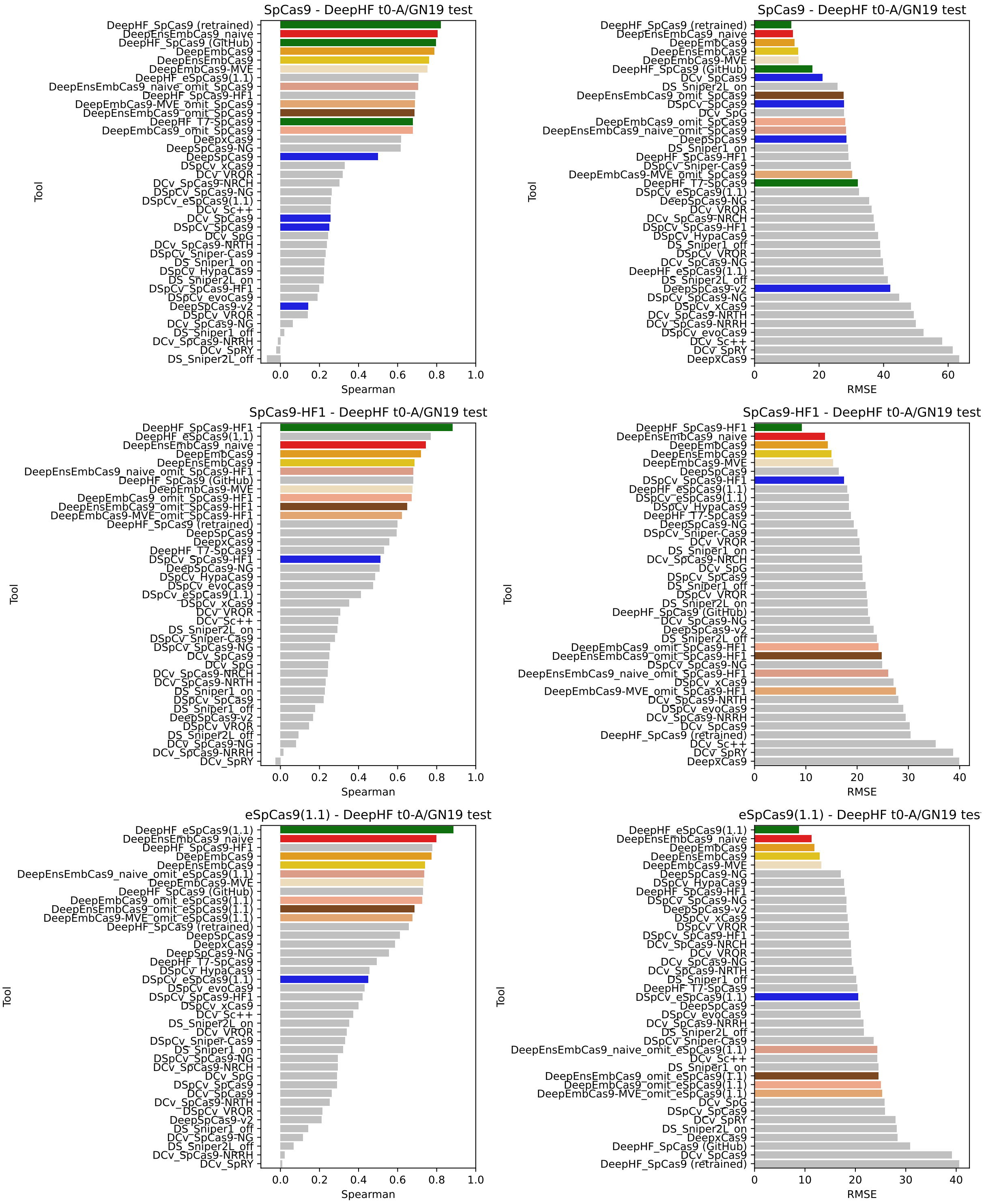


Figure S7: Benchmark test Spearman correlation (left) and RMSE (right) comparisons for DeepEmbCas9 (orange), DeepEnsEmbCas9_naive (red), DeepEmbCas9-MVE (wheat-colored), DeepEnsEmbCas9 (gold), DeepEmbCas9_omit (dark salmon) and DeepEnsEmbCas9_omit (brown) against relevant individual Cas9 cleavage activity tools for matched A/GN$_{19}$ wild type SpCas9 (top), SpCas9-HF1 (middle) and eSpCas9(1.1) (bottom) interfaces from Wang et al. [2], where green and blue bars denote DeepHF and other individual Cas9 cleavage activity tools trained on matched interfaces of the test nuclease, respectively.

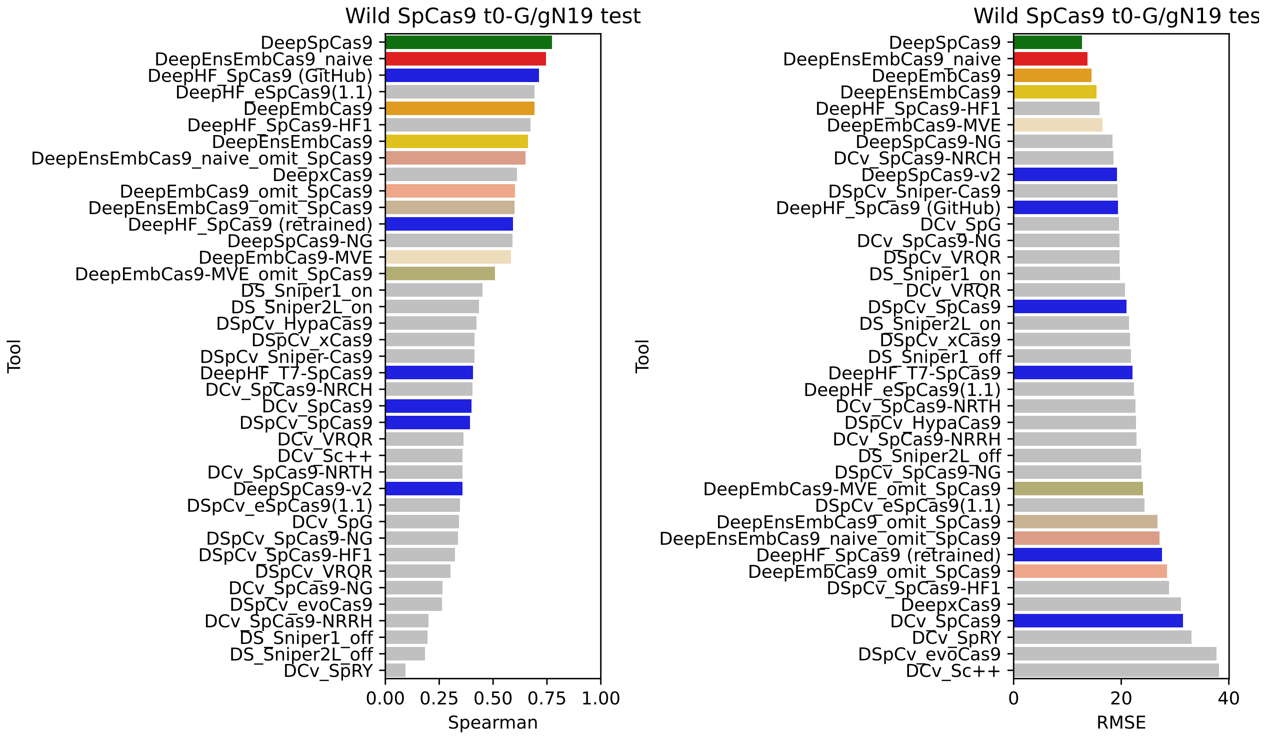


Figure S8: Benchmark test Spearman correlation (left) and RMSE (right) comparisons for DeepEmbCas9 (orange), DeepEnsEmbCas9_naive (red), DeepEmbCas9-MVE (wheat-colored), DeepEnsEmbCas9 (gold), DeepEmbCas9_omit (dark salmon) and DeepEnsEmbCas9_omit (brown) against relevant individual Cas9 cleavage activity tools for matched G/gN$_{19}$ wild type SpCas9 interfaces from Kim, Kim et al. [1], where green and blue bars denote DeepSpCas9 and other individual Cas9 cleavage activity tools trained on matched wild type SpCas9 interfaces, respectively.

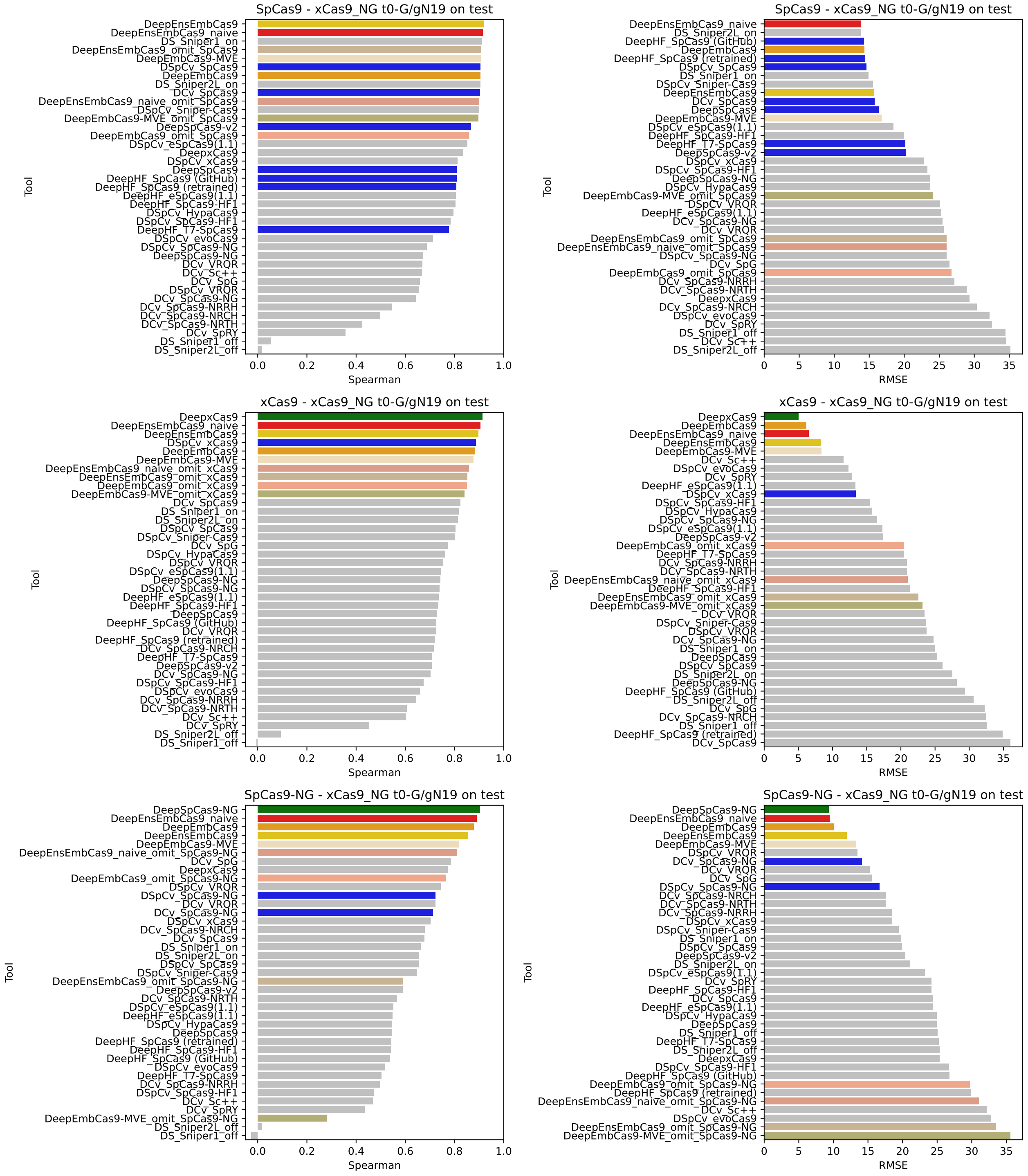

Figure S9: Benchmark test Spearman correlation (left) and RMSE (right) comparisons for DeepEmbCas9 (orange), DeepEnsEmbCas9_naive (red), DeepEmbCas9-MVE (wheat-colored), DeepEnsEmbCas9 (gold), DeepEmbCas9_omit (dark salmon) and DeepEnsEmbCas9_omit (brown) against relevant individual Cas9 cleavage activity tools for matched G/gN$_{19}$ wild type SpCas9 (top), xCas9 (middle) and SpCas9-NG (bottom) interfaces from Kim et al. [3], where green bars denote DeepxCas9 (middle row) and DeepSpCas9-NG (bottom row), and blue bars denote other individual Cas9 cleavage activity tools trained on matched interfaces of the test nuclease.

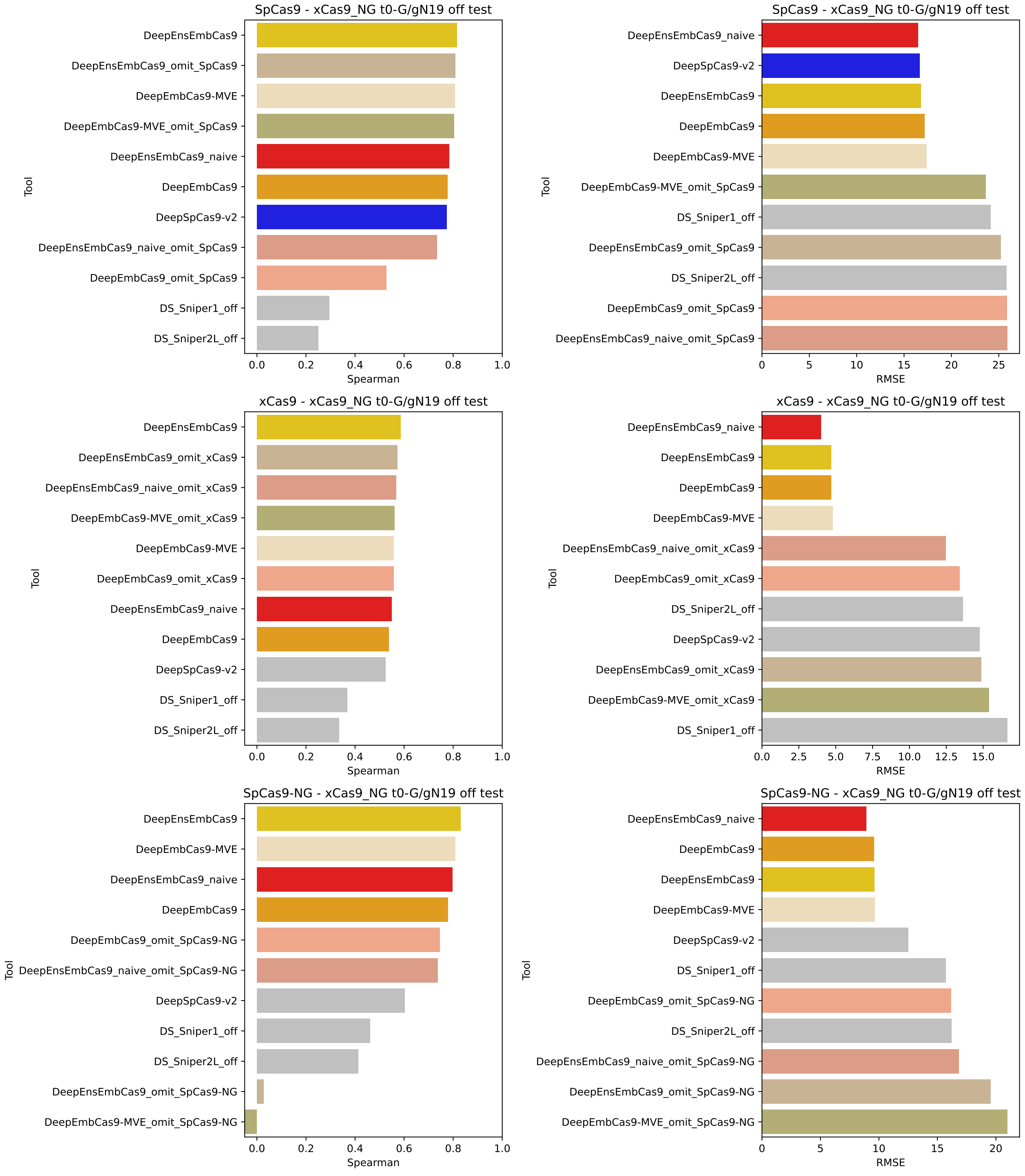


Figure S10: Benchmark test Spearman correlation (left) and RMSE (right) comparisons for DeepEmbCas9 (orange), DeepEnsEmbCas9_naive (red), DeepEmbCas9-MVE (wheat-colored), DeepEnsEmbCas9 (gold), DeepEmbCas9_omit (dark salmon) and DeepEnsEmbCas9_omit (brown) against relevant individual Cas9 cleavage activity tools for mismatched G/g$N_{19}$ wild type SpCas9 (top), SpCas9-NG (middle) and xCas9 (bottom) interfaces from Kim et al. [3], where blue bars denote individual Cas9 cleavage activity tools trained on mismatched interfaces of the test nuclease.

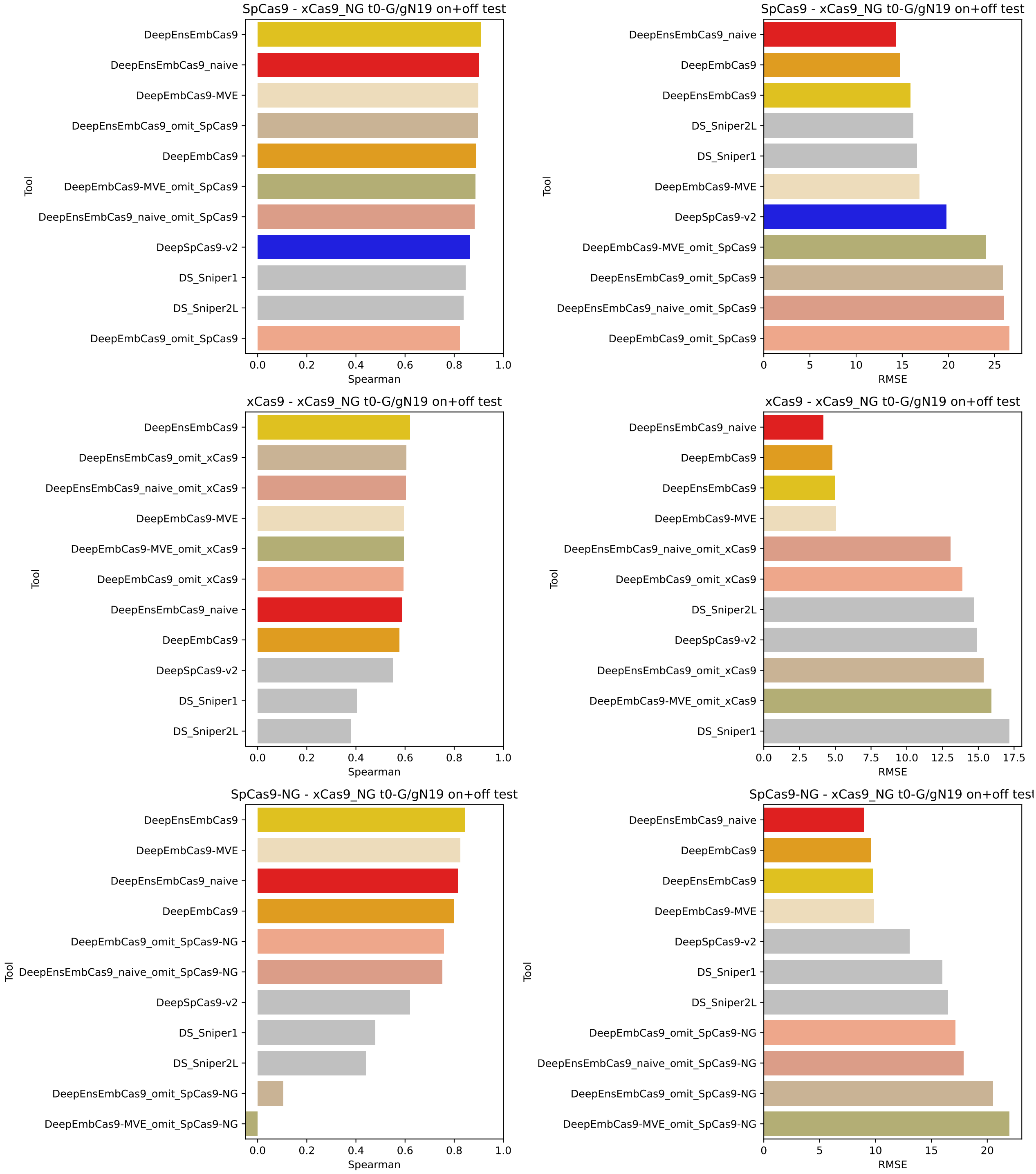


Figure S11: Benchmark test Spearman correlation (left) and RMSE (right) comparisons for DeepEmbCas9 (orange), DeepEnsEmbCas9_naive (red), DeepEmbCas9-MVE (wheat-colored), DeepEnsEmbCas9 (gold), DeepEmbCas9_omit (dark salmon) and DeepEnsEmbCas9_omit (brown) against relevant individual Cas9 cleavage activity tools for (mis)matched G/g$N_{19}$ wild type SpCas9 (top), SpCas9-NG (middle) and xCas9 (bottom) interfaces from Kim et al. [3], where blue bars denote individual Cas9 cleavage activity tools trained on (mis)matched interfaces of the test nuclease.

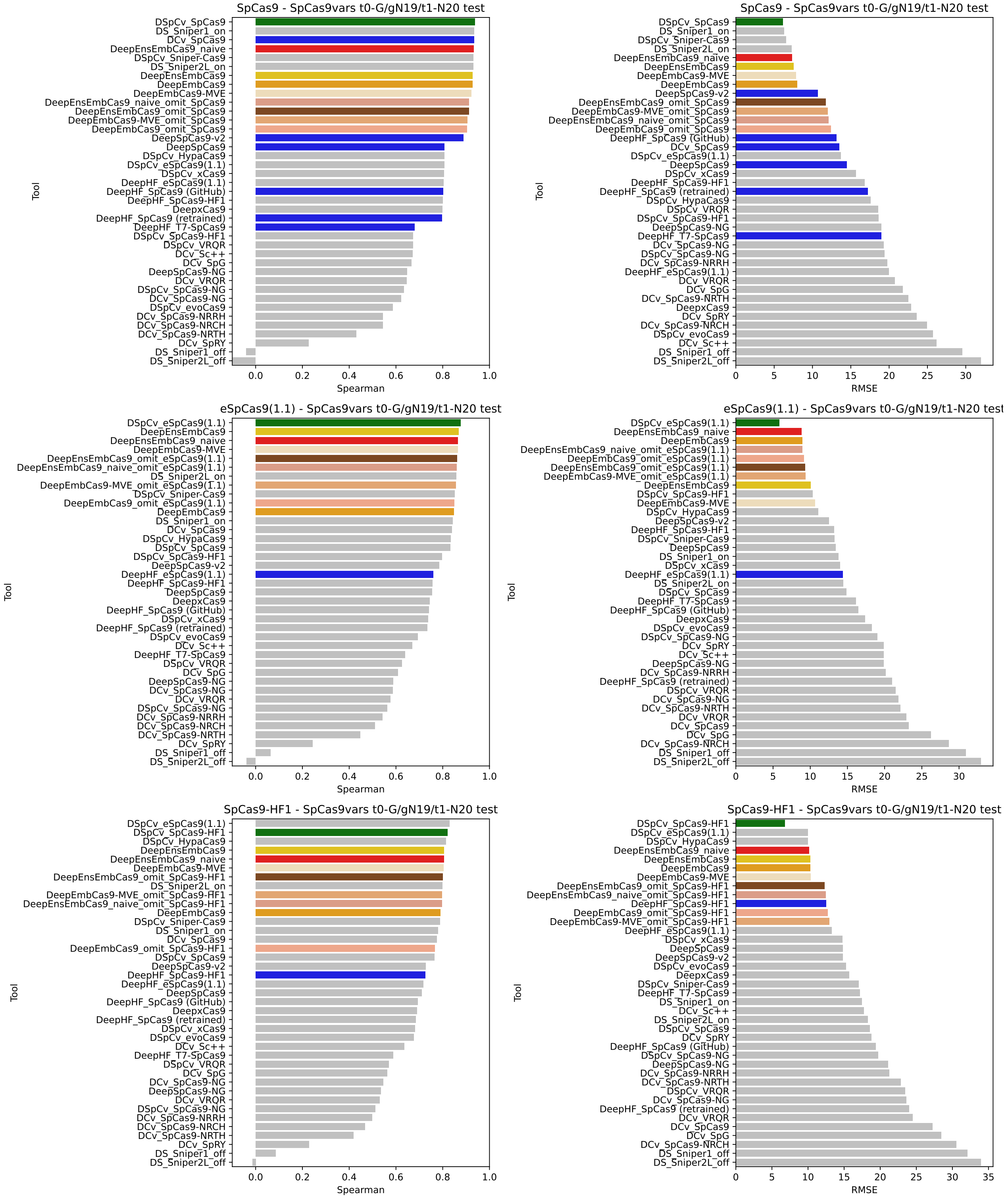


Figure S12: Benchmark test Spearman correlation (left) and RMSE (right) comparisons for DeepEmbCas9 (orange), DeepEnsEmbCas9_naive (red), DeepEmbCas9-MVE (wheat-colored), DeepEnsEmbCas9 (gold), DeepEmbCas9_omit (dark salmon) and DeepEnsEmbCas9_omit (brown) against relevant individual Cas9 cleavage activity tools for matched G/g$N_{19}$ and tRNA$^{Gln}$-$N_{20}$ wild type SpCas9 (top), eSpCas9(1.1) (middle) and SpCas9-HF1 (bottom) interfaces from Kim, Kim et al. [4], where green bars denote DeepSpCas9variants, and blue bars denote other individual Cas9 cleavage activity tools trained on matched interfaces of the test nuclease.

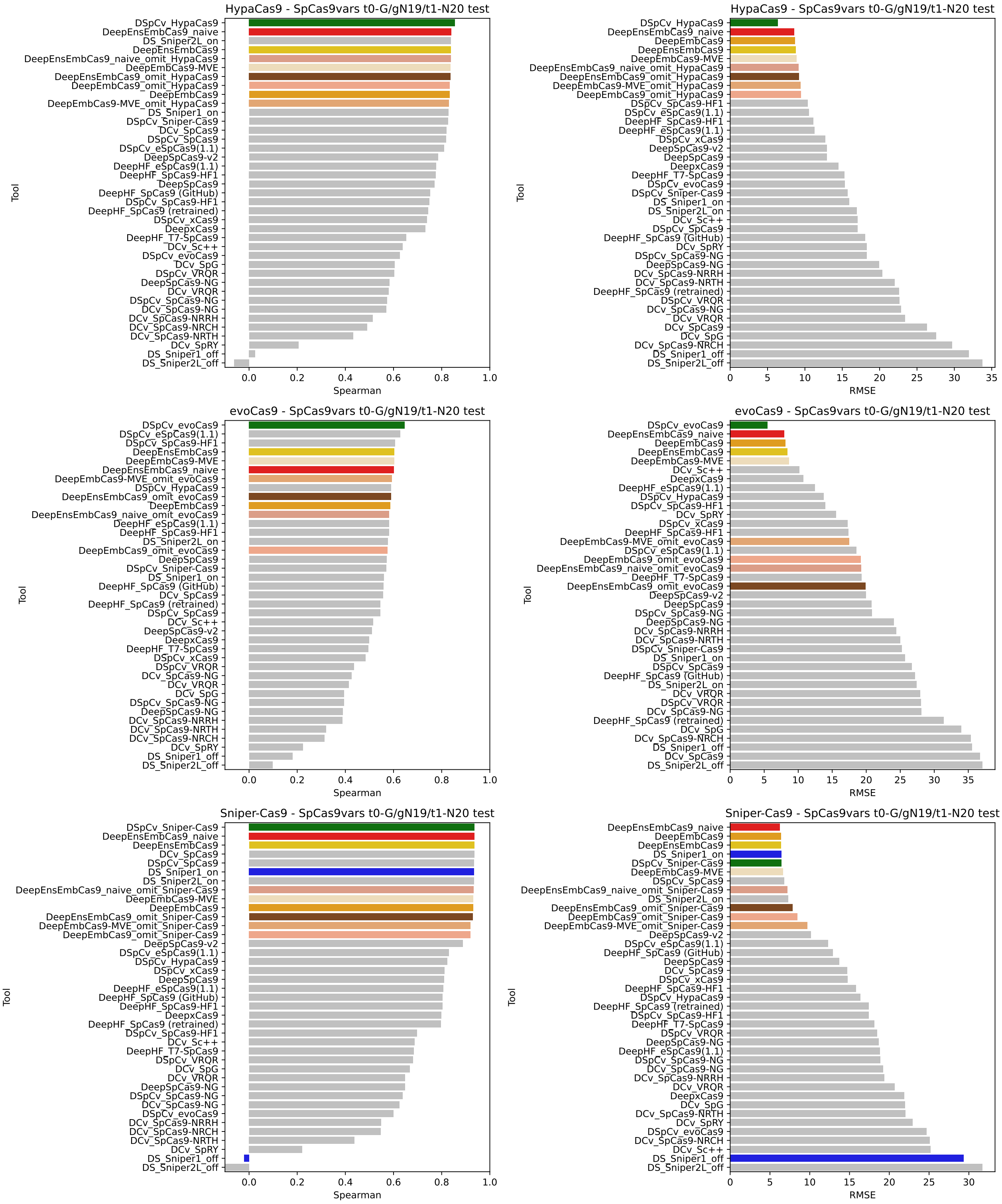


Figure S13: Benchmark test Spearman correlation (left) and RMSE (right) comparisons for DeepEmbCas9 (orange), DeepEnsEmbCas9_naive (red), DeepEmbCas9-MVE (wheat-colored), DeepEnsEmbCas9 (gold), DeepEmbCas9_omit (dark salmon) and DeepEnsEmbCas9_omit (brown) against relevant individual Cas9 cleavage activity tools for matched G/g$N_{19}$ and tRNA$^{Gln}$-$N_{20}$ HypaCas9 (top), evoCas9 (middle) and Sniper-Cas9 (bottom) interfaces from Kim, Kim et al. [4], where green bars denote DeepSpCas9variants, and blue bars denote other individual Cas9 cleavage activity tools trained on matched interfaces of the test nuclease.

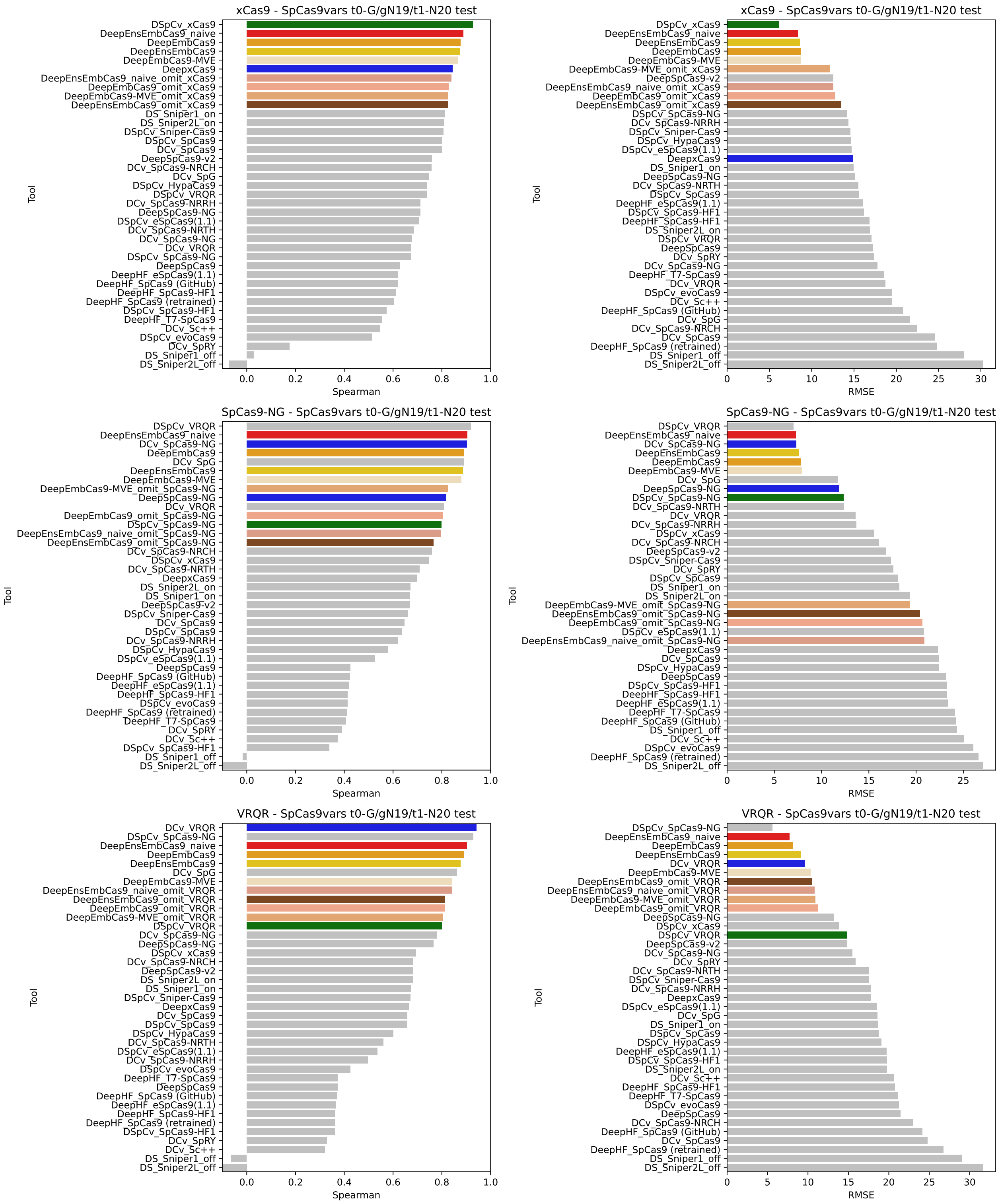


Figure S14: Benchmark test Spearman correlation (left) and RMSE (right) comparisons for DeepEmbCas9 (orange), DeepEnsEmbCas9_naive (red), DeepEmbCas9-MVE (wheat-colored), DeepEnsEmbCas9 (gold), DeepEmbCas9_omit (dark salmon) and DeepEnsEmbCas9_omit (brown) against relevant individual Cas9 cleavage activity tools for matched G/gN$_{19}$ and tRNA$^{\mathrm{Gln}}$-N$_{20}$ xCas9 (top), SpCas9-NG (middle) and VRQR (bottom) interfaces from Kim, Kim et al. [4], where green bars denote DeepSpCas9variants, and blue bars denote other individual Cas9 cleavage activity tools trained on matched interfaces of the test nuclease.

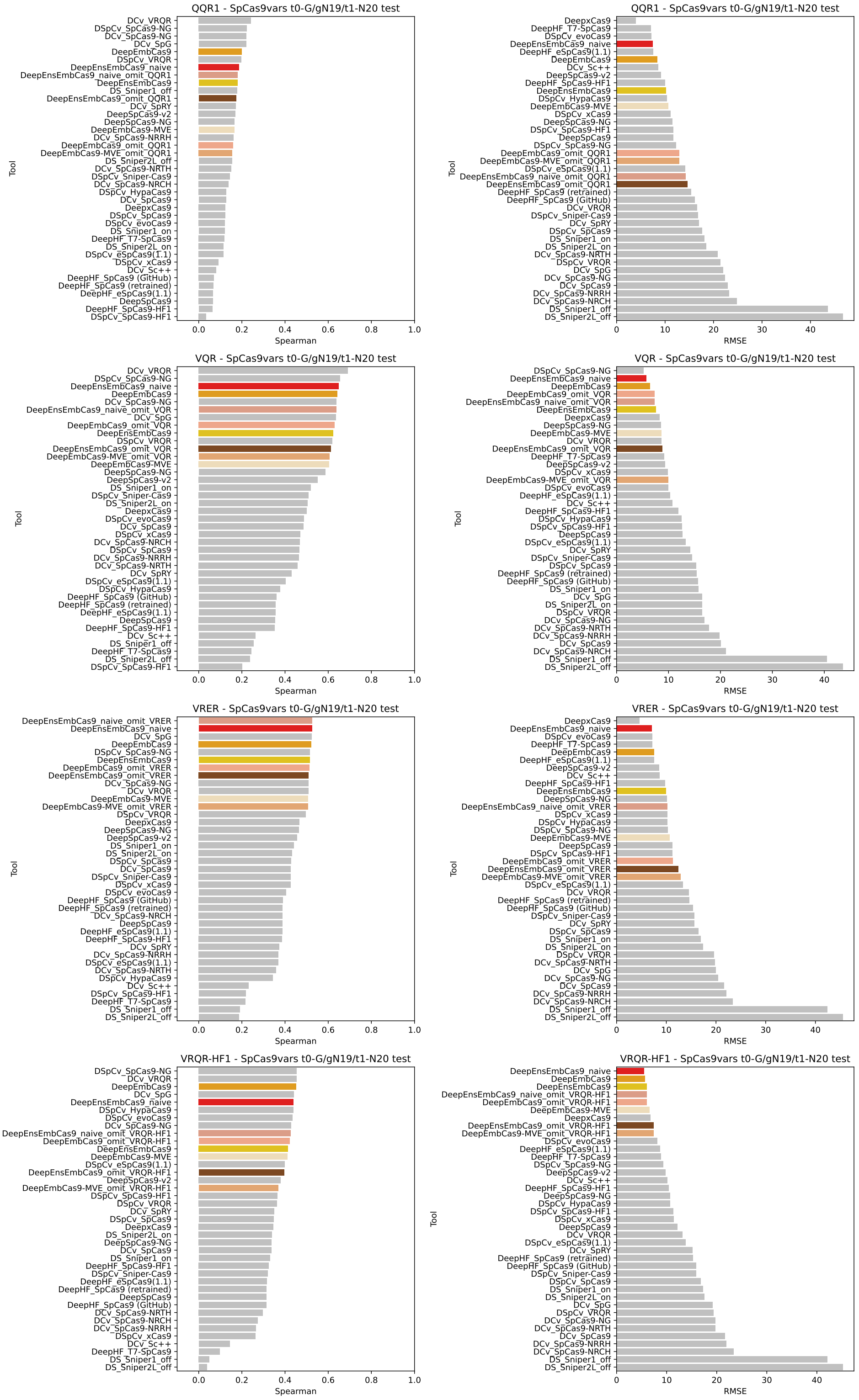


Figure S15: Benchmark test Spearman correlation (left) and RMSE (right) comparisons for DeepEmbCas9 (orange), DeepEnsEmbCas9_naive (red), DeepEmbCas9-MVE (wheat-colored), DeepEnsEmbCas9 (gold), DeepEmbCas9_omit (dark salmon) and DeepEnsEmbCas9_omit (brown) against relevant individual Cas9 cleavage activity tools for matched G/g$N_{19}$ and tRNA$^{Gln}$-$N_{20}$ QQR1 (row 1), VQR (row 2), VRER (row 3) and VRQR-HF1 (row 4) interfaces from Kim, Kim et al. [4], where green bars denote DeepSpCas9variants.

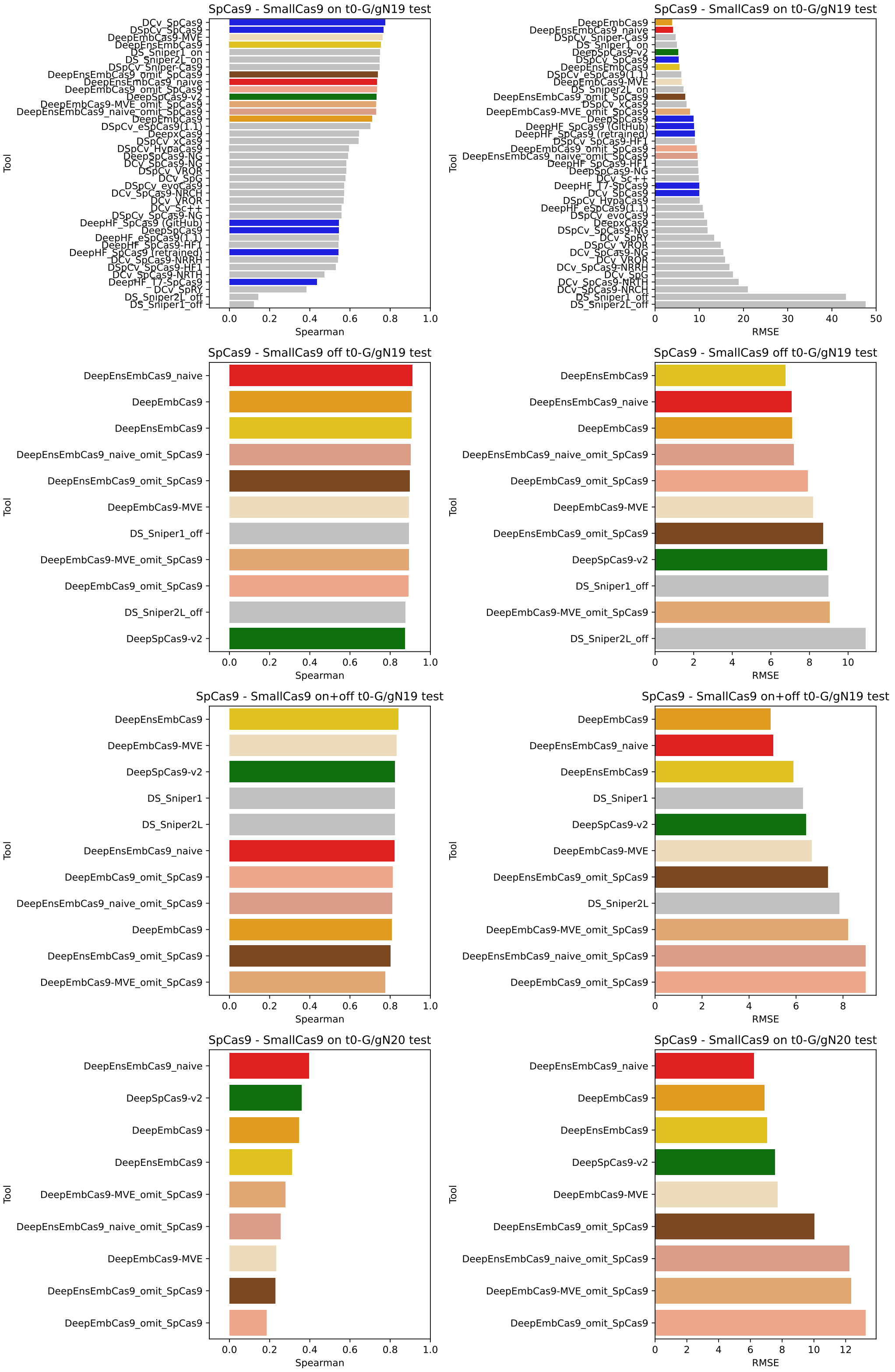


Figure S16: Benchmark test Spearman correlation (left) and RMSE (right) comparisons for DeepEmbCas9 (orange), DeepEnsEmbCas9_naive (red), DeepEmbCas9-MVE (wheat-colored), DeepEnsEmbCas9 (gold), DeepEmbCas9_omit (dark salmon) and DeepEnsEmbCas9_omit (brown) against relevant individual Cas9 cleavage activity tools for wild type SpCas9 (specifically NLS-SpCas9-NLS-FLAG-P2A) with matched G/g$N_{19}$ (row 1), mismatched G/g$N_{19}$ (row 2), (mis)matched G/g$N_{19}$ (row 3) and matched G/gN20 interfaces (row 4) from Seo et al. [5], where green bars denote DeepSpCas9-v2, and blue bars denote other individual Cas9 cleavage activity tools trained on matched interfaces of the test nuclease.

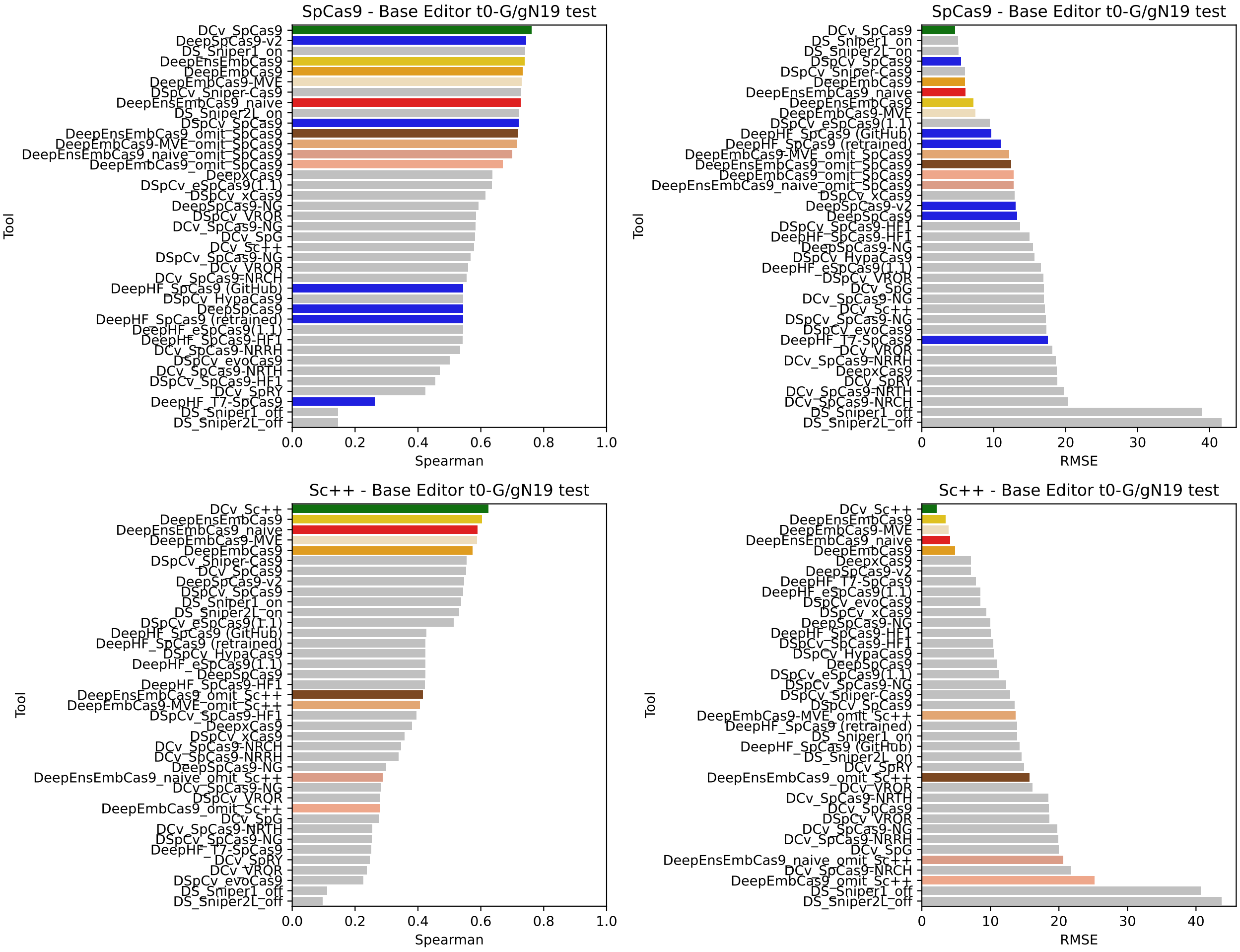


Figure S17: Benchmark test Spearman correlation (left) and RMSE (right) comparisons for DeepEmbCas9 (orange), DeepEnsEmbCas9_naive (red), DeepEmbCas9-MVE (wheat-colored), DeepEnsEmbCas9 (gold), DeepEmbCas9_omit (dark salmon) and DeepEnsEmbCas9_omit (brown) against relevant individual Cas9 cleavage activity tools for matched G/g$N_{19}$ SpCas9 (top) and Sc++ (bottom) interfaces from Kim, Choi et al. [7], where green bars denote DeepCas9variants, and blue bars denote other individual Cas9 cleavage activity tools trained on matched interfaces of the test nuclease.

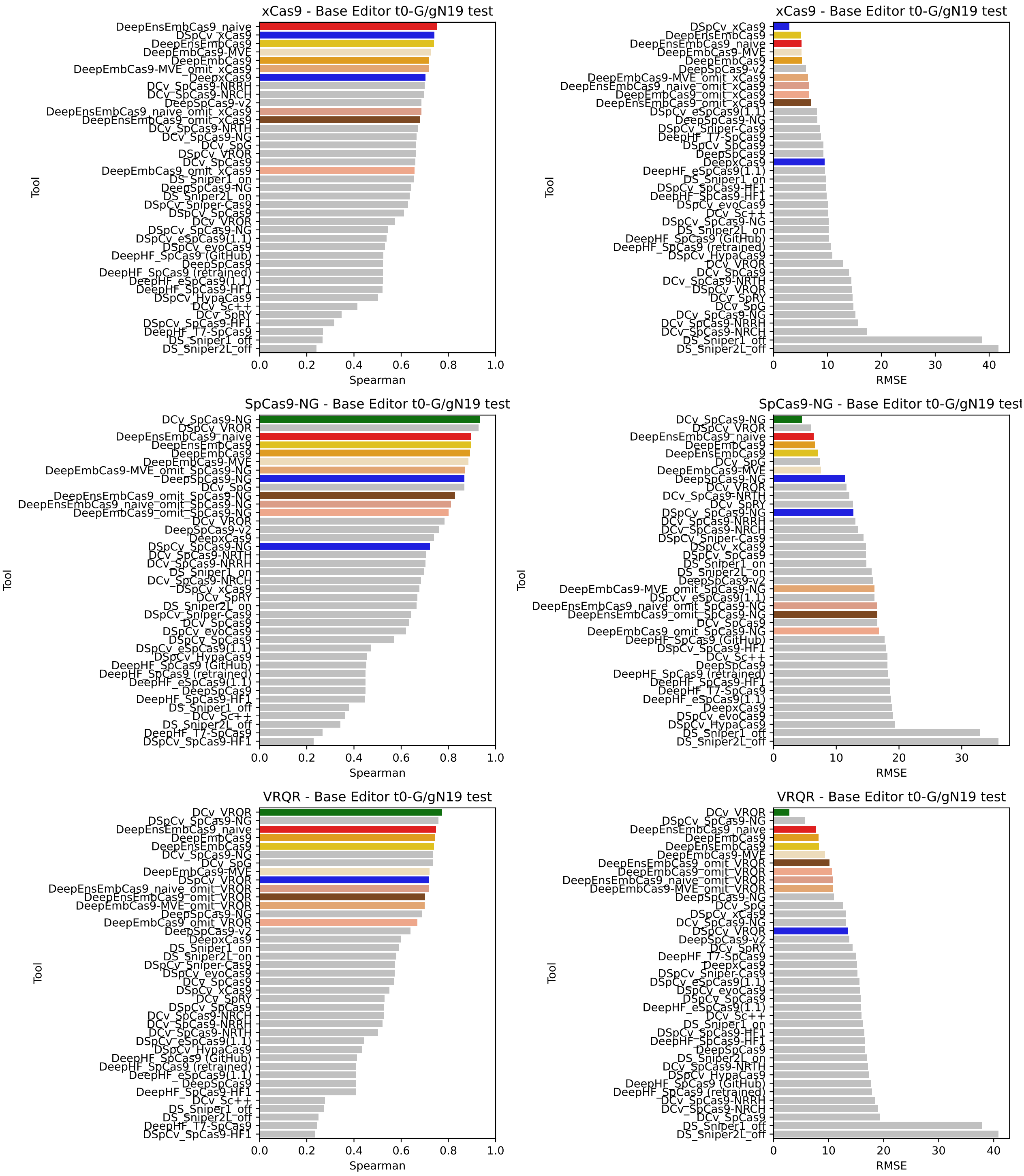


Figure S18: Benchmark test Spearman correlation (left) and RMSE (right) comparisons for DeepEmbCas9 (orange), DeepEnsEmbCas9_naive (red), DeepEmbCas9-MVE (wheat-colored), DeepEnsEmbCas9 (gold), DeepEmbCas9_omit (dark salmon) and DeepEnsEmbCas9_omit (brown) against relevant individual Cas9 cleavage activity tools for matched G/g$N_{19}$ xCas9 (row 1), SpCas9-NG (row 2) and VRQR (row 3) interfaces from Kim, Choi et al. [7], where green bars denote DeepCas9variants, and blue bars denote other individual Cas9 cleavage activity tools trained on matched interfaces of the test nuclease.

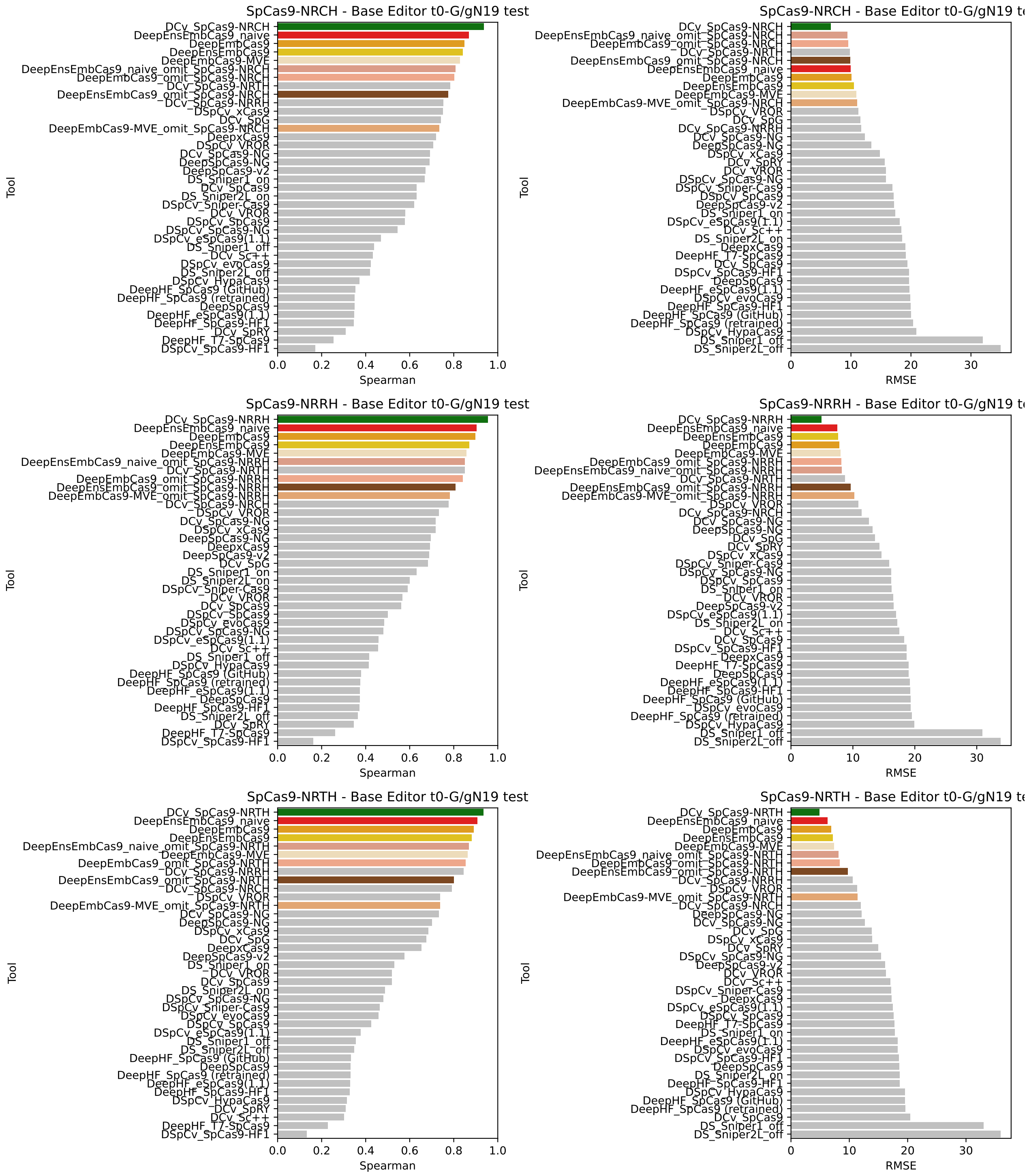


Figure S19: Benchmark test Spearman correlation (left) and RMSE (right) comparisons for DeepEmbCas9 (orange), DeepEnsEmbCas9_naive (red), DeepEmbCas9-MVE (wheat-colored), DeepEnsEmbCas9 (gold), DeepEmbCas9_omit (dark salmon) and DeepEnsEmbCas9_omit (brown) against relevant individual Cas9 cleavage activity tools for matched G/gN$_{19}$ SpCas9-NRCH (row 1), SpCas9-NRRH (row 2) and SpCas9-NRTH (row 3) interfaces from Kim, Choi et al. [7], where green bars denote DeepCas9variants, and blue bars denote other individual Cas9 cleavage activity tools trained on matched interfaces of the test nuclease.

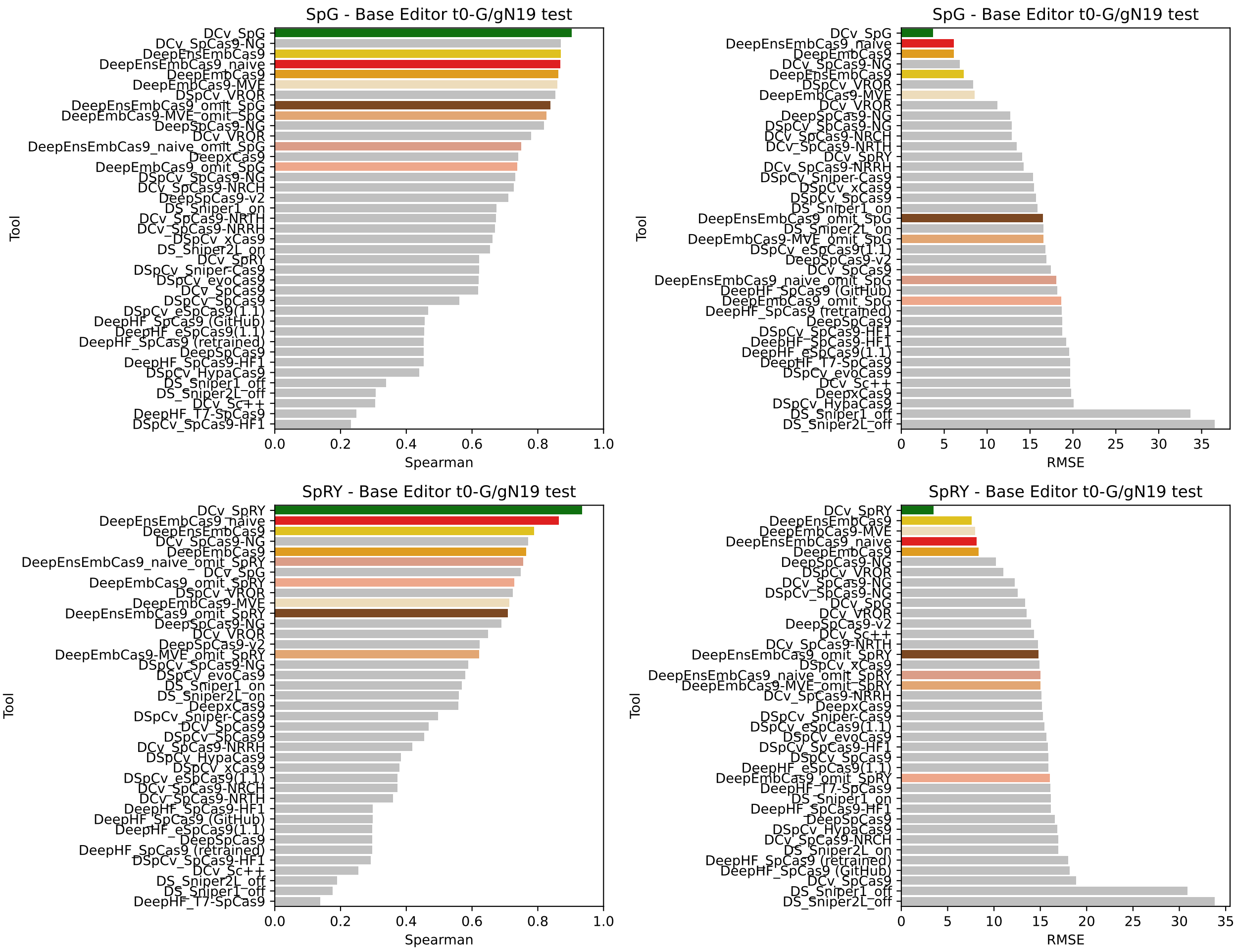


Figure S20: Benchmark test Spearman correlation (left) and RMSE (right) comparisons for DeepEmbCas9 (orange), DeepEnsEmbCas9_naive (red), DeepEmbCas9-MVE (wheat-colored), DeepEnsEmbCas9 (gold), DeepEmbCas9_omit (dark salmon) and DeepEnsEmbCas9_omit (brown) against relevant individual Cas9 cleavage activity tools for matched G/g$N_{19}$ SpG (top) and SpRY (bottom) interfaces from Kim, Choi et al. [7], where green bars denote DeepCas9variants, and blue bars denote other individual Cas9 cleavage activity tools trained on matched interfaces of the test nuclease.

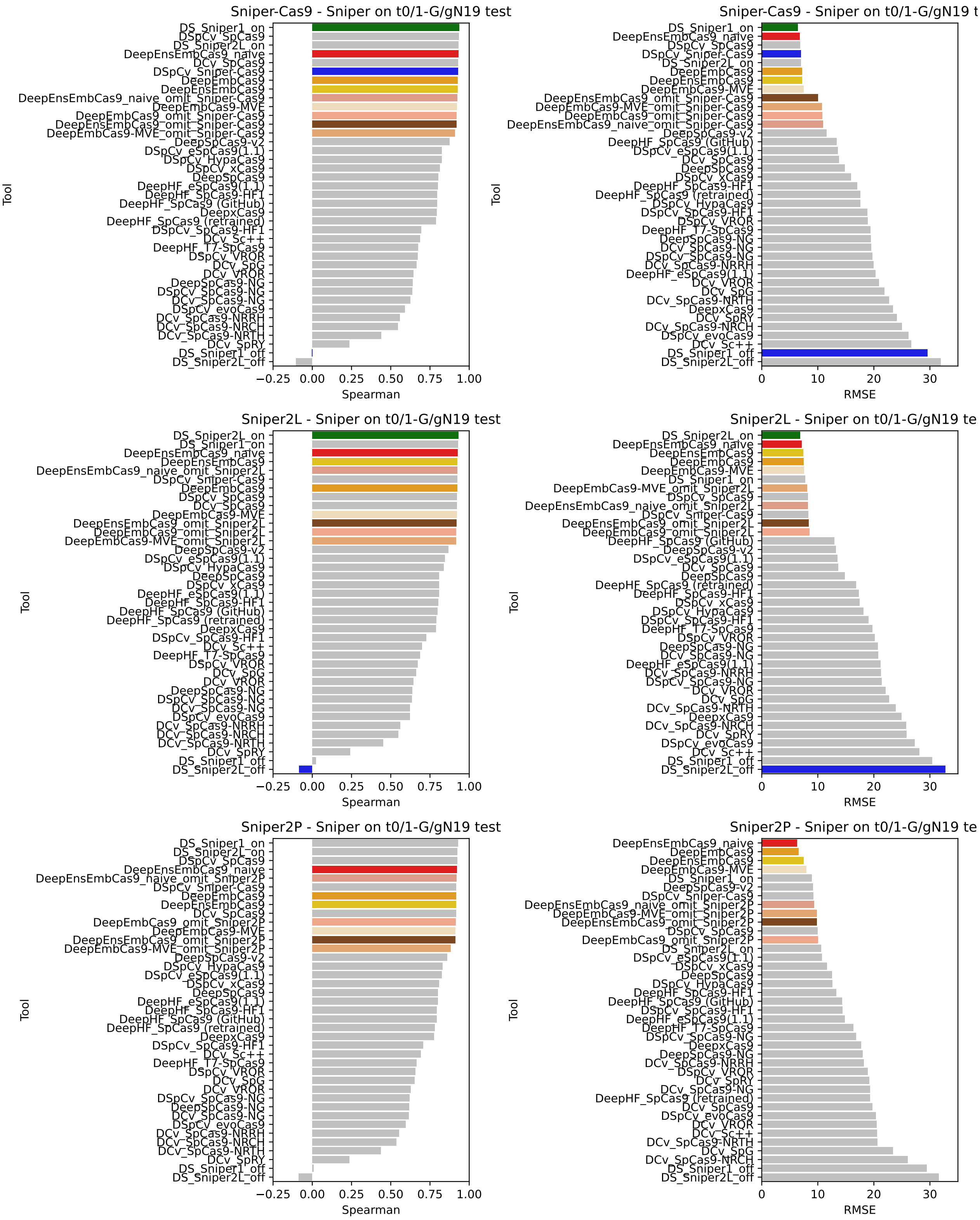


Figure S21: Benchmark test Spearman correlation (left) and RMSE (right) comparisons for DeepEmbCas9 (orange), DeepEnsEmbCas9_naive (red), DeepEmbCas9-MVE (wheat-colored), DeepEnsEmbCas9 (gold), DeepEmbCas9_omit (dark salmon) and DeepEnsEmbCas9_omit (brown) against relevant individual Cas9 cleavage activity tools for matched G/gN$_{19}$ and tRNA$^{\mathrm{Gln}}$-N$_{20}$ Sniper-Cas9 (top), Sniper2L (middle) and Sniper2P (bottom) interfaces from Kim, Kim and Okafor et al. [6], where green bars denote DeepSniper's Sniper1_on (top) and Sniper2L_on (middle), and blue bars denote other individual Cas9 cleavage activity tools trained on interfaces of the test nuclease.

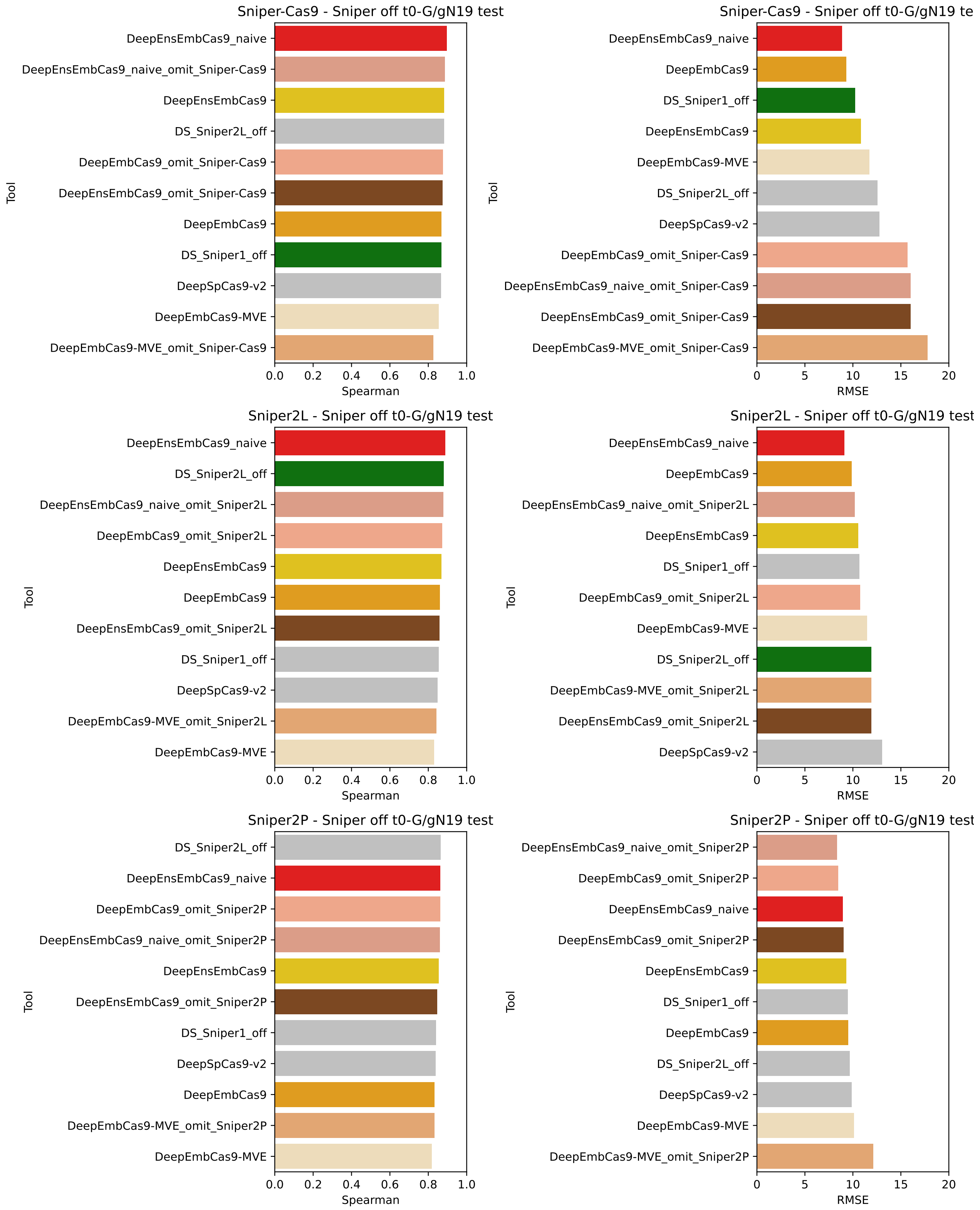


Figure S22: Benchmark test Spearman correlation (left) and RMSE (right) comparisons for DeepEmbCas9 (orange), DeepEnsEmbCas9_naive (red), DeepEmbCas9-MVE (wheat-colored), DeepEnsEmbCas9 (gold), DeepEmbCas9_omit (dark salmon) and DeepEnsEmbCas9_omit (brown) against relevant individual Cas9 cleavage activity tools for mismatched G/g$N_{19}$ Sniper-Cas9 (top), Sniper2L (middle) and Sniper2P (bottom) interfaces from Kim, Kim and Okafor et al. [6], where green bars denote DeepSniper's Sniper1_off (top) and Sniper2L_off (middle), and blue bars denote other individual Cas9 cleavage activity tools trained on interfaces of the test nuclease.

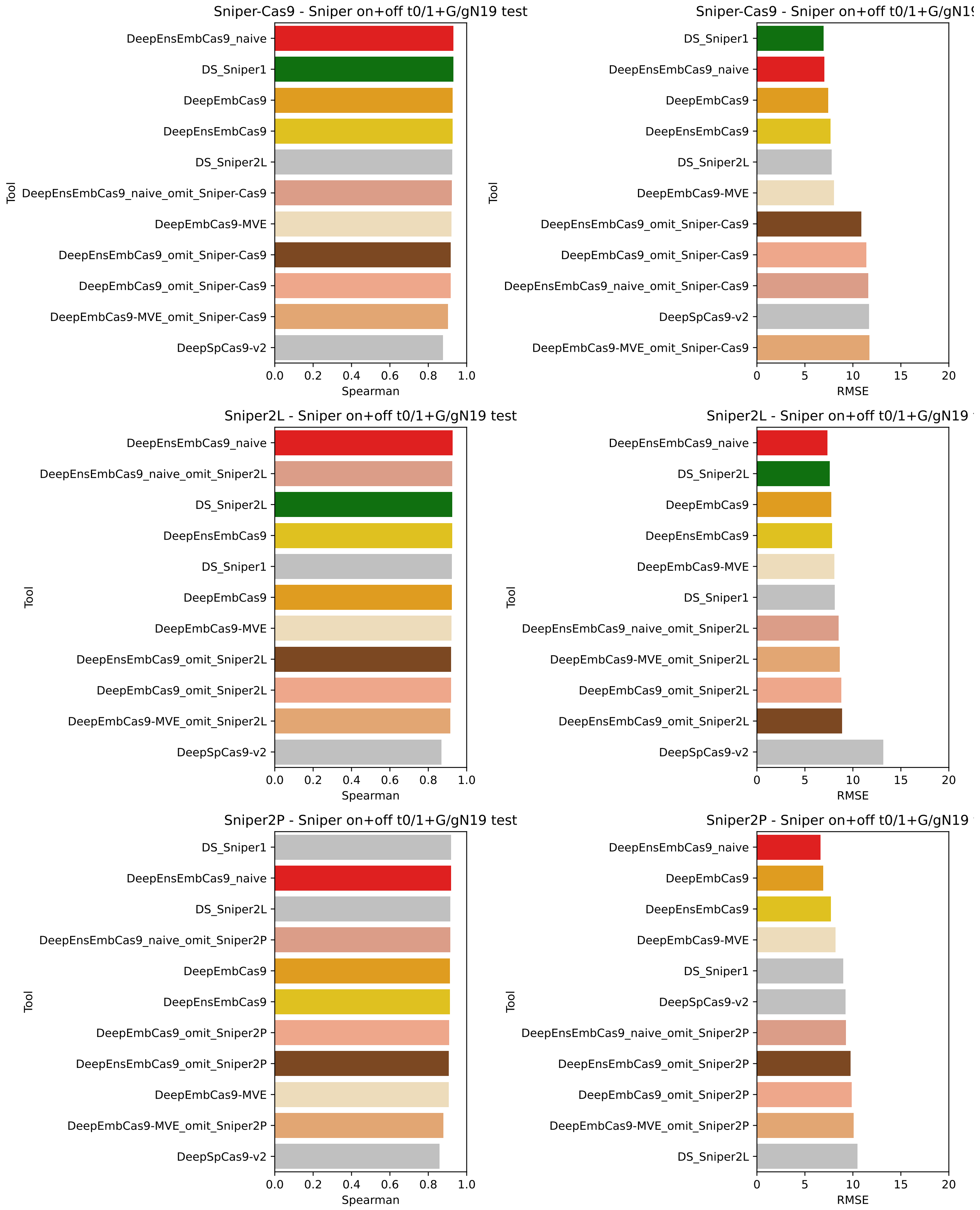


Figure S23: Benchmark test Spearman correlation (left) and RMSE (right) comparisons for DeepEmbCas9 (orange), DeepEnsEmbCas9_naive (red), DeepEmbCas9-MVE (wheat-colored), DeepEnsEmbCas9 (gold), DeepEmbCas9_omit (dark salmon) and DeepEnsEmbCas9_omit (brown) against relevant individual Cas9 cleavage activity tools for mismatched G/gN$_{19}$ Sniper-Cas9 (top), Sniper2L (middle) and Sniper2P (bottom) interfaces from Kim, Kim and Okafor et al. [6], where green bars denote DeepSniper, and blue bars denote other individual Cas9 cleavage activity tools trained on interfaces of the test nuclease.

##### 2.3.2 Small Cas9 variants

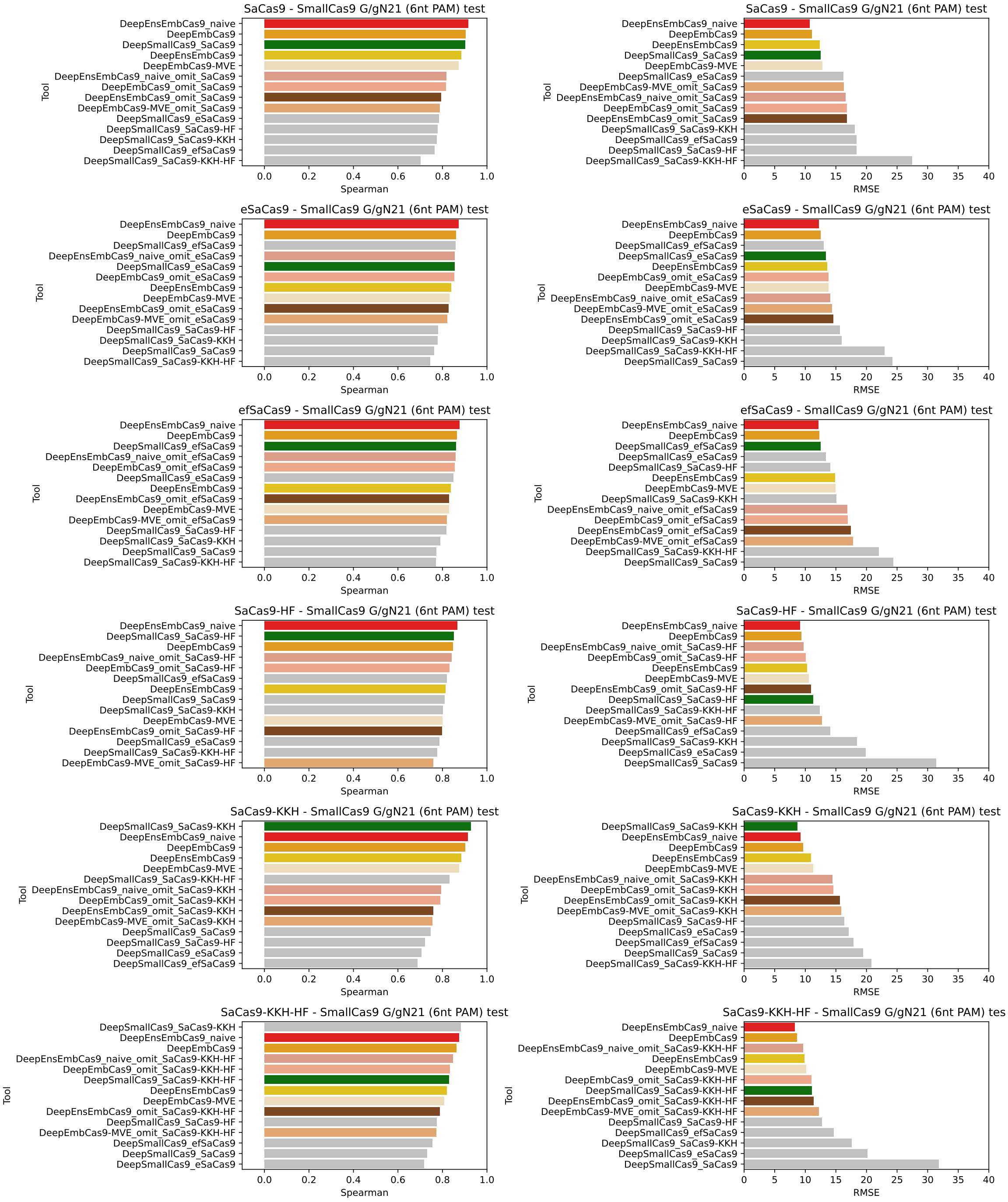


Figure S24: Benchmark test Spearman correlation (left) and RMSE (right) comparisons for DeepEmbCas9 (orange), DeepEnsEmbCas9 (red), DeepEmbCas9_omit (dark salmon) and DeepEnsEmbCas9_omit (brown) against relevant individual Cas9 cleavage activity tools for (mis)matched G/gN$_{21}$ wild type and engineered SaCas9 interfaces from Seo et al. [5], where green bars denote DeepSmallCas9.

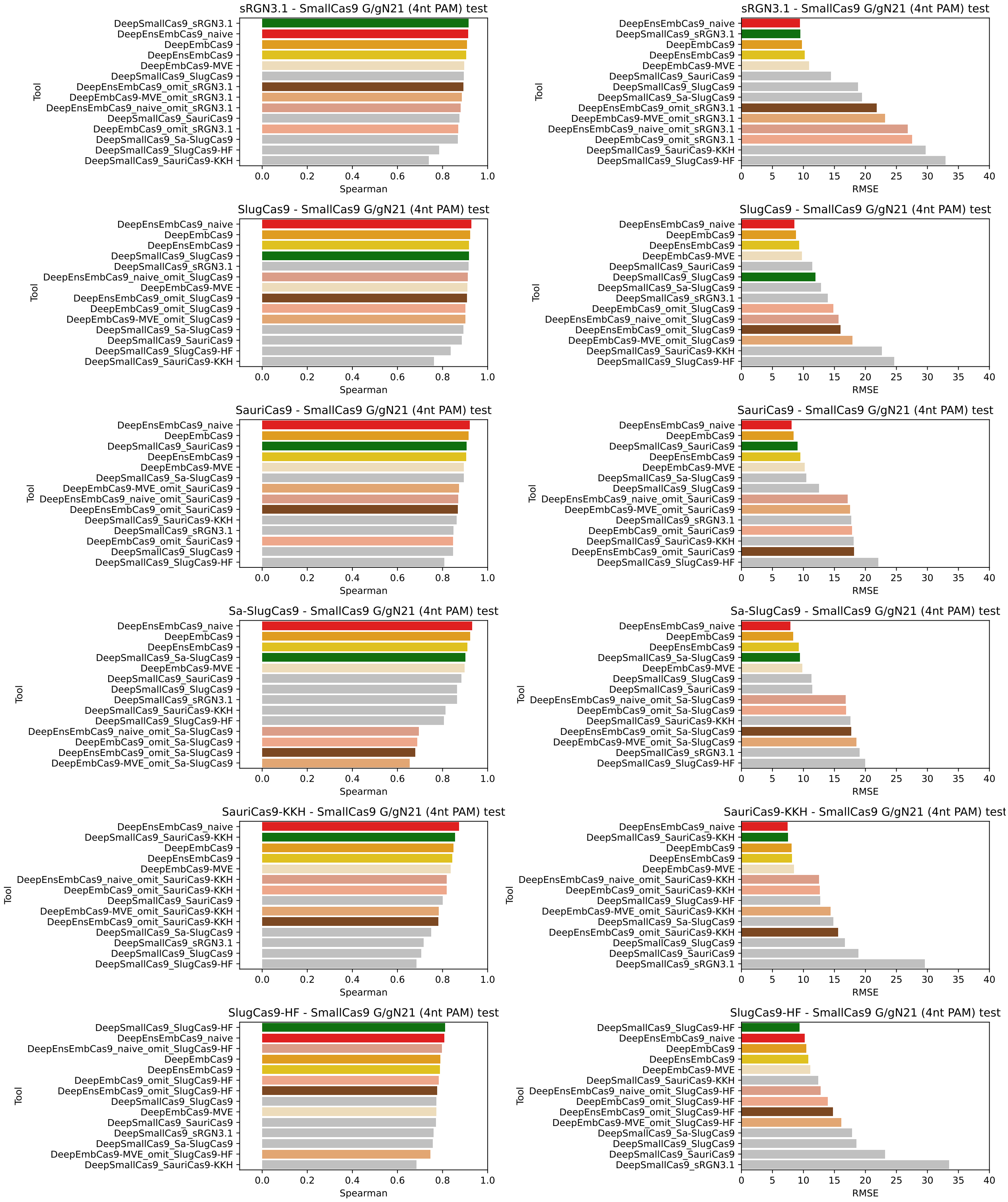


Figure S25: Benchmark test Spearman correlation (left) and RMSE (right) comparisons for DeepEmbCas9 (orange), DeepEnsEmbCas9 (red), DeepEmbCas9_omit (dark salmon) and DeepEnsEmbCas9_omit (brown) against relevant individual Cas9 cleavage activity tools for (mis)matched G/g$N_{21}$ wild type and engineered SlugCas9/sRGN3.1/SauriCas9 interfaces from Seo et al. [5], where green bars denote DeepSmallCas9.

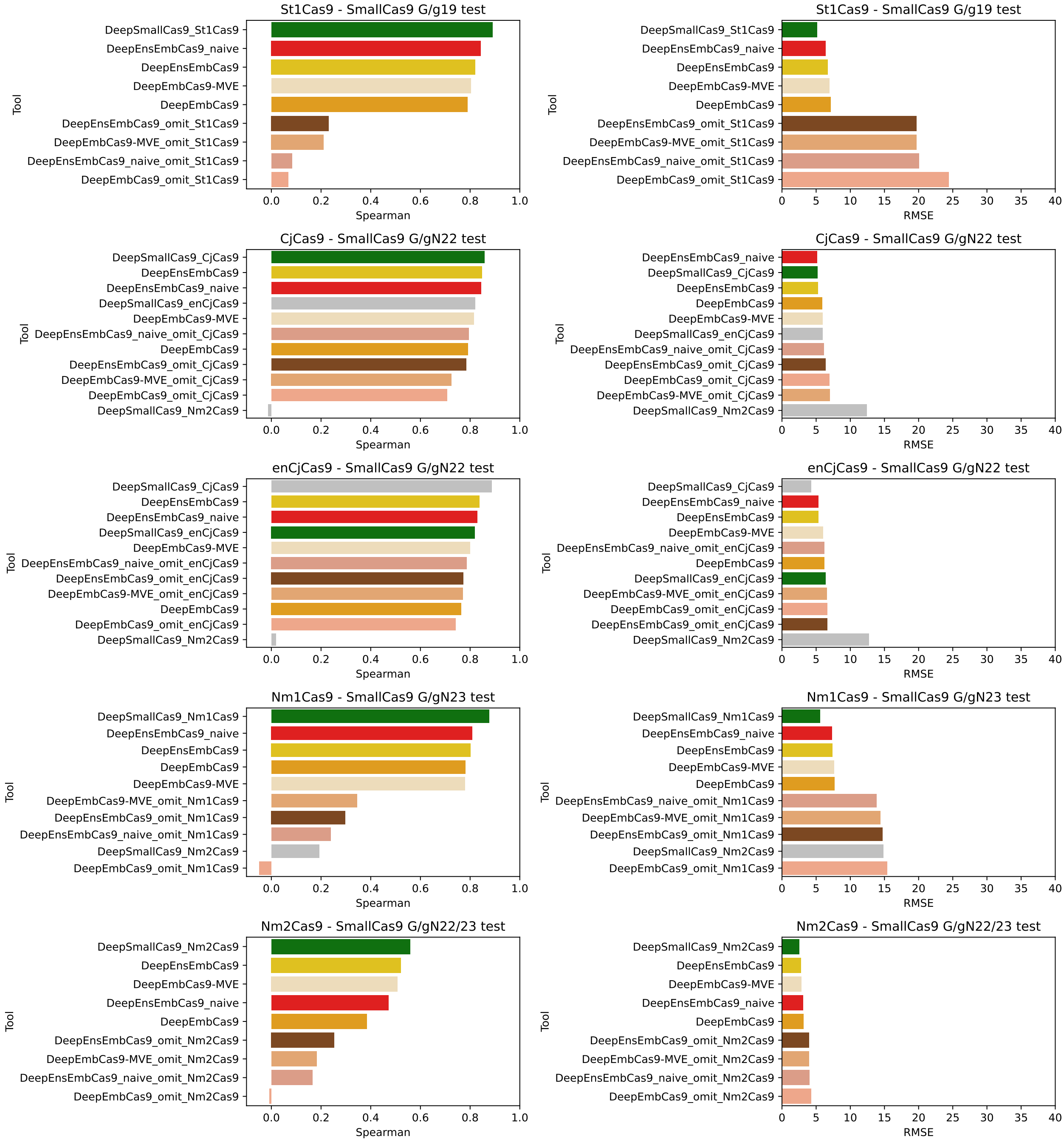


Figure S26: Benchmark test Spearman correlation (left) and RMSE (right) comparisons for DeepEmbCas9 (orange), DeepEnsEmbCas9 (red), DeepEmbCas9_omit (dark salmon) and DeepEnsEmbCas9_omit (brown) against relevant individual Cas9 cleavage activity tools for (mis)matched G/gN$_{21}$ G/gN$_{19}$ St1Cas9 (row 1), G/gN$_{22}$ CjCas9 (row 2), G/gN$_{22}$ enCjCas9 (row 3), G/gN$_{23}$ Nm1Cas9 (row 4) and G/gN$_{22-23}$ Nm2Cas9 (row 5) interfaces from Seo et al. [5], where green bars denote DeepSmallCas9.

#### 2.4 Extrapolation performance on whole dataset

| pLM | rLM | Guide-target 5-fold CV | Cas9 variants 5-fold CV | gRNA scaffold LOOCV | Overall |
|---|---|---|---|---|---|
| ESM-C-300M | BEACON-B | 0.8989 ± 0.0026 | 0.6993 ± 0.0944 | **0.5679 ± 0.1396** | **0.7220 ± 0.1666** |
| ESM-C-600M | RiNALMo | 0.8988 ± 0.0021 | **0.7200 ± 0.0767** | 0.5470 ± 0.1624 | 0.7219 ± 0.1759 |
| ESM-C-600M | RNA-FM | **0.9017 ± 0.0011** | 0.6865 ± 0.0842 | 0.5670 ± 0.1411 | 0.7184 ± 0.1696 |
| ESM-C-300M | RNA-FM | 0.8986 ± 0.0016 | 0.6996 ± 0.1040 | 0.5548 ± 0.1325 | 0.7177 ± 0.1726 |
| ESM-C-600M | BEACON-B512 | 0.8991 ± 0.0012 | 0.7016 ± 0.0674 | 0.5474 ± 0.1393 | 0.7160 ± 0.1763 |
| ESM-C-300M | RiNALMo | 0.8988 ± 0.0010 | 0.6991 ± 0.0811 | 0.5443 ± 0.1755 | 0.7141 ± 0.1777 |
| ESM-C-300M | BEACON-B512 | 0.8963 ± 0.0009 | 0.6899 ± 0.0976 | 0.5524 ± 0.1493 | 0.7129 ± 0.1731 |
| ESM-C-600M | evo-1-8k | 0.8980 ± 0.0007 | 0.7085 ± 0.0698 | 0.5247 ± 0.1636 | 0.7104 ± 0.1866 |
| ESM-C-6B | BEACON-B | 0.8951 ± 0.0016 | 0.7047 ± 0.0957 | 0.5302 ± 0.1634 | 0.7100 ± 0.1825 |
| ESM-C-600M | BEACON-B | 0.9013 ± 0.0016 | 0.6973 ± 0.0607 | 0.5289 ± 0.1650 | 0.7092 ± 0.1865 |
| ESM-C-6B | RNA-FM | 0.8977 ± 0.0031 | 0.6965 ± 0.0775 | 0.5306 ± 0.1972 | 0.7083 ± 0.1839 |
| ESM-C-6B | BEACON-B512 | 0.8919 ± 0.0018 | 0.6937 ± 0.0892 | 0.5334 ± 0.1674 | 0.7064 ± 0.1796 |
| ESM-C-300M | evo-1-8k | 0.8947 ± 0.0009 | 0.6985 ± 0.0876 | 0.5228 ± 0.1294 | 0.7053 ± 0.1860 |
| ProtT5 | evo-1-8k | 0.8898 ± 0.0033 | 0.6725 ± 0.1078 | 0.5456 ± 0.1433 | 0.7026 ± 0.1741 |
| ESM-C-6B | RiNALMo | 0.8947 ± 0.0020 | 0.6843 ± 0.0856 | 0.5242 ± 0.1808 | 0.7011 ± 0.1858 |
| Ankh-large | RNA-FM | 0.8914 ± 0.0026 | 0.6593 ± 0.1050 | 0.5438 ± 0.1577 | 0.6982 ± 0.1770 |
| Ankh-large | BEACON-B | 0.8919 ± 0.0016 | 0.6600 ± 0.1020 | 0.5341 ± 0.1781 | 0.6953 ± 0.1815 |
| ProtT5 | RNA-FM | 0.8950 ± 0.0016 | 0.6418 ± 0.1175 | 0.5470 ± 0.1636 | 0.6946 ± 0.1799 |
| Ankh-large | BEACON-B512 | 0.8873 ± 0.0015 | 0.6608 ± 0.0943 | 0.5323 ± 0.1484 | 0.6935 ± 0.1797 |
| ProtT5 | RiNALMo | 0.8934 ± 0.0033 | 0.6687 ± 0.1049 | 0.5168 ± 0.2108 | 0.6930 ± 0.1895 |
| ProtT5 | BEACON-B512 | 0.8890 ± 0.0022 | 0.6504 ± 0.0957 | 0.5379 ± 0.1812 | 0.6924 ± 0.1793 |
| ProtT5 | BEACON-B | 0.8927 ± 0.0026 | 0.6496 ± 0.1228 | 0.5223 ± 0.2016 | 0.6882 ± 0.1882 |
| Ankh-large | evo-1-8k | 0.8903 ± 0.0013 | 0.6552 ± 0.1163 | 0.5178 ± 0.1452 | 0.6878 ± 0.1884 |
| Ankh-large | RiNALMo | 0.8913 ± 0.0024 | 0.6503 ± 0.1159 | 0.5182 ± 0.2031 | 0.6866 ± 0.1892 |
| ESM-C-6B | evo-1-8k | 0.8898 ± 0.0013 | 0.6747 ± 0.0997 | 0.4879 ± 0.1958 | 0.6841 ± 0.2011 |
| gLM2-650M | BEACON-B512 | 0.8193 ± 0.0024 | 0.6447 ± 0.0730 | 0.4594 ± 0.1553 | 0.6411 ± 0.1800 |
| gLM2-650M | RNA-FM | 0.8251 ± 0.0035 | 0.6386 ± 0.0512 | 0.4514 ± 0.2073 | 0.6384 ± 0.1869 |
| ESM3 | RNA-FM | 0.8239 ± 0.0042 | 0.6325 ± 0.1043 | 0.4531 ± 0.1879 | 0.6365 ± 0.1854 |
| gLM2-650M | evo-1-8k | 0.8193 ± 0.0031 | 0.6614 ± 0.0645 | 0.4251 ± 0.1809 | 0.6353 ± 0.1984 |
| gLM2-650M | RiNALMo | 0.7964 ± 0.0725 | 0.6252 ± 0.0477 | 0.4705 ± 0.1621 | 0.6307 ± 0.1630 |
| gLM2-650M | BEACON-B | 0.8239 ± 0.0043 | 0.5894 ± 0.0987 | 0.4675 ± 0.1719 | 0.6269 ± 0.1811 |
| ESM3 | BEACON-B512 | 0.8007 ± 0.0415 | 0.6301 ± 0.1038 | 0.4127 ± 0.2113 | 0.6145 ± 0.1944 |
| ESM3 | RiNALMo | 0.7954 ± 0.0671 | 0.6136 ± 0.1152 | 0.4262 ± 0.1379 | 0.6117 ± 0.1846 |
| ESM3 | evo-1-8k | 0.8225 ± 0.0021 | 0.5896 ± 0.1003 | 0.4191 ± 0.1518 | 0.6104 ± 0.2025 |
| ESM3 | BEACON-B | 0.7397 ± 0.0826 | 0.5351 ± 0.0827 | 0.3915 ± 0.1875 | 0.5554 ± 0.1750 |

Table S6: Test Spearman correlation of the 30 pLM-rLM embedding combinations (arising from 6 pLM (Ankh-large, ESM3, ESM-C-300M, ESM-C-600M, ESM-C-6B, gLM2-650M, ProtT5) and 5 rLM (BEACON-B, BEACON-B512, RNA-FM, RiNALMo, evo-1-8k) embeddings) considered for Deep-EmbCas9 across three tasks — guide-target 5-fold cross validation (CV), Cas9 variants 5-fold cross validation and gRNA scaffold leave-one-out cross validation (LOOCV). The pLM-rLM combinations are ranked by decreasing "Overall" score, which denotes the average between the mean performances in the three tasks.

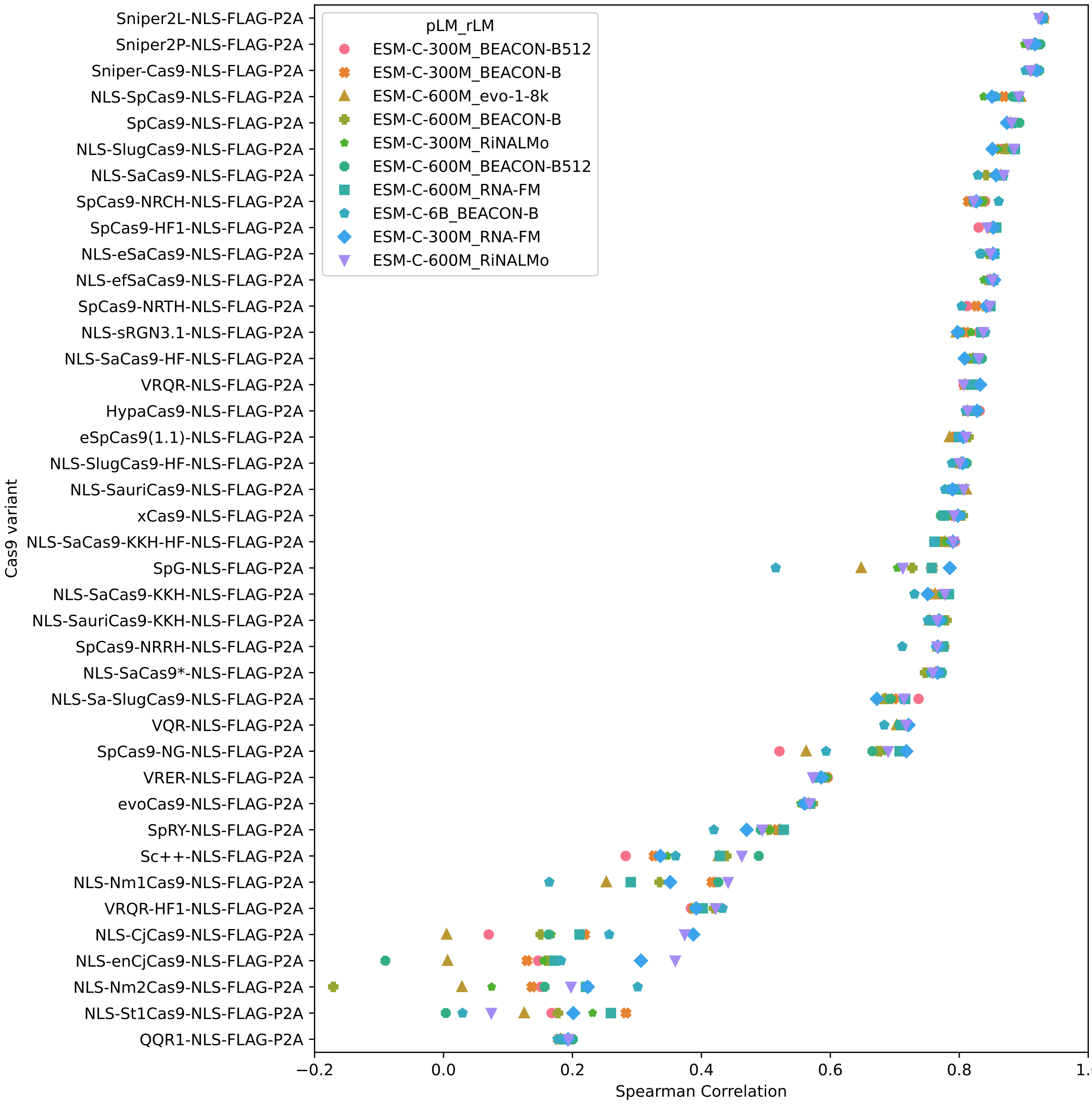


Figure S27: Test Spearman correlations when holding out data associated with one Cas9 variant for testing for the 10 pLM-rLM combinations with the highest "Overall" score in Table S6 (see list of Cas9 mutations in Table S4), with Cas9 variants roughly sorted in descending Spearman correlation.

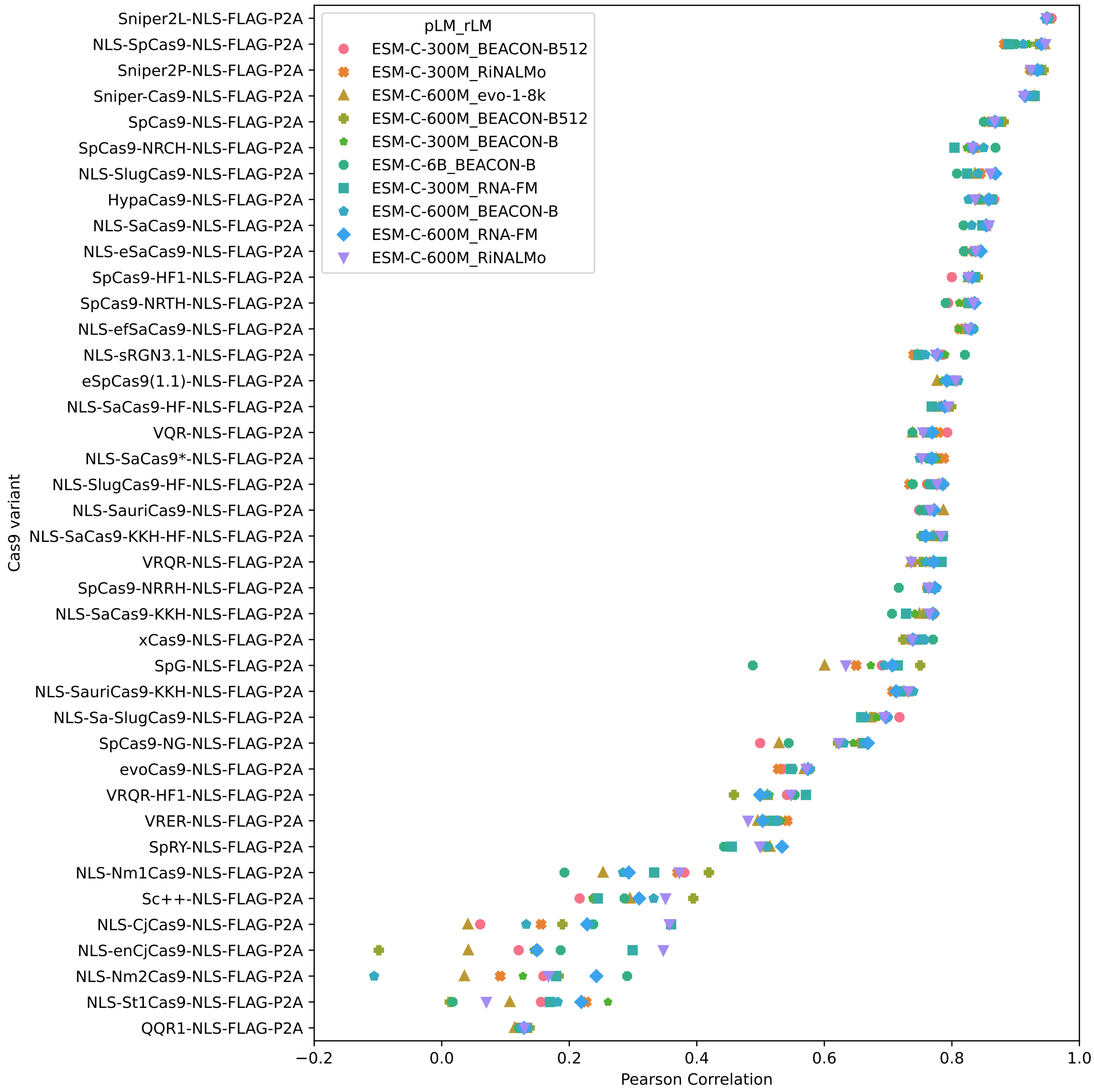


Figure S28: Test Pearson correlation of the 18 pLM-rLM embedding combinations (arising from 6 pLM (Ankh-large, ESM3, ESM-C-300M, ESM-C-600M, ESM-C-6B, gLM2-650M, ProtT5) and 3 rLM (BEACON-B, BEACON-B512, RNA-FM) embeddings) considered for DeepEmbCas9 across three tasks — guide-target 5-fold cross validation (CV), Cas9 variants 5-fold cross validation and gRNA scaffold leave-one-out cross validation (LOOCV). The pLM-rLM combinations are ranked by decreasing "Overall" score, which denotes the average between the mean performances in the three tasks.

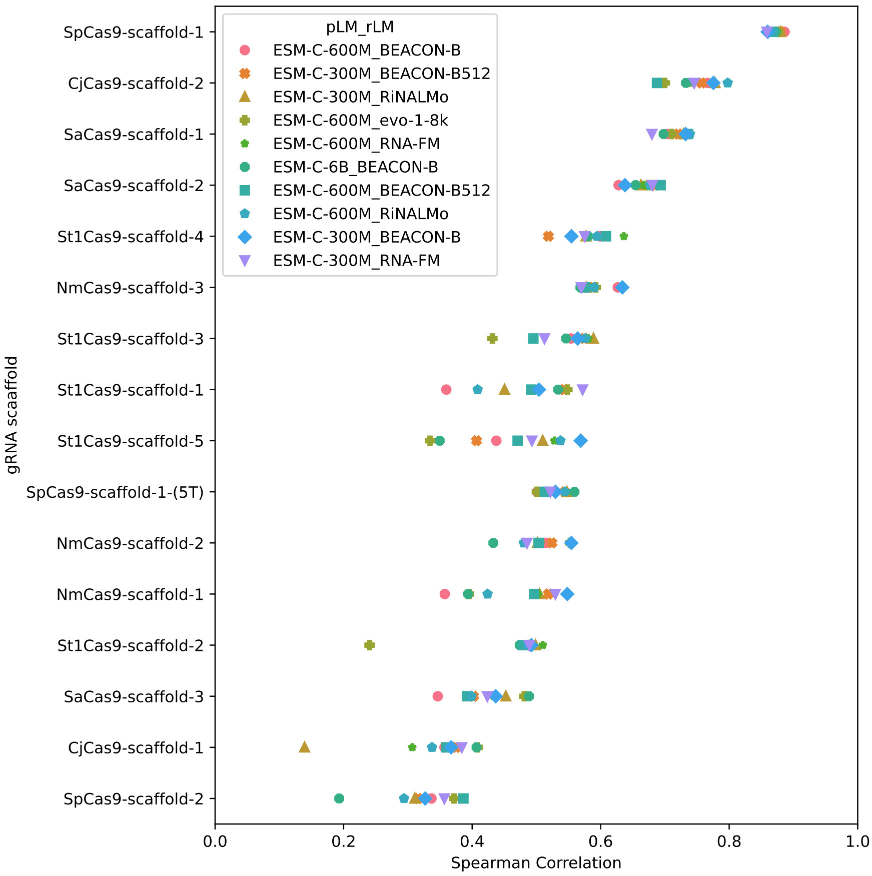


Figure S29: Test Spearman correlations when holding out data associated with one gRNA scaffold for testing for the 10 pLM-rLM combinations with the highest "Overall" score in Table S6, with Cas9 variants roughly sorted in descending Spearman correlation.

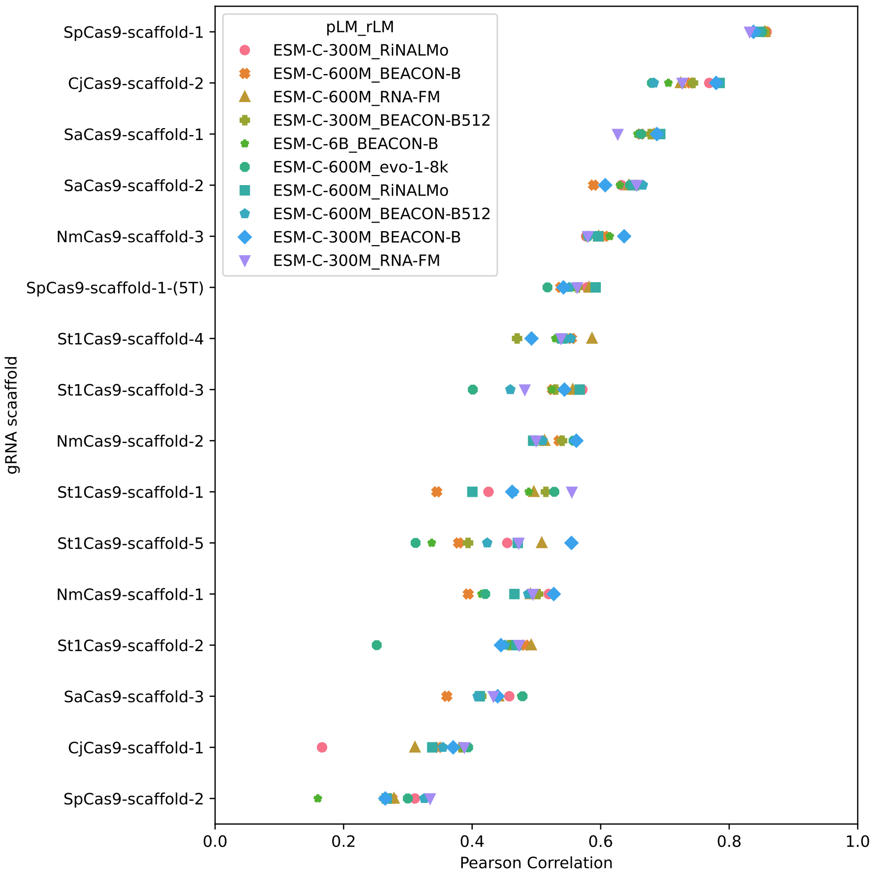


Figure S30: Test Pearson correlations when holding out data associated with one Cas9 variant for testing for 10 pLM-rLM combinations with the highest "Overall" score in Table S6 (see list of Cas9 mutations in Table S4).

#### 2.5 In-distribution calibration

##### 2.5.1 Quantile calibration plots - DeepEnsEmbCas9

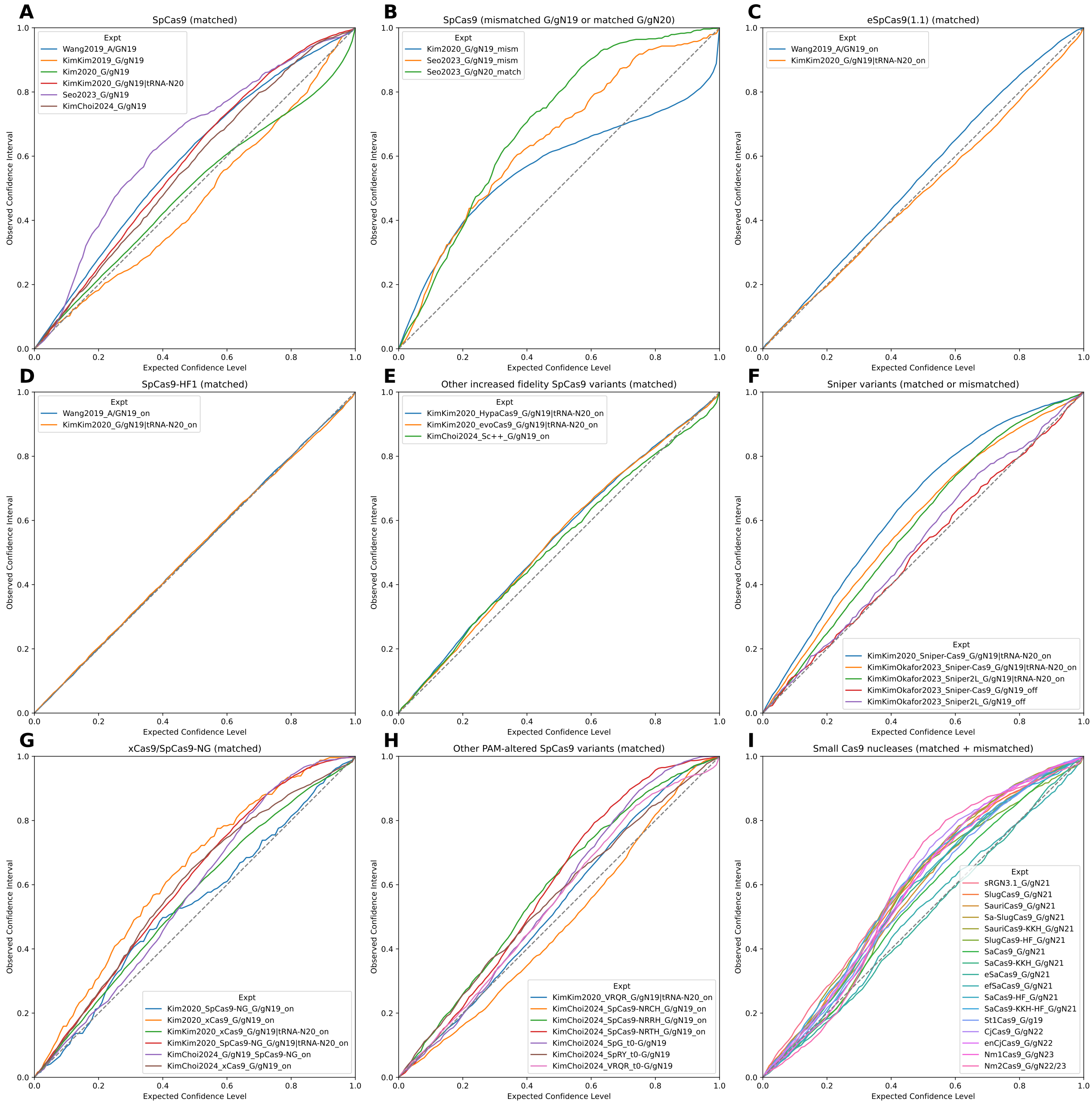


Figure S31: Confidence interval-based calibration curves for DeepEnsEmbCas9, conditioned on (A) matched $AN_{19}$, $G/gN_{19}$ and $tRNA^{Gln}$-$N_{20}$ wild type SpCas9 interfaces; (B) mismatched $G/gN_{19}$ and matched $G/gN_{20}$ wild type SpCas9 interfaces; (C) matched $AN_{19}$, $G/gN_{19}$ and $tRNA^{Gln}$-$N_{20}$ eSpCas9(1.1) interfaces; (D) matched $AN_{19}$, $G/gN_{19}$ and $tRNA^{Gln}$-$N_{20}$ SpCas9-HF1 interfaces; for matched $G/gN_{19}$ and $tRNA^{Gln}$-$N_{20}$ HypaCas9/evoCas9 and $G/gN_{19}$ Sc++ interfaces; (F) matched $G/gN_{19}$ and $tRNA^{Gln}$-$N_{20}$ and mismatched $G/gN_{19}$ interfaces for 2 Sniper variants; (G,H) matched $G/gN_{19}$ and $tRNA^{Gln}$-$N_{20}$ interfaces for xCas9/SpCas9-NG (G) and 6 other PAM-altered SpCas9 variants (H); and (I) matched and mismatched interfaces for 17 wild type or engineered small Cas9 nucleases.

##### 2.5.2 Quantile calibration plots - DeepEmbCas9-MVE

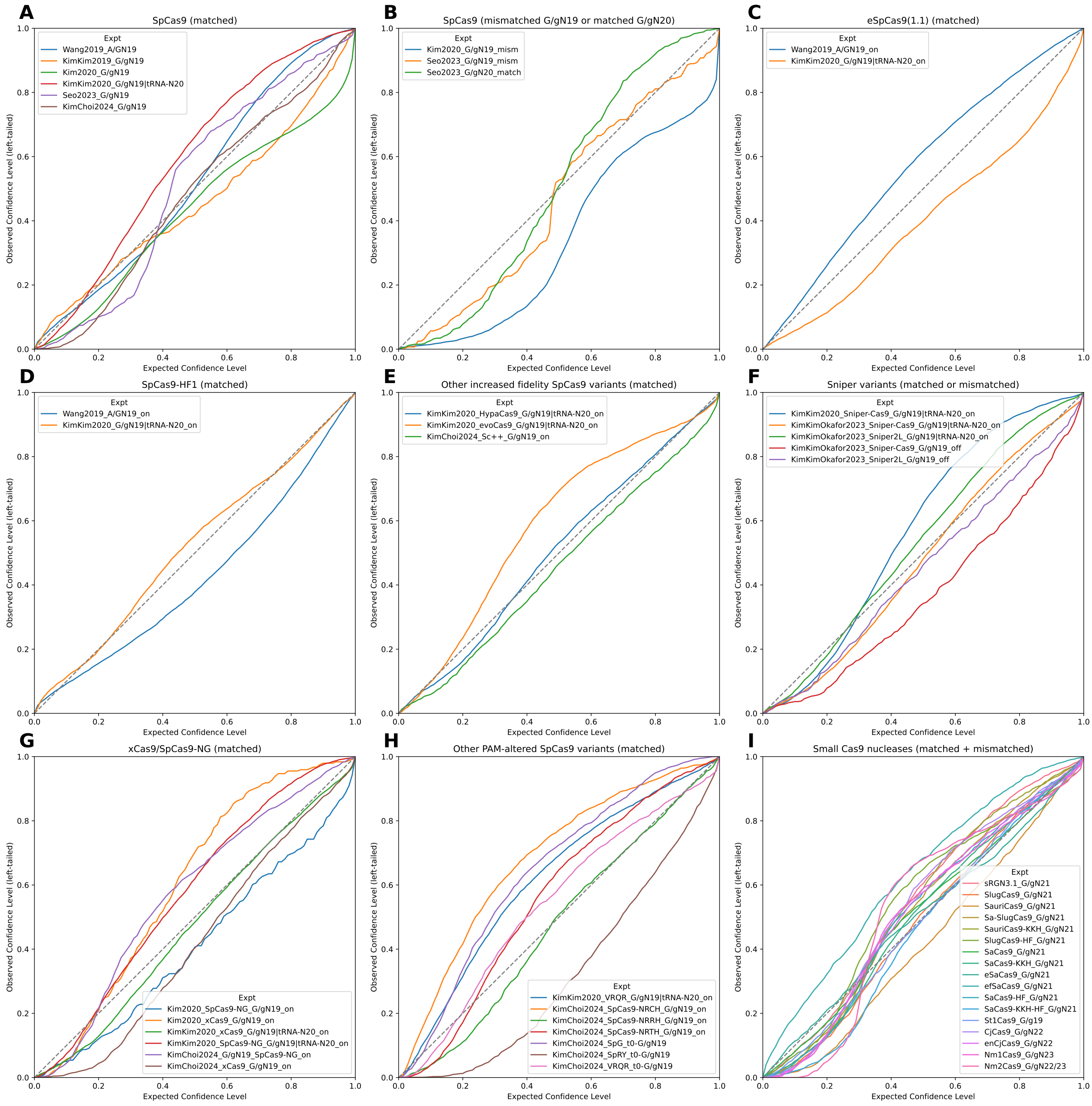


Figure S32: Quantile calibration plots for DeepEmbCas9-MVE, conditioned on (A) matched $AN_{19}$, $G/gN_{19}$ and $tRNA^{Gln}$-$N_{20}$ wild type SpCas9 interfaces; (B) mismatched $G/gN_{19}$ and matched $G/gN_{20}$ wild type SpCas9 interfaces; (C) matched $AN_{19}$, $G/gN_{19}$ and $tRNA^{Gln}$-$N_{20}$ eSpCas9(1.1) interfaces; (D) matched $AN_{19}$, $G/gN_{19}$ and $tRNA^{Gln}$-$N_{20}$ SpCas9-HF1 interfaces; for matched $G/gN_{19}$ and $tRNA^{Gln}$-$N_{20}$ HypaCas9/evoCas9 and $G/gN_{19}$ Sc++ interfaces; (F) matched $G/gN_{19}$ and $tRNA^{Gln}$-$N_{20}$ and mismatched $G/gN_{19}$ interfaces for 2 Sniper variants; (G,H) matched $G/gN_{19}$ and $tRNA^{Gln}$-$N_{20}$ interfaces for xCas9/SpCas9-NG (G) and 6 other PAM-altered SpCas9 variants (H); and (I) matched and mismatched interfaces for 17 wild type or engineered small Cas9 nucleases.

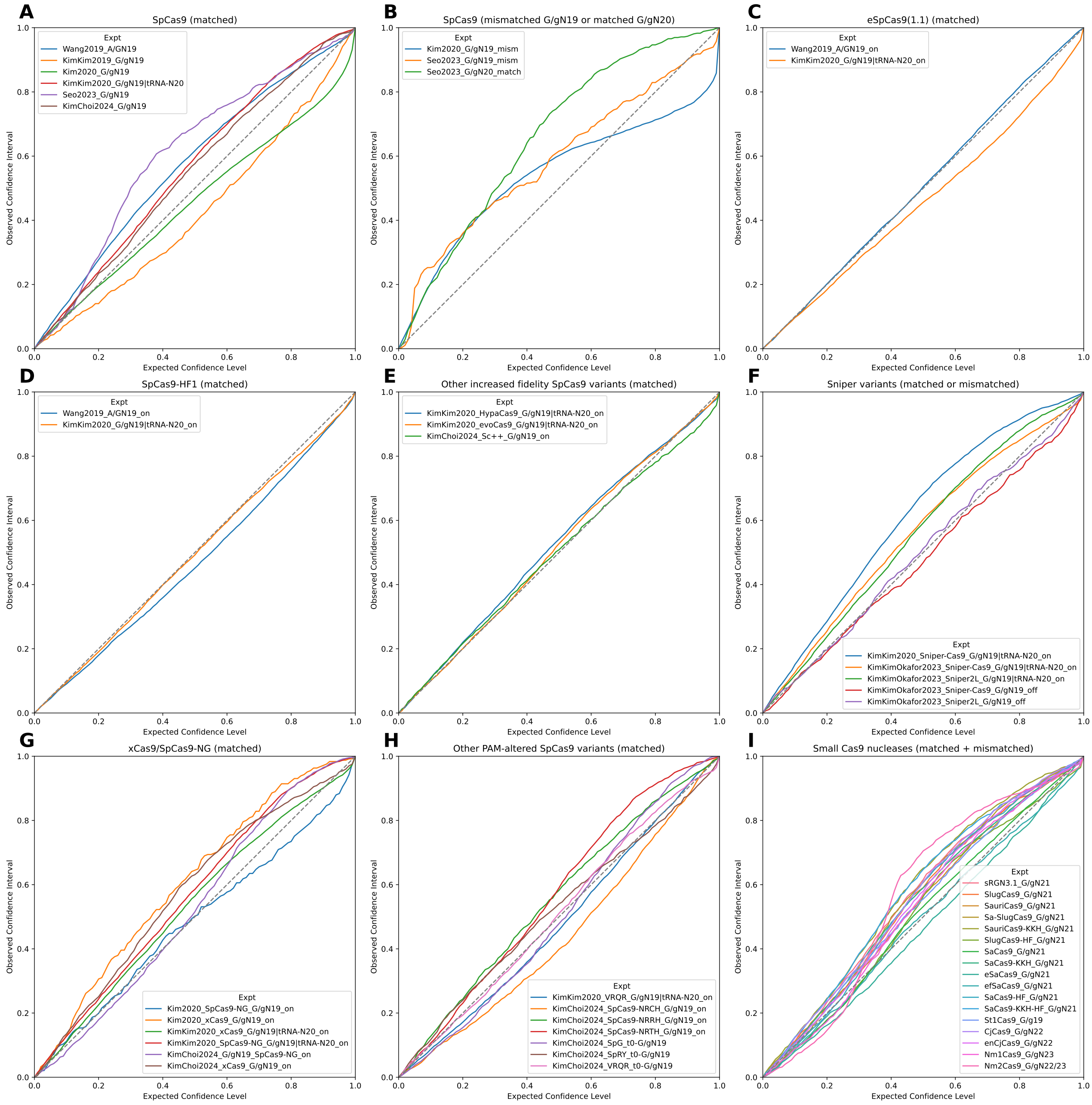


Figure S33: Confidence interval-based calibration curves for DeepEmbCas9-MVE, conditioned on (A) matched $AN_{19}$, $G/gN_{19}$ and $tRNA^{Gln}$-$N_{20}$ wild type SpCas9 interfaces; (B) mismatched $G/gN_{19}$ and matched $G/gN_{20}$ wild type SpCas9 interfaces; (C) matched $AN_{19}$, $G/gN_{19}$ and $tRNA^{Gln}$-$N_{20}$ eSpCas9(1.1) interfaces; (D) matched $AN_{19}$, $G/gN_{19}$ and $tRNA^{Gln}$-$N_{20}$ SpCas9-HF1 interfaces; for matched $G/gN_{19}$ and $tRNA^{Gln}$-$N_{20}$ HypaCas9/evoCas9 and $G/gN_{19}$ Sc++ interfaces; (F) matched $G/gN_{19}$ and $tRNA^{Gln}$-$N_{20}$ and mismatched $G/gN_{19}$ interfaces for 2 Sniper variants; (G,H) matched $G/gN_{19}$ and $tRNA^{Gln}$-$N_{20}$ interfaces for xCas9/SpCas9-NG (G) and 6 other PAM-altered SpCas9 variants (H); and (I) matched and mismatched interfaces for 17 wild type or engineered small Cas9 nucleases.

##### 2.5.3 Quantile calibration plots - DeepEnsEmbCas9_naive

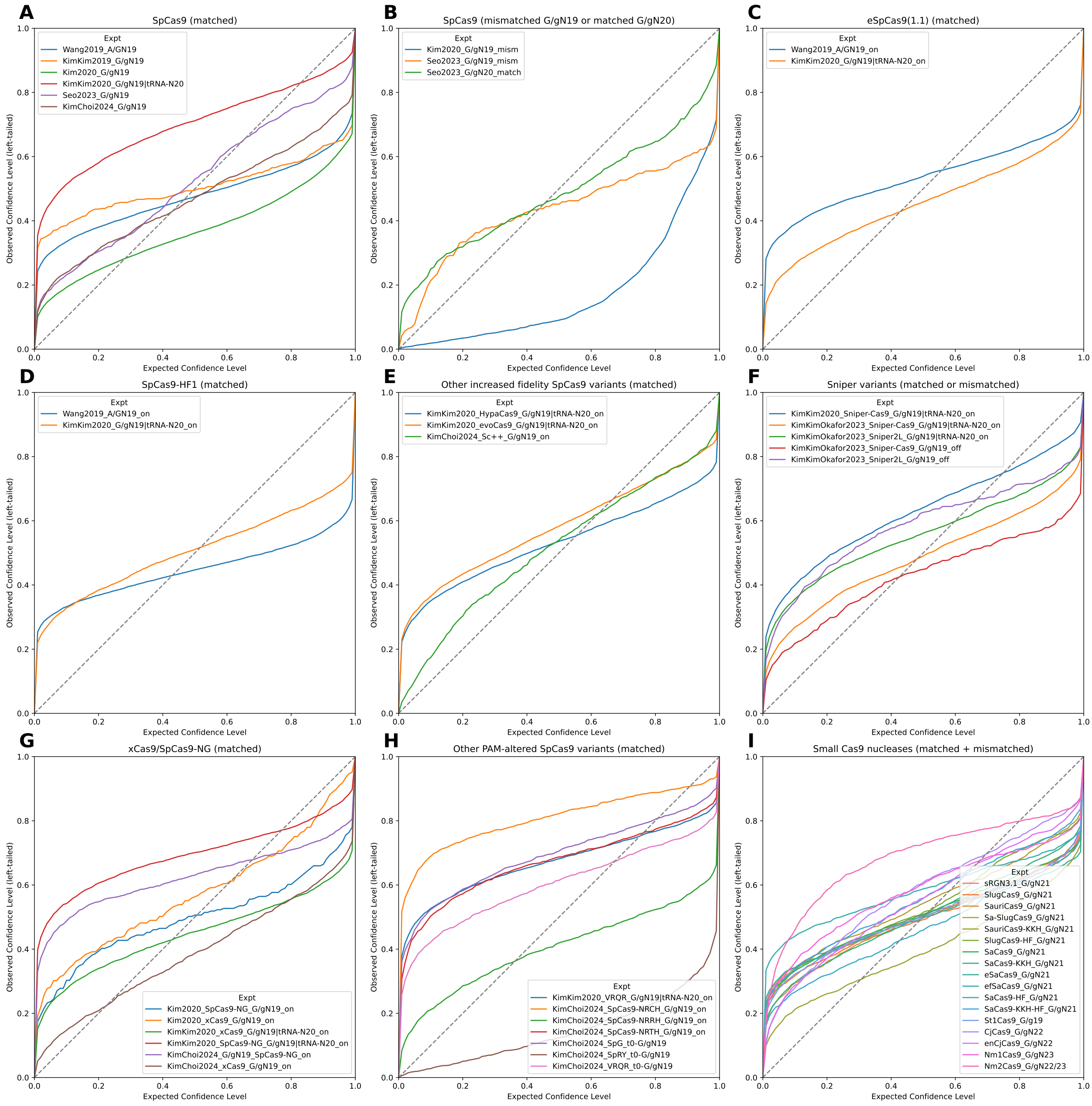


Figure S34: Quantile calibration plots for DeepEnsEmbCas9_naive, conditioned on (A) matched AN$_{19}$, G/gN$_{19}$ and tRNA$^{Gln}$-N$_{20}$ wild type SpCas9 interfaces; (B) mismatched G/gN$_{19}$ and matched G/gN$_{20}$ wild type SpCas9 interfaces; (C) matched AN$_{19}$, G/gN$_{19}$ and tRNA$^{Gln}$-N$_{20}$ eSpCas9(1.1) interfaces; (D) matched AN$_{19}$, G/gN$_{19}$ and tRNA$^{Gln}$-N$_{20}$ SpCas9-HF1 interfaces; for matched G/gN$_{19}$ and tRNA$^{Gln}$-N$_{20}$ HypaCas9/evoCas9 and G/gN$_{19}$ Sc++ interfaces; (F) matched G/gN$_{19}$ and tRNA$^{Gln}$-N$_{20}$ and mismatched G/gN$_{19}$ interfaces for 2 Sniper variants; (G,H) matched G/gN$_{19}$ and tRNA$^{Gln}$-N$_{20}$ interfaces for xCas9/SpCas9-NG (G) and 6 other PAM-altered SpCas9 variants (H); and (I) matched and mismatched interfaces for 17 wild type or engineered small Cas9 nucleases.

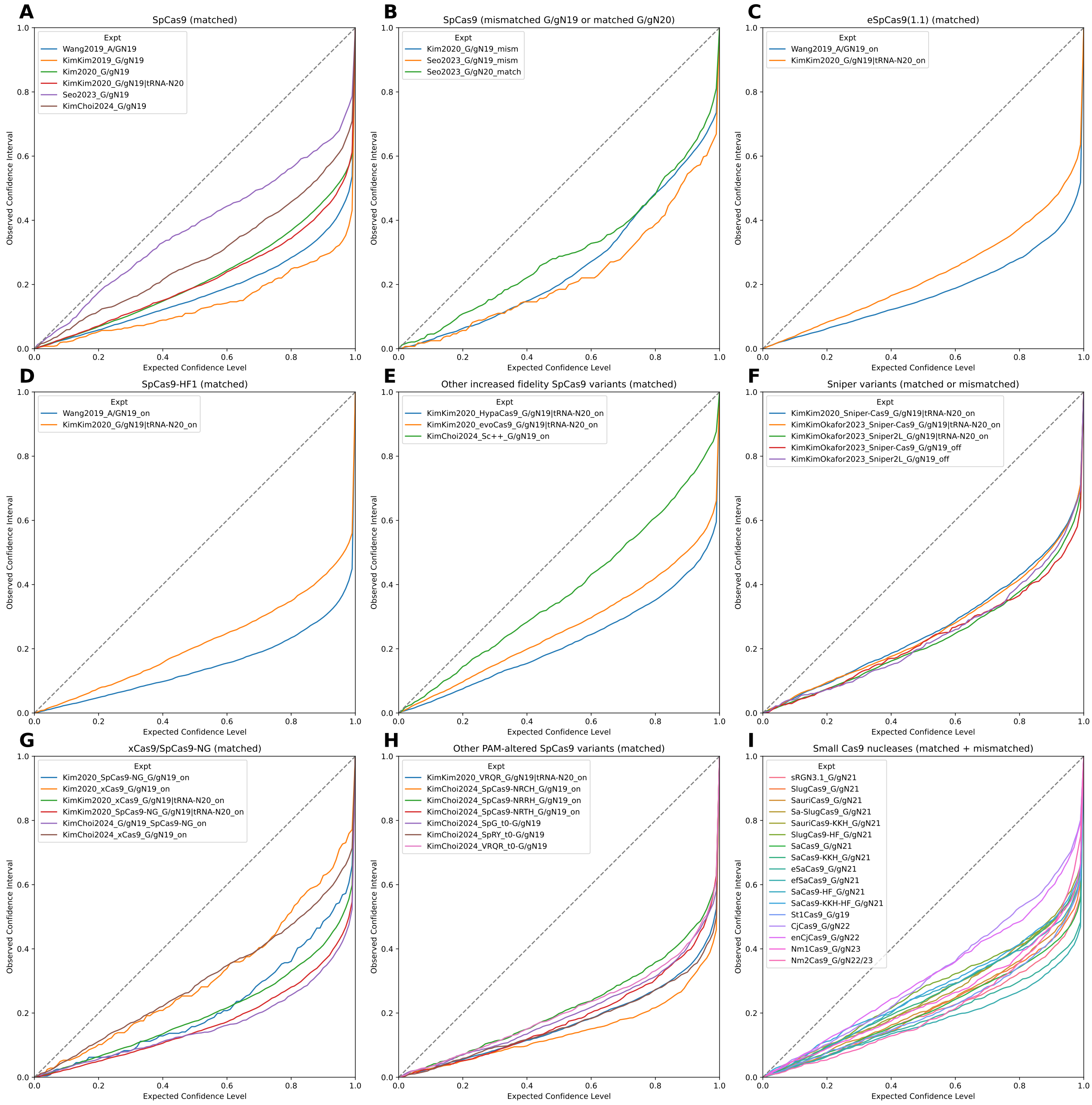


Figure S35: Confidence interval-based calibration curves for DeepEnsEmbCas9_naive, conditioned on (A) matched $AN_{19}$, $G/gN_{19}$ and $tRNA^{Gln}$-$N_{20}$ wild type SpCas9 interfaces; (B) mismatched $G/gN_{19}$ and matched $G/gN_{20}$ wild type SpCas9 interfaces; (C) matched $AN_{19}$, $G/gN_{19}$ and $tRNA^{Gln}$-$N_{20}$ eSpCas9(1.1) interfaces; (D) matched $AN_{19}$, $G/gN_{19}$ and $tRNA^{Gln}$-$N_{20}$ SpCas9-HF1 interfaces; for matched $G/gN_{19}$ and $tRNA^{Gln}$-$N_{20}$ HypaCas9/evoCas9 and $G/gN_{19}$ Sc++ interfaces; (F) matched $G/gN_{19}$ and $tRNA^{Gln}$-$N_{20}$ and mismatched $G/gN_{19}$ interfaces for 2 Sniper variants; (G,H) matched $G/gN_{19}$ and $tRNA^{Gln}$-$N_{20}$ interfaces for xCas9/SpCas9-NG (G) and 6 other PAM-altered SpCas9 variants (H); and (I) matched and mismatched interfaces for 17 wild type or engineered small Cas9 nucleases.

##### 2.5.4 Quantile calibration error

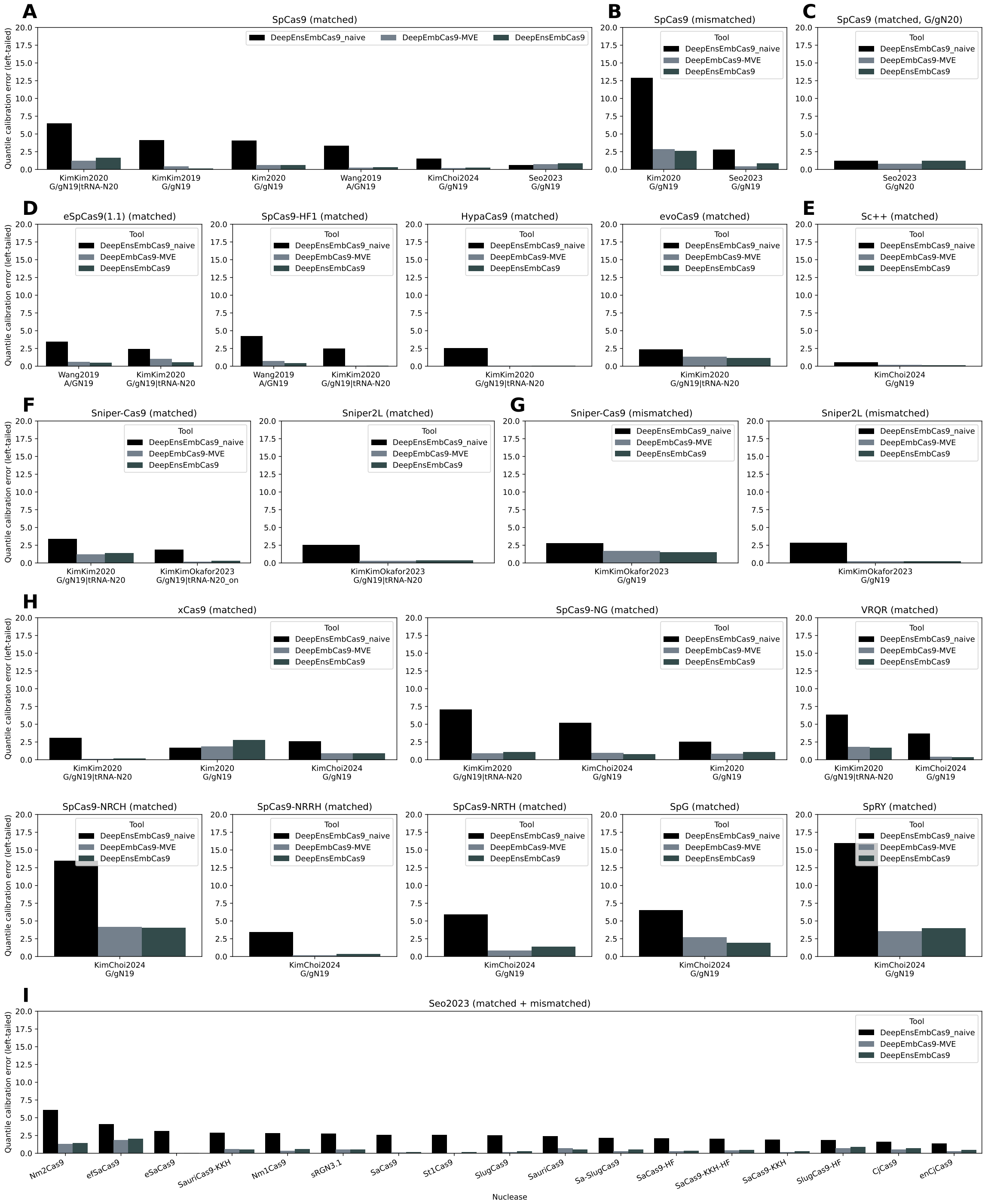


Figure S36: Quantile calibration errors for DeepEnsEmbCas9_naive, DeepEmbCas9-MVE and DeepEnsEmbCas9, conditioned on (A) matched $AN_{19}$, $G/gN_{19}$ and $tRNA^{Gln}$-$N_{20}$ wild type SpCas9 interfaces; (B) mismatched $G/gN_{19}$ and matched $G/gN_{20}$ wild type SpCas9 interfaces; (C) matched $AN_{19}$, $G/gN_{19}$ and $tRNA^{Gln}$-$N_{20}$ eSpCas9(1.1) interfaces; (D) matched $AN_{19}$, $G/gN_{19}$ and $tRNA^{Gln}$-$N_{20}$ SpCas9-HF1 interfaces; for matched $G/gN_{19}$ and $tRNA^{Gln}$-$N_{20}$ HypaCas9/evoCas9 and $G/gN_{19}$ Sc++ interfaces; (F) matched $G/gN_{19}$ and $tRNA^{Gln}$-$N_{20}$ and mismatched $G/gN_{19}$ interfaces for 2 Sniper variants; (G,H) matched $G/gN_{19}$ and $tRNA^{Gln}$-$N_{20}$ interfaces for xCas9/SpCas9-NG (G) and 6 other PAM-altered SpCas9 variants (H); and (I) matched and mismatched interfaces for 17 wild type or engineered small Cas9 nucleases.

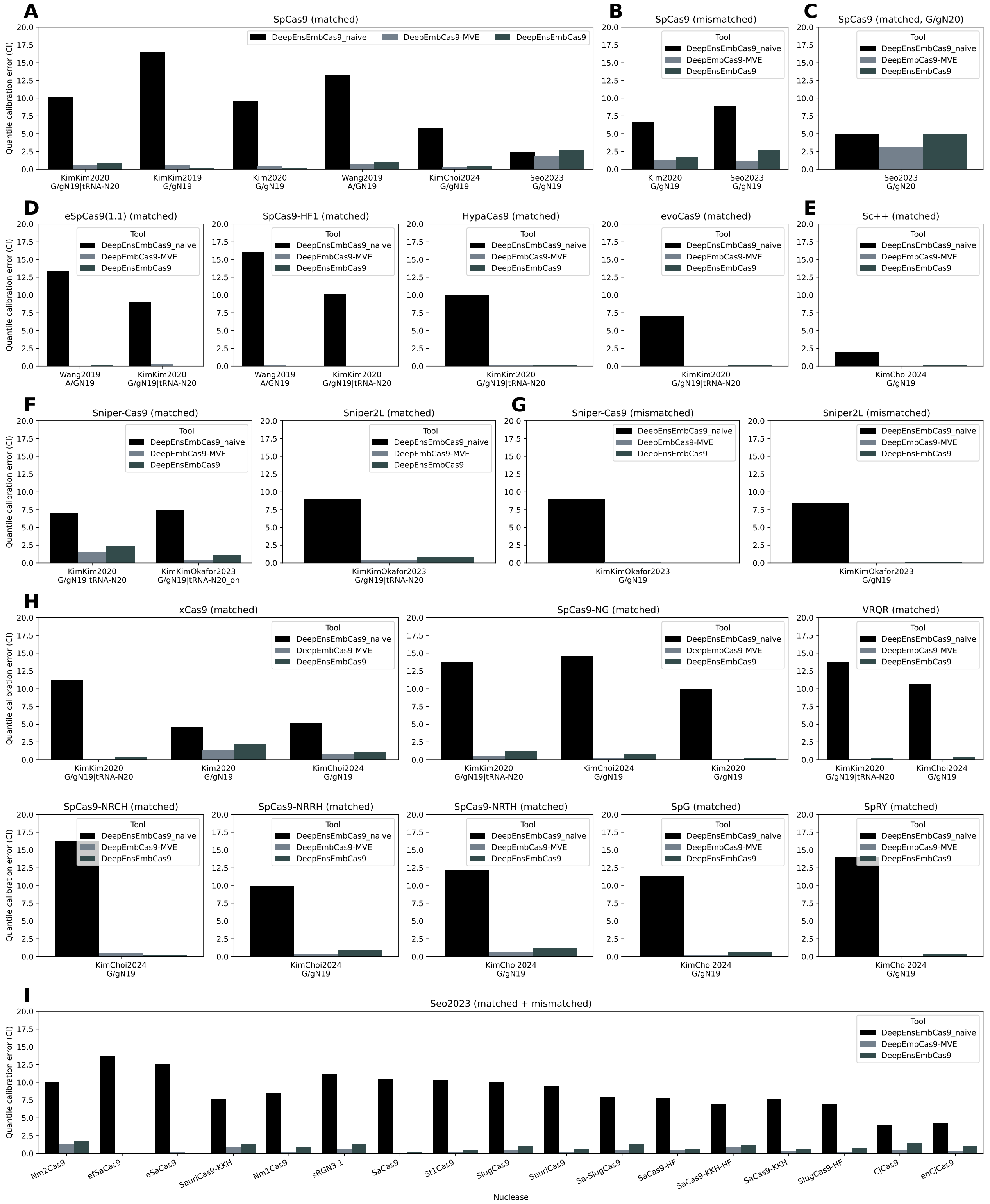


Figure S37: Confidence interval-based quantile calibration errors for DeepEnsEmbCas9_naive, DeepEmbCas9-MVE and DeepEnsEmbCas9, conditioned on (A) matched AN$_{19}$, G/gN$_{19}$ and tRNA$^{Gln}$-N$_{20}$ wild type SpCas9 interfaces; (B) mismatched G/gN$_{19}$ and matched G/gN$_{20}$ wild type SpCas9 interfaces; (C) matched AN$_{19}$, G/gN$_{19}$ and tRNA$^{Gln}$-N$_{20}$ eSpCas9(1.1) interfaces; (D) matched AN$_{19}$, G/gN$_{19}$ and tRNA$^{Gln}$-N$_{20}$ SpCas9-HF1 interfaces; for matched G/gN$_{19}$ and tRNA$^{Gln}$-N$_{20}$ HypaCas9/evoCas9 and G/gN$_{19}$ Sc++ interfaces; (F) matched G/gN$_{19}$ and tRNA$^{Gln}$-N$_{20}$ and mismatched G/gN$_{19}$ interfaces for 2 Sniper variants; (G,H) matched G/gN$_{19}$ and tRNA$^{Gln}$-N$_{20}$ interfaces for xCas9/SpCas9-NG (G) and 6 other PAM-altered SpCas9 variants (H); and (I) matched and mismatched interfaces for 17 wild type or engineered small Cas9 nucleases.

#### 2.6 Per-nuclease extrapolation calibration

##### 2.6.1 Quantile calibration plots - DeepEnsEmbCas9_omit

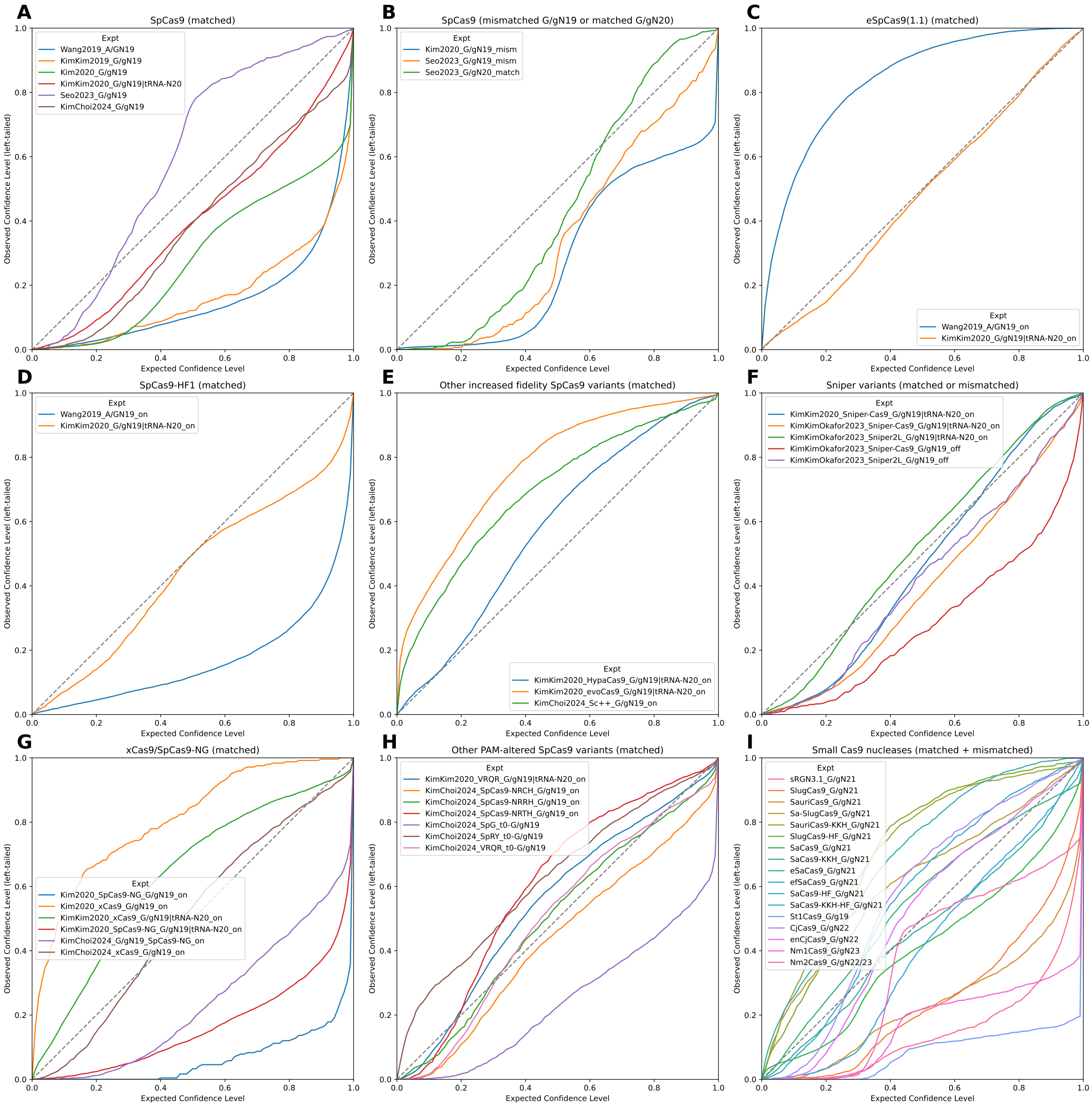


Figure S38: Quantile calibration plots for DeepEnsEmbCas9_omit, conditioned on (A) matched AN$_{19}$, G/gN$_{19}$ and tRNA$^{Gln}$-N$_{20}$ wild type SpCas9 interfaces; (B) mismatched G/gN$_{19}$ and matched G/gN$_{20}$ wild type SpCas9 interfaces; (C) matched AN$_{19}$, G/gN$_{19}$ and tRNA$^{Gln}$-N$_{20}$ eSpCas9(1.1) interfaces; (D) matched AN$_{19}$, G/gN$_{19}$ and tRNA$^{Gln}$-N$_{20}$ SpCas9-HF1 interfaces; for matched G/gN$_{19}$ and tRNA$^{Gln}$-N$_{20}$ HypaCas9/evoCas9 and G/gN$_{19}$ Sc++ interfaces; (F) matched G/gN$_{19}$ and tRNA$^{Gln}$-N$_{20}$ and mismatched G/gN$_{19}$ interfaces for 2 Sniper variants; (G,H) matched G/gN$_{19}$ and tRNA$^{Gln}$-N$_{20}$ interfaces for xCas9/SpCas9-NG (G) and 6 other PAM-altered SpCas9 variants (H); and (I) matched and mismatched interfaces for 17 wild type or engineered small Cas9 nucleases.

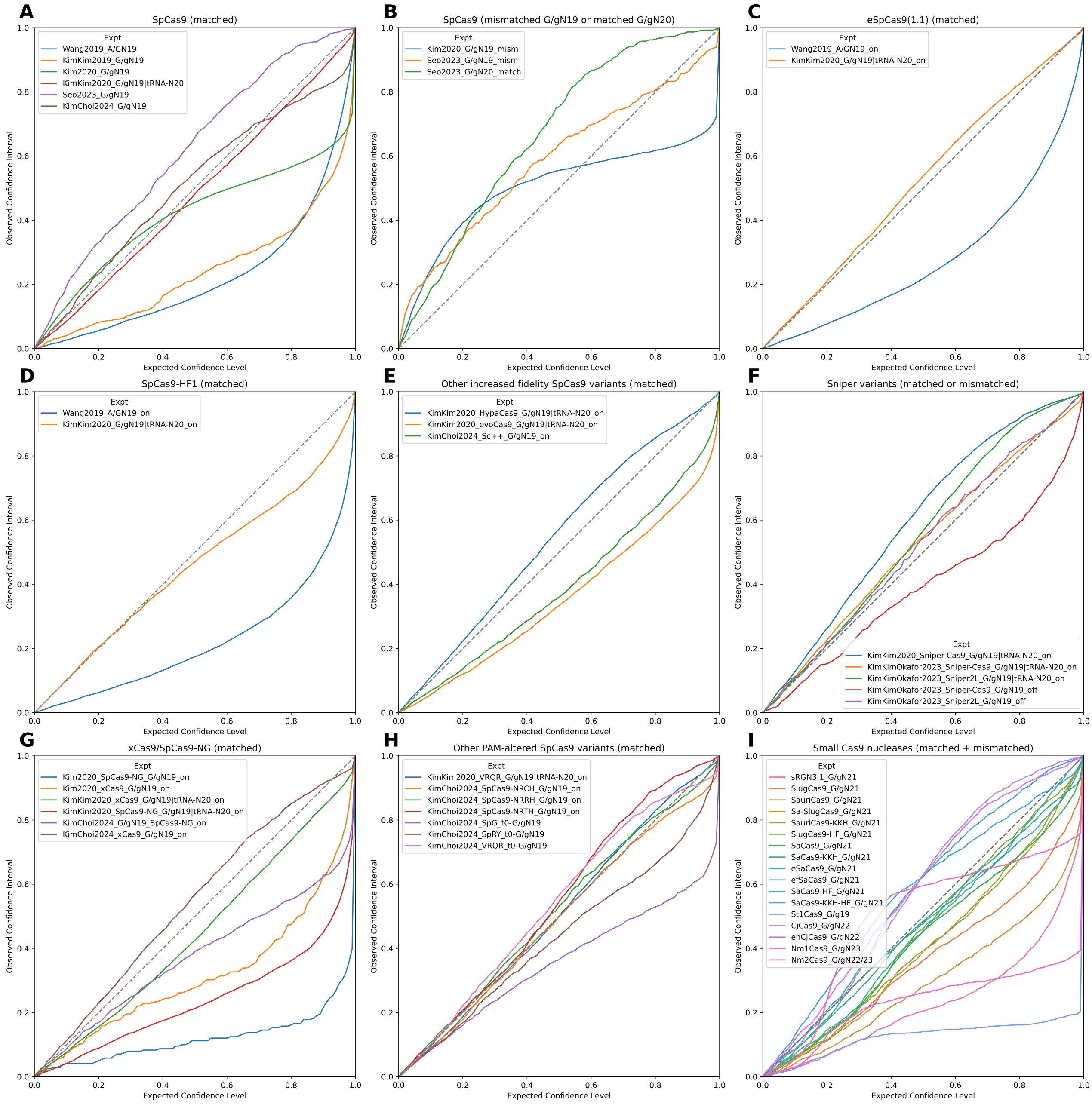


Figure S39: Confidence interval-based calibration curves for DeepEnsEmbCas9_omit, conditioned on (A) matched $AN_{19}$, $G/gN_{19}$ and $tRNA^{Gln}$-$N_{20}$ wild type SpCas9 interfaces; (B) mismatched $G/gN_{19}$ and matched $G/gN_{20}$ wild type SpCas9 interfaces; (C) matched $AN_{19}$, $G/gN_{19}$ and $tRNA^{Gln}$-$N_{20}$ eSpCas9(1.1) interfaces; (D) matched $AN_{19}$, $G/gN_{19}$ and $tRNA^{Gln}$-$N_{20}$ SpCas9-HF1 interfaces; for matched $G/gN_{19}$ and $tRNA^{Gln}$-$N_{20}$ HypaCas9/evoCas9 and $G/gN_{19}$ Sc++ interfaces; (F) matched $G/gN_{19}$ and $tRNA^{Gln}$-$N_{20}$ and mismatched $G/gN_{19}$ interfaces for 2 Sniper variants; (G,H) matched $G/gN_{19}$ and $tRNA^{Gln}$-$N_{20}$ interfaces for xCas9/SpCas9-NG (G) and 6 other PAM-altered SpCas9 variants (H); and (I) matched and mismatched interfaces for 17 wild type or engineered small Cas9 nucleases.

##### 2.6.2 Quantile calibration plots - DeepEmbCas9-MVE_omit

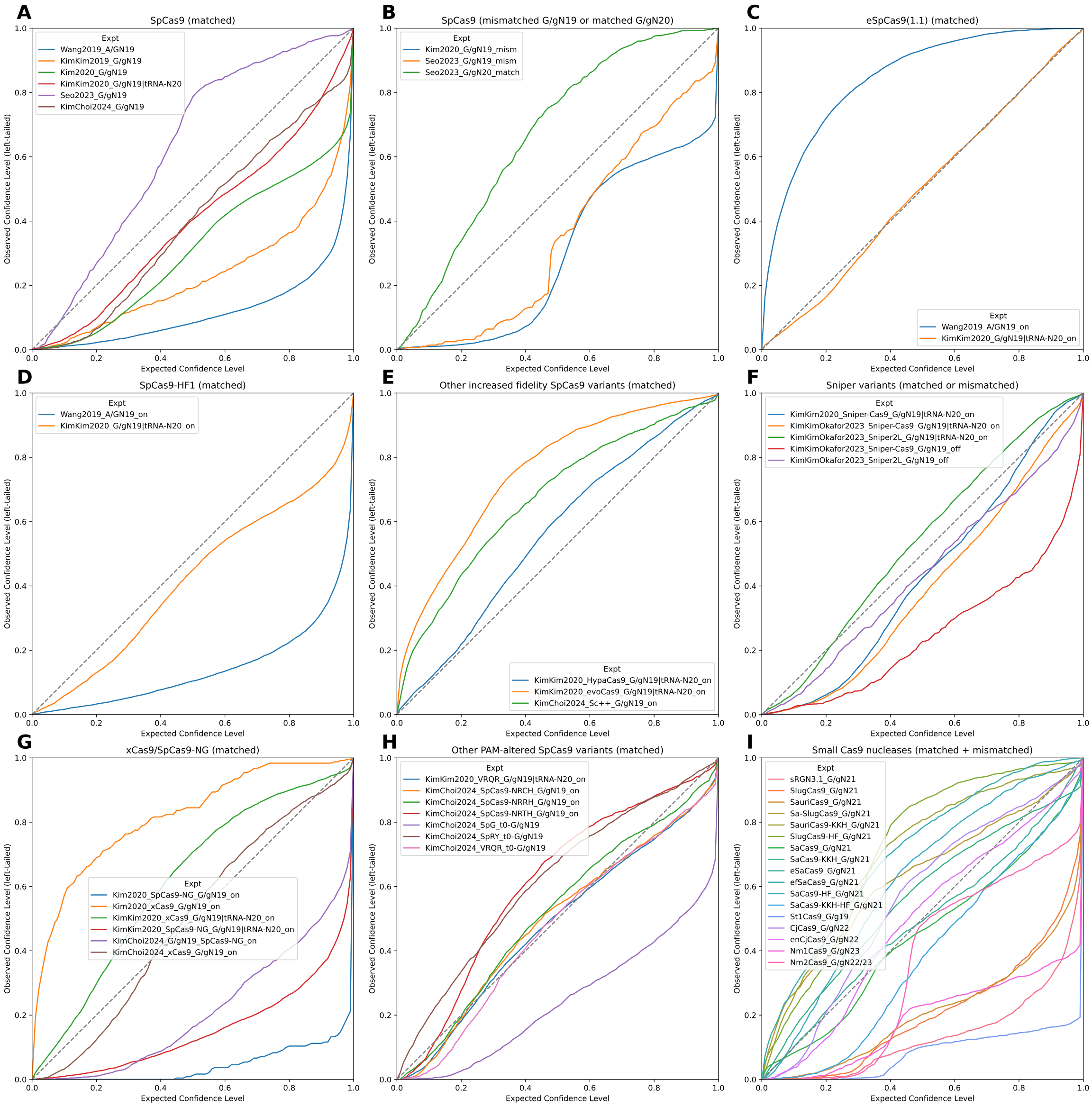


Figure S40: Quantile calibration plots for DeepEmbCas9-MVE_omit, conditioned on (A) matched $AN_{19}$, $G/gN_{19}$ and $tRNA^{Gln}$-$N_{20}$ wild type SpCas9 interfaces; (B) mismatched $G/gN_{19}$ and matched $G/gN_{20}$ wild type SpCas9 interfaces; (C) matched $AN_{19}$, $G/gN_{19}$ and $tRNA^{Gln}$-$N_{20}$ eSpCas9(1.1) interfaces; (D) matched $AN_{19}$, $G/gN_{19}$ and $tRNA^{Gln}$-$N_{20}$ SpCas9-HF1 interfaces; for matched $G/gN_{19}$ and $tRNA^{Gln}$-$N_{20}$ HypaCas9/evoCas9 and $G/gN_{19}$ Sc++ interfaces; (F) matched $G/gN_{19}$ and $tRNA^{Gln}$-$N_{20}$ and mismatched $G/gN_{19}$ interfaces for 2 Sniper variants; (G,H) matched $G/gN_{19}$ and $tRNA^{Gln}$-$N_{20}$ interfaces for xCas9/SpCas9-NG (G) and 6 other PAM-altered SpCas9 variants (H); and (I) matched and mismatched interfaces for 17 wild type or engineered small Cas9 nucleases.

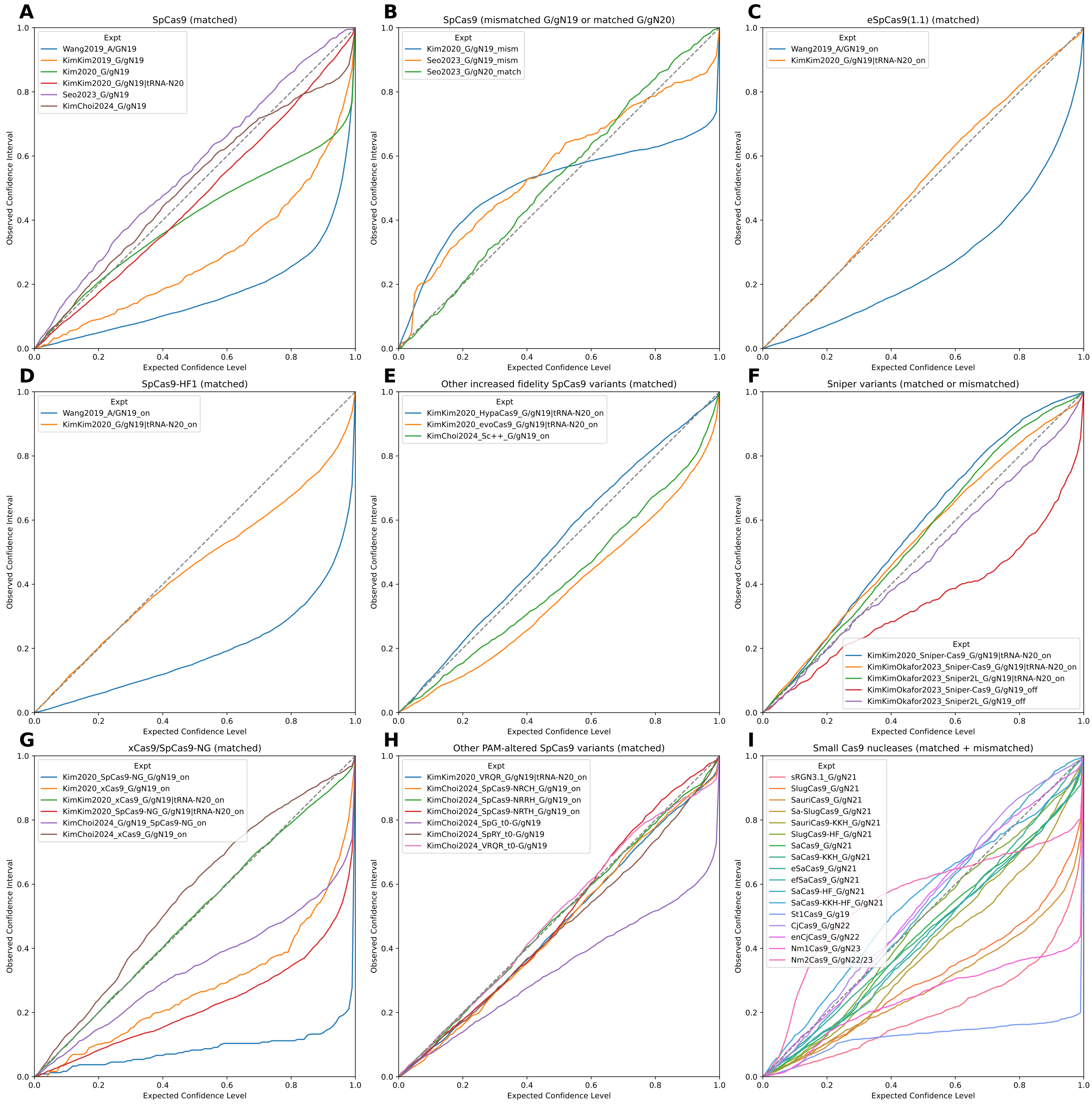


Figure S41: Confidence interval-based calibration curves for DeepEmbCas9-MVE_omit, conditioned on (A) matched $AN_{19}$, $G/gN_{19}$ and $tRNA^{Gln}$-$N_{20}$ wild type SpCas9 interfaces; (B) mismatched $G/gN_{19}$ and matched $G/gN_{20}$ wild type SpCas9 interfaces; (C) matched $AN_{19}$, $G/gN_{19}$ and $tRNA^{Gln}$-$N_{20}$ eSpCas9(1.1) interfaces; (D) matched $AN_{19}$, $G/gN_{19}$ and $tRNA^{Gln}$-$N_{20}$ SpCas9-HF1 interfaces; for matched $G/gN_{19}$ and $tRNA^{Gln}$-$N_{20}$ HypaCas9/evoCas9 and $G/gN_{19}$ Sc++ interfaces; (F) matched $G/gN_{19}$ and $tRNA^{Gln}$-$N_{20}$ and mismatched $G/gN_{19}$ interfaces for 2 Sniper variants; (G,H) matched $G/gN_{19}$ and $tRNA^{Gln}$-$N_{20}$ interfaces for xCas9/SpCas9-NG (G) and 6 other PAM-altered SpCas9 variants (H); and (I) matched and mismatched interfaces for 17 wild type or engineered small Cas9 nucleases.

##### 2.6.3 Quantile calibration plots - DeepEnsEmbCas9_naive_omit

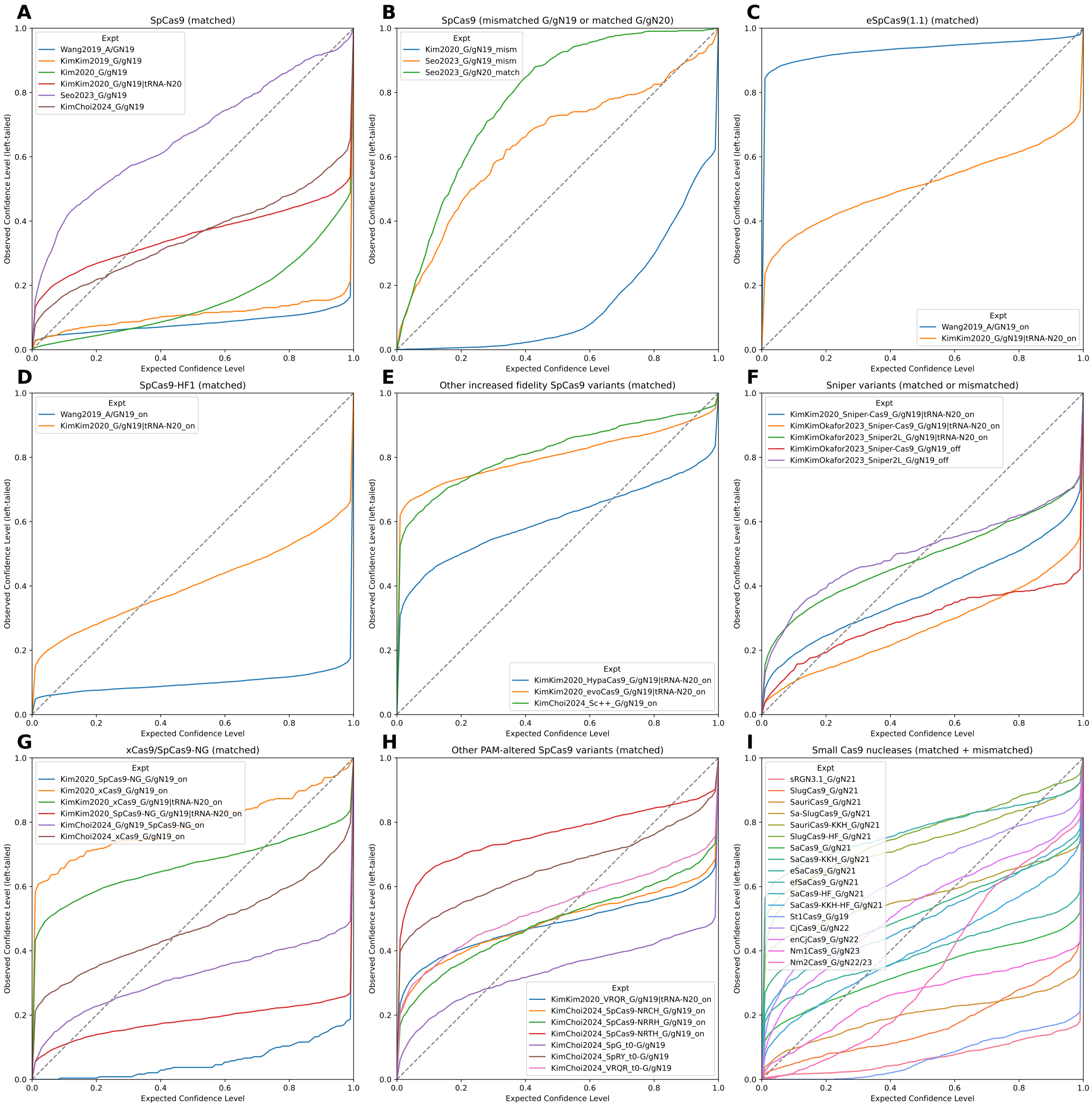

Figure S42: Quantile calibration plots for DeepEnsEmbCas9_naive_omit, conditioned on (A) matched $AN_{19}$, $G/gN_{19}$ and $tRNA^{Gln}$-$N_{20}$ wild type SpCas9 interfaces; (B) mismatched $G/gN_{19}$ and matched $G/gN_{20}$ wild type SpCas9 interfaces; (C) matched $AN_{19}$, $G/gN_{19}$ and $tRNA^{Gln}$-$N_{20}$ eSpCas9(1.1) interfaces; (D) matched $AN_{19}$, $G/gN_{19}$ and $tRNA^{Gln}$-$N_{20}$ SpCas9-HF1 interfaces; for matched $G/gN_{19}$ and $tRNA^{Gln}$-$N_{20}$ HypaCas9/evoCas9 and $G/gN_{19}$ Sc++ interfaces; (F) matched $G/gN_{19}$ and $tRNA^{Gln}$-$N_{20}$ and mismatched $G/gN_{19}$ interfaces for 2 Sniper variants; (G,H) matched $G/gN_{19}$ and $tRNA^{Gln}$-$N_{20}$ interfaces for xCas9/SpCas9-NG (G) and 6 other PAM-altered SpCas9 variants (H); and (I) matched and mismatched interfaces for 17 wild type or engineered small Cas9 nucleases.

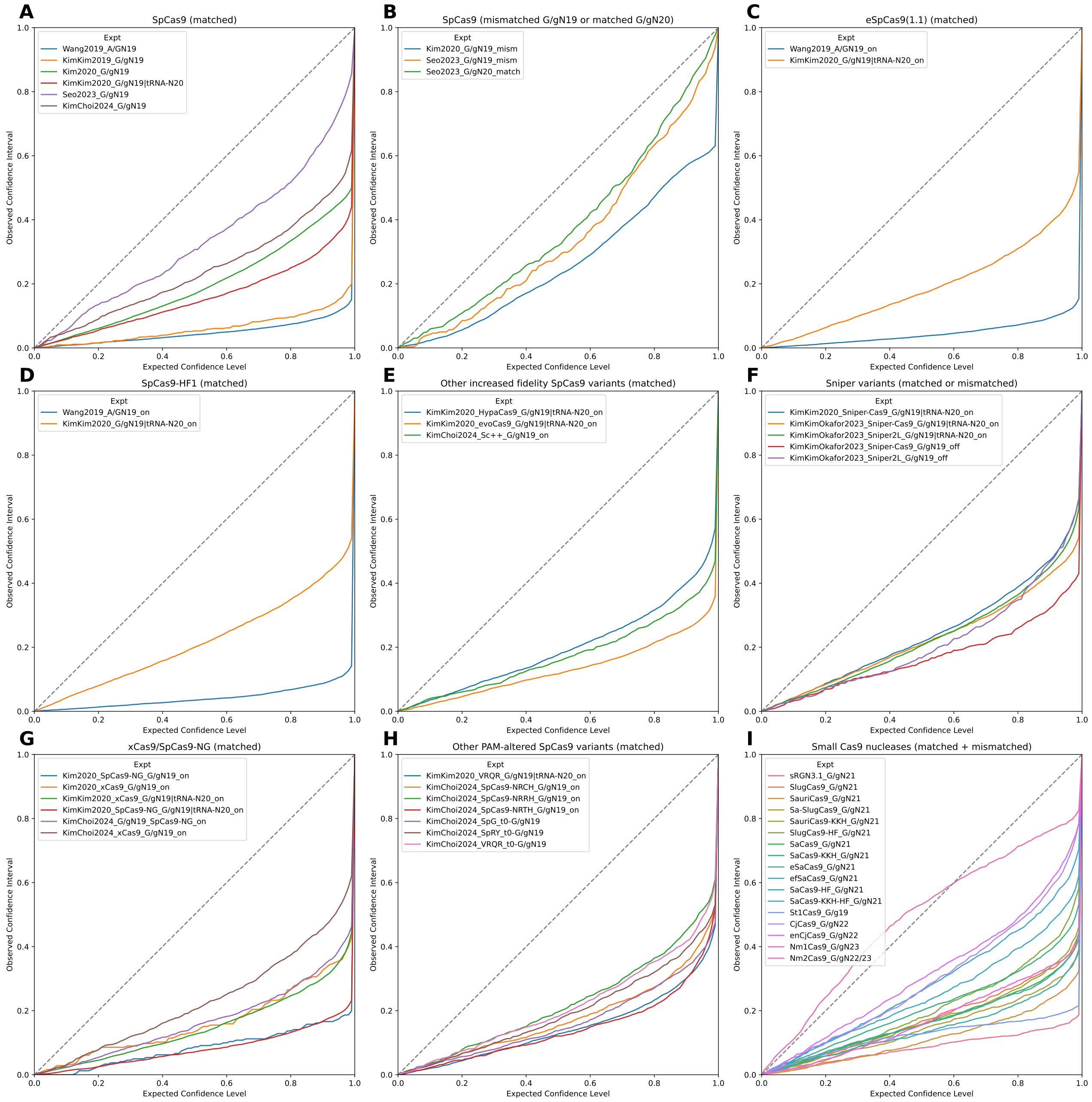


Figure S43: Confidence interval-based calibration curves for DeepEnsEmbCas9_naive_omit, conditioned on (A) matched $AN_{19}$, $G/gN_{19}$ and $tRNA^{Gln}$-$N_{20}$ wild type SpCas9 interfaces; (B) mismatched $G/gN_{19}$ and matched $G/gN_{20}$ wild type SpCas9 interfaces; (C) matched $AN_{19}$, $G/gN_{19}$ and $tRNA^{Gln}$-$N_{20}$ eSpCas9(1.1) interfaces; (D) matched $AN_{19}$, $G/gN_{19}$ and $tRNA^{Gln}$-$N_{20}$ SpCas9-HF1 interfaces; for matched $G/gN_{19}$ and $tRNA^{Gln}$-$N_{20}$ HypaCas9/evoCas9 and $G/gN_{19}$ Sc++ interfaces; (F) matched $G/gN_{19}$ and $tRNA^{Gln}$-$N_{20}$ and mismatched $G/gN_{19}$ interfaces for 2 Sniper variants; (G,H) matched $G/gN_{19}$ and $tRNA^{Gln}$-$N_{20}$ interfaces for xCas9/SpCas9-NG (G) and 6 other PAM-altered SpCas9 variants (H); and (I) matched and mismatched interfaces for 17 wild type or engineered small Cas9 nucleases.

##### 2.6.4 Quantile calibration error

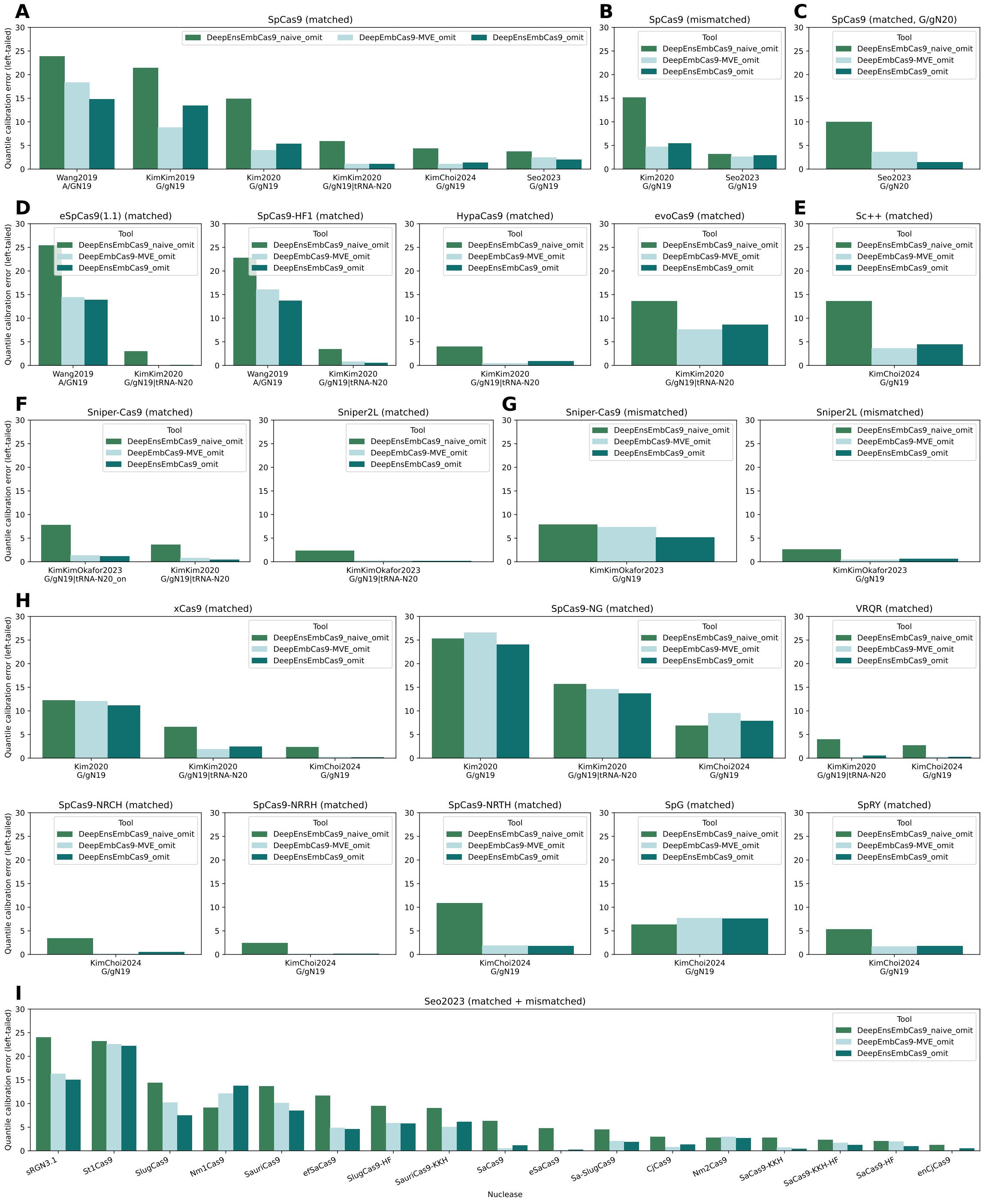

Figure S44: Quantile calibration errors for DeepEnsEmbCas9_naive_omit, DeepEmbCas9-MVE_omit and DeepEnsEmbCas9_omit, conditioned on (A) matched $AN_{19}$, $G/gN_{19}$ and $tRNA^{Gln}$-$N_{20}$ wild type SpCas9 interfaces; (B) mismatched $G/gN_{19}$ and matched $G/gN_{20}$ wild type SpCas9 interfaces; (C) matched $AN_{19}$, $G/gN_{19}$ and $tRNA^{Gln}$-$N_{20}$ eSpCas9(1.1) interfaces; (D) matched $AN_{19}$, $G/gN_{19}$ and $tRNA^{Gln}$-$N_{20}$ SpCas9-HF1 interfaces; for matched $G/gN_{19}$ and $tRNA^{Gln}$-$N_{20}$ HypaCas9/evoCas9 and $G/gN_{19}$ Sc++ interfaces; (F) matched $G/gN_{19}$ and $tRNA^{Gln}$-$N_{20}$ and mismatched $G/gN_{19}$ interfaces for 2 Sniper variants; (G,H) matched $G/gN_{19}$ and $tRNA^{Gln}$-$N_{20}$ interfaces for xCas9/SpCas9-NG (G) and 6 other PAM-altered SpCas9 variants (H); and (I) matched and mismatched interfaces for 17 wild type or engineered small Cas9 nucleases.

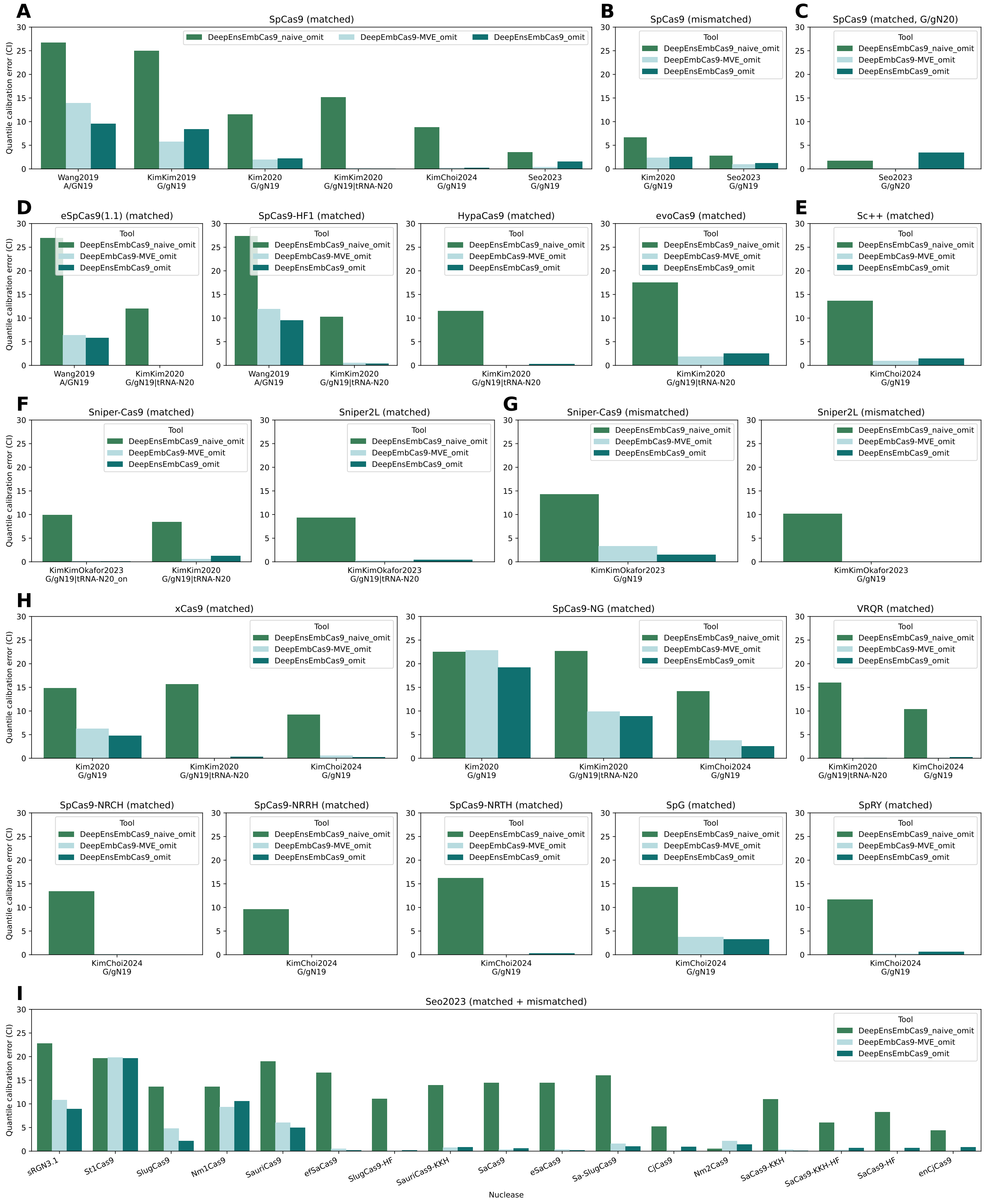


Figure S45: Confidence interval-based quantile calibration errors for DeepEnsEmbCas9_naive_omit, DeepEmbCas9-MVE_omit and DeepEnsEmbCas9_omit, conditioned on (A) matched $AN_{19}$, $G/gN_{19}$ and $tRNA^{Gln}$-$N_{20}$ wild type SpCas9 interfaces; (B) mismatched $G/gN_{19}$ and matched $G/gN_{20}$ wild type SpCas9 interfaces; (C) matched $AN_{19}$, $G/gN_{19}$ and $tRNA^{Gln}$-$N_{20}$ eSpCas9(1.1) interfaces; (D) matched $AN_{19}$, $G/gN_{19}$ and $tRNA^{Gln}$-$N_{20}$ SpCas9-HF1 interfaces; for matched $G/gN_{19}$ and $tRNA^{Gln}$-$N_{20}$ HypaCas9/evoCas9 and $G/gN_{19}$ Sc++ interfaces; (F) matched $G/gN_{19}$ and $tRNA^{Gln}$-$N_{20}$ and mismatched $G/gN_{19}$ interfaces for 2 Sniper variants; (G,H) matched $G/gN_{19}$ and $tRNA^{Gln}$-$N_{20}$ interfaces for xCas9/SpCas9-NG (G) and 6 other PAM-altered SpCas9 variants (H); and (I) matched and mismatched interfaces for 17 wild type or engineered small Cas9 nucleases.

#### 2.7 Model Interpretation

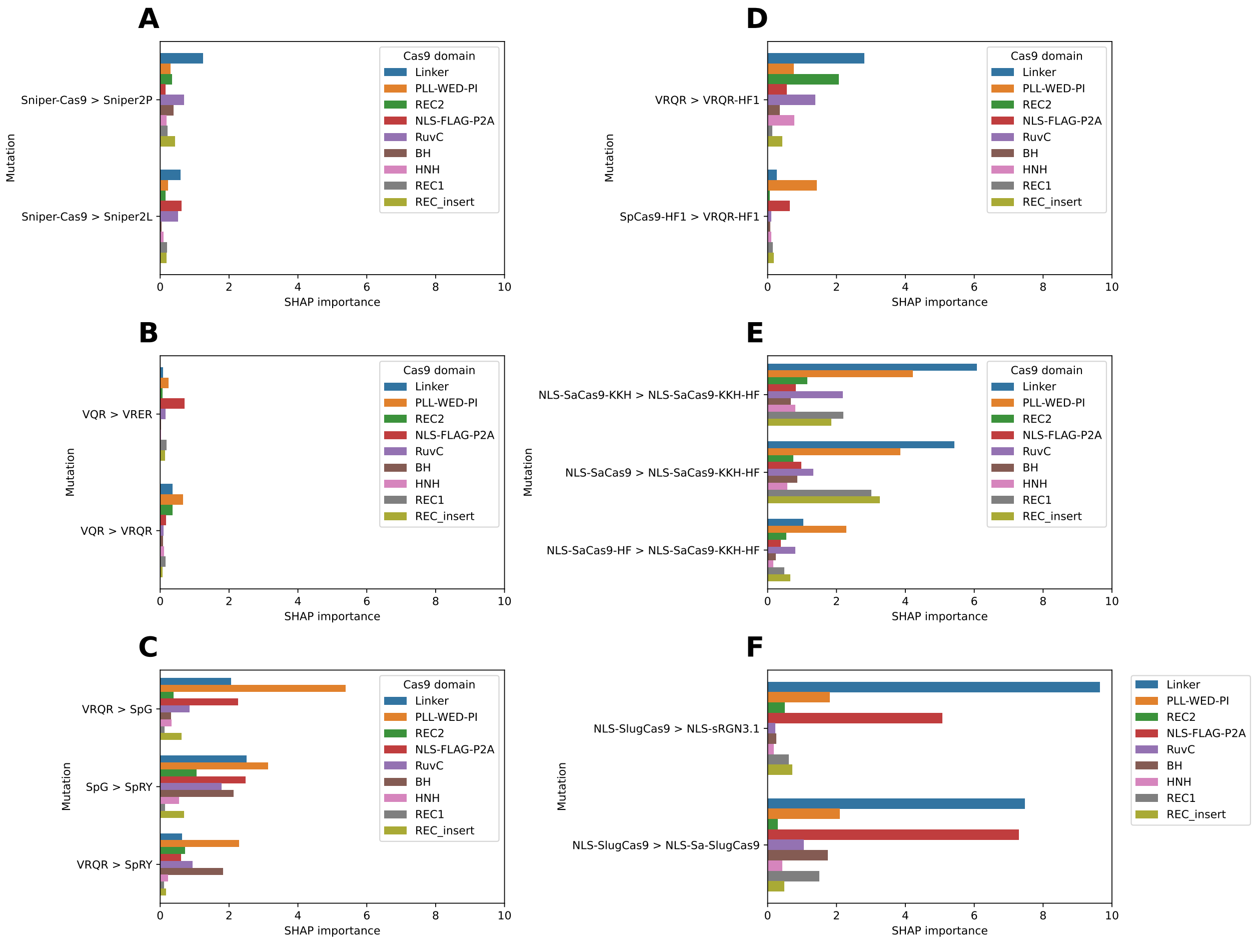


Figure S46: Important Cas9 domains driving DeepEmbCas9's change in predicted activity when (A) mutating Sniper-Cas9 into Sniper2 variants; (B) mutating VQR into VRER and VRQR; (C) mutating into PAM-relaxed/PAMless SpCas9 variants; (D) mutating to VRQR-HF1 from VRQR and SpCas9-HF1; (E) mutating SaCas9-KKH, SaCas9 and SaCas9-HF to SaCas9-KKH-HF; and (F) mutating SlugCas9 into sRGN3.1 and Sa-SlugCas9.

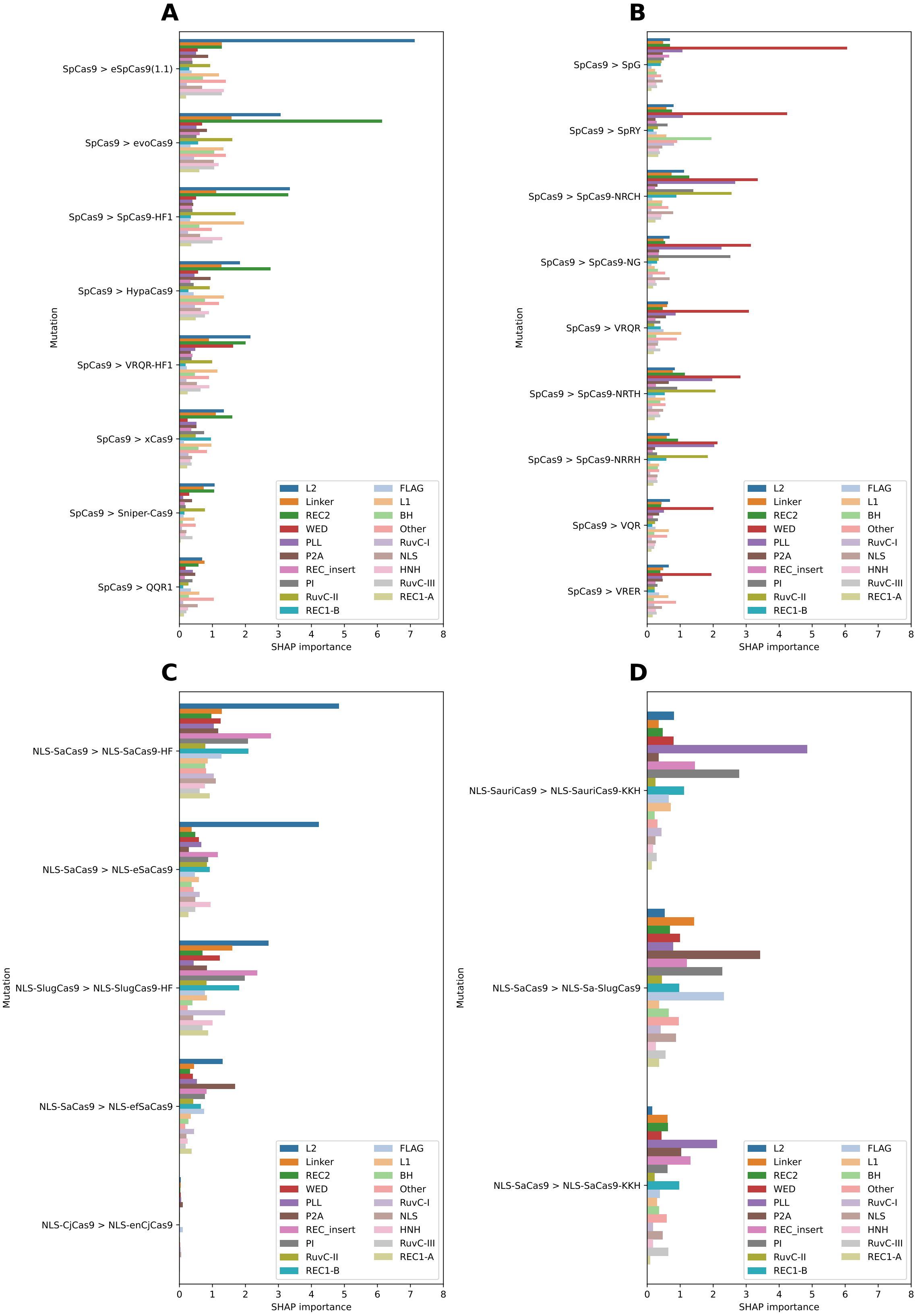


Figure S47: CRISPR-Cas9 Cas9 regions driving DeepEmbCas9's change in predicted activity when introducing residue mutations in Cas9. (C,D) Cas9 region importances for SpCas9 variants with (C) and without (D) D1135 mutations. (E,F) Cas9 region importances for small Cas9 variants with (E) increased fidelity and (F) PAM-altering variants.

| Feature group | Description of features | No. of features |
|---|---|---|
| spacer + spacer_MFE + spacer_GCcount | spacer one-hot encoding | $4 \times 42$ |
| | spacer MFE | 1 |
| | spacer GC count | 1 |
| | sgRNA rLM embedding for spacer region | 768 |
| upstream + protospacer + protospacer_Tm | 5' upstream and protospacer target one-hot encoding | $4 \times 27$ |
| | protospacer DNA melting temperatures | 21 |
| | protospacer GC count | 1 |
| PAM + downstream + PAM_downstream_Tm | PAM and 3' downstream one-hot encoding | $4 \times 15$ |
| | PAM melting temperatures | 6 |
| | 3' downstream DNA melting temperatures | 9 |
| tRNA preprocessing | tRNA feature | 1 |
| Day | Day | 1 |
| Cas9_ESM-C-600M_RuvC-I | Cas9 pLM embedding features for the RuvC-I region | 960 |
| Cas9_ESM-C-600M_BH | Cas9 pLM embedding features for BH region | 960 |
| Cas9_ESM-C-600M_REC1-A | Cas9 pLM embedding features for the REC1-A region | 960 |
| Cas9_ESM-C-600M_REC_insert | Cas9 pLM embedding features for the REC_insert region | 960 |
| Cas9_ESM-C-600M_REC1-B | Cas9 pLM embedding features for the REC1-B region | 960 |
| Cas9_ESM-C-600M_REC2 | Cas9 pLM embedding features for the REC2 region | 960 |
| Cas9_ESM-C-600M_Linker | Cas9 pLM embedding features for the Linker region | 960 |
| Cas9_ESM-C-600M_RuvC-II | Cas9 pLM embedding features for the RuvC-II region | 960 |
| Cas9_ESM-C-600M_L1 | Cas9 pLM embedding features for the L1 region | 960 |
| Cas9_ESM-C-600M_HNH | Cas9 pLM embedding features for the HNH region | 960 |
| Cas9_ESM-C-600M_L2 | Cas9 pLM embedding features for the L2 region | 960 |
| Cas9_ESM-C-600M_RuvC-III | Cas9 pLM embedding features for the RuvC-III region | 960 |
| Cas9_ESM-C-600M_PLL | Cas9 pLM embedding features for the PLL region | 960 |
| Cas9_ESM-C-600M_WED | Cas9 pLM embedding features for the WED region | 960 |
| Cas9_ESM-C-600M_PI | Cas9 pLM embedding features for the PI region | 960 |
| Cas9_ESM-C-600M_NLS | Cas9 pLM embedding features for the NLS region | 960 |
| Cas9_ESM-C-600M_FLAG | Cas9 pLM embedding features for the FLAG region | 960 |
| Cas9_ESM-C-600M_P2A | Cas9 pLM embedding features for the P2A region | 960 |
| Cas9_ESM-C-600M_Other | Cas9 pLM embedding features for the Other region | 960 |
| sgRNA_BEACON-B_repeat-antirepeat | sgRNA rLM embedding features for the repeat-antirepeat region | 960 |
| sgRNA_BEACON-B_tracrRNA-rest | sgRNA rLM embedding features for the repeat-antirepeat region | 960 |
| sgRNA_BEACON-B_polyT | sgRNA rLM embedding features for the polyT region | 960 |

Table S7: List and descriptions of fine resolution CRISPR-Cas9 complex component feature groups used in SHAP importance analysis.

| Feature Group | Features | No. of features |
|---|---|---|
| spacer | spacer one-hot encoding | $4 \times 42$ |
| | spacer MFE | 1 |
| | $\frac{1}{2}\times$ sgRNA MFE | 0.5 |
| | spacer GC count | 1 |
| | sgRNA rLM embedding for spacer region | 768 |
| target | target context sequence one-hot encoding | $4 \times 42$ |
| | DNA melting temperature features | 36 |
| | protospacer GC count | 36 |
| Cas9 | Cas9 pLM embedding for all Cas9 regions | |
| scaffold | sgRNA rLM embedding for repeat-antirepeat region | 768 |
| | sgRNA rLM embedding for tracrRNA-rest region | 768 |
| | sgRNA rLM embedding for polyT region | 768 |
| | $\frac{1}{2}\times$ sgRNA MFE | 0.5 |
| tRNA preprocessing | tRNA feature | 1 |
| Day | Day | 1 |

Table S8: List and descriptions of coarse resolution CRISPR-Cas9 complex component feature groups used in SHAP importance analysis.

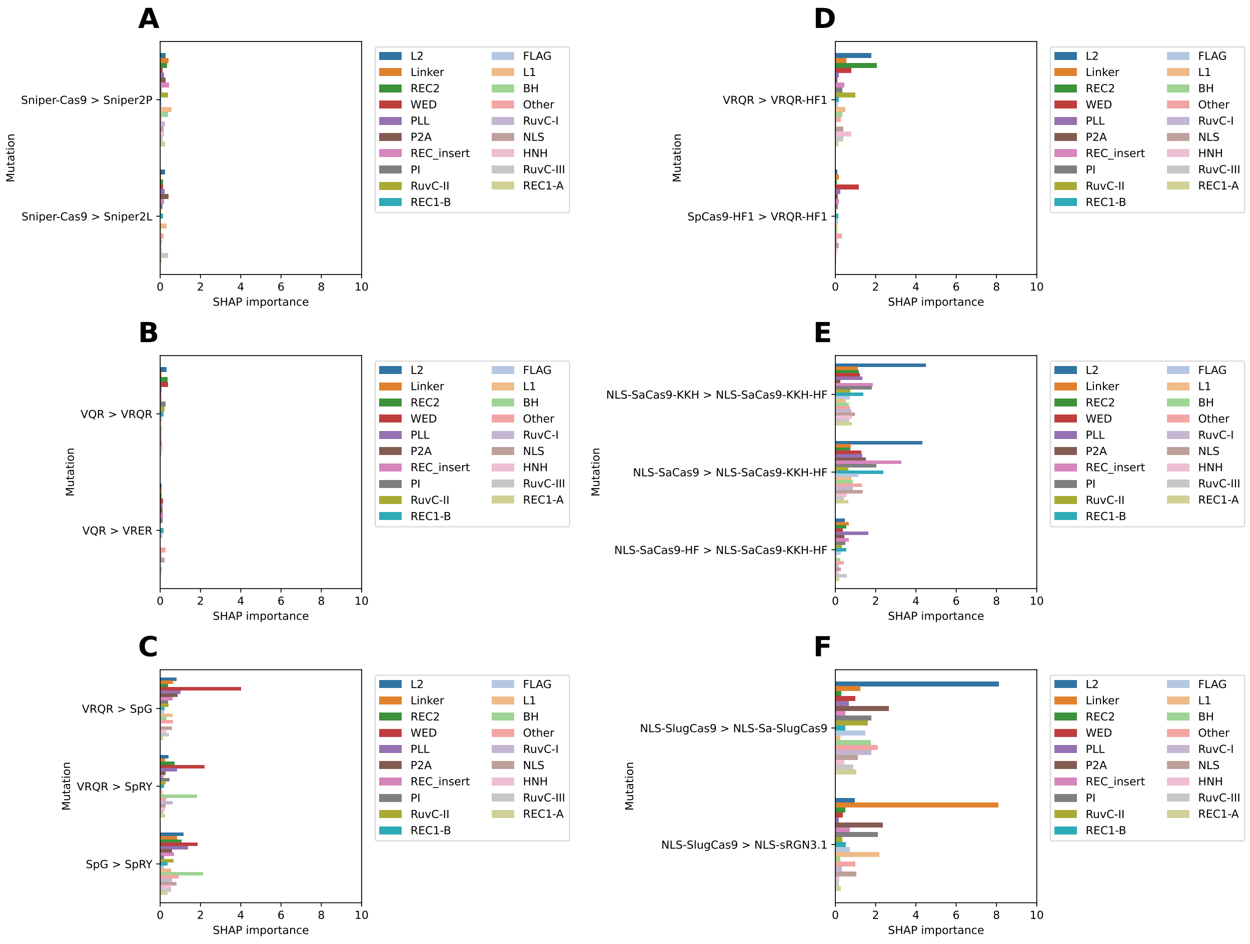


Figure S48: Important Cas9 regions driving DeepEmbCas9's change in predicted activity when (A) mutating Sniper-Cas9 into Sniper2 variants; (B) mutating VQR into VRER and VRQR; (C) mutating into PAM-relaxed/PAMless SpCas9 variants; (D) mutating to VRQR-HF1 from VRQR and SpCas9-HF1; (E) mutating SaCas9-KKH, SaCas9 and SaCas9-HF to SaCas9-KKH-HF; and (F) mutating SlugCas9 into sRGN3.1 and Sa-SlugCas9.

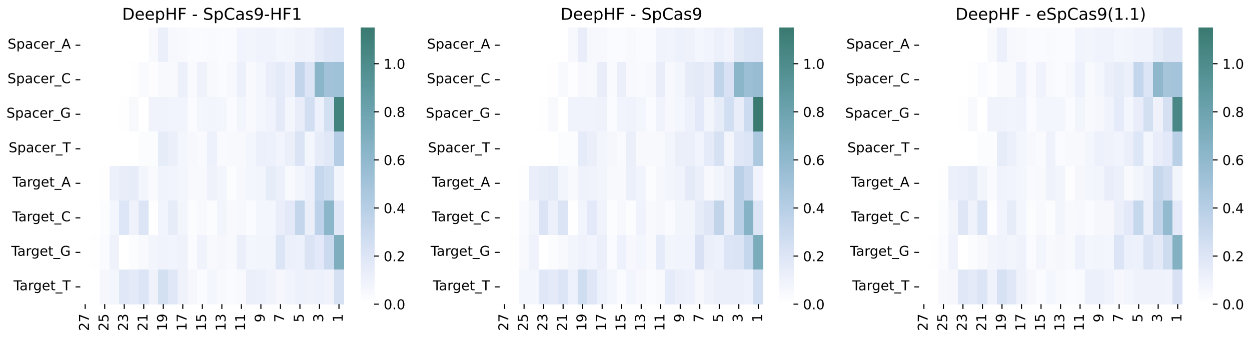


Figure S49: SHAP importance of spacer-target one-hot encoding features in driving DeepEmbCas9's change in predicted activity for Cas9 variants from Wang et al. [2].

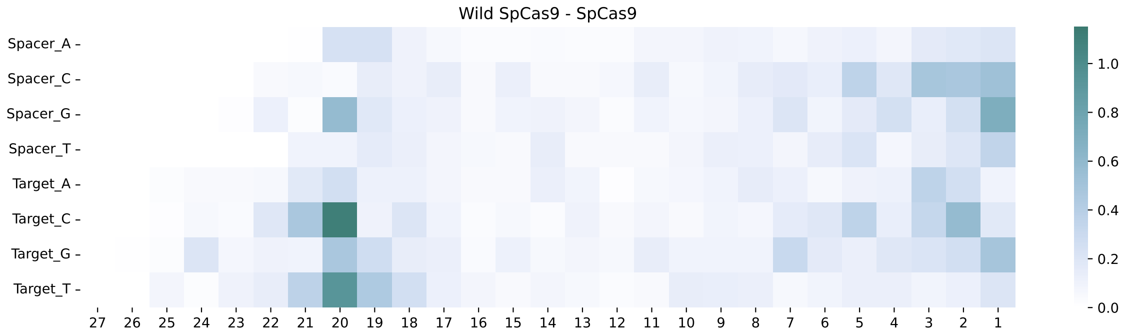


Figure S50: SHAP importance of spacer-target one-hot encoding features in driving DeepEmbCas9's change in predicted activity for Cas9 variants from Kim, Kim et al. [1].

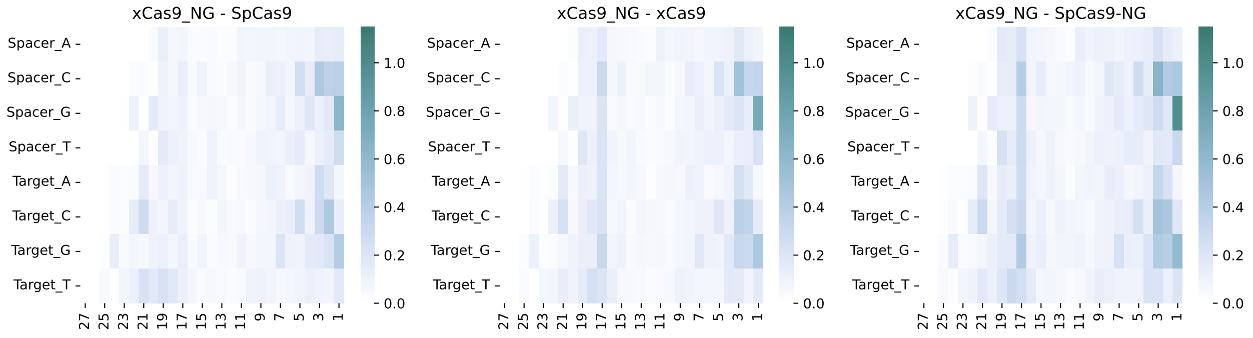


Figure S51: SHAP importance of spacer-target one-hot encoding features in driving DeepEmbCas9's change in predicted activity for Cas9 variants from Kim et al. [3].

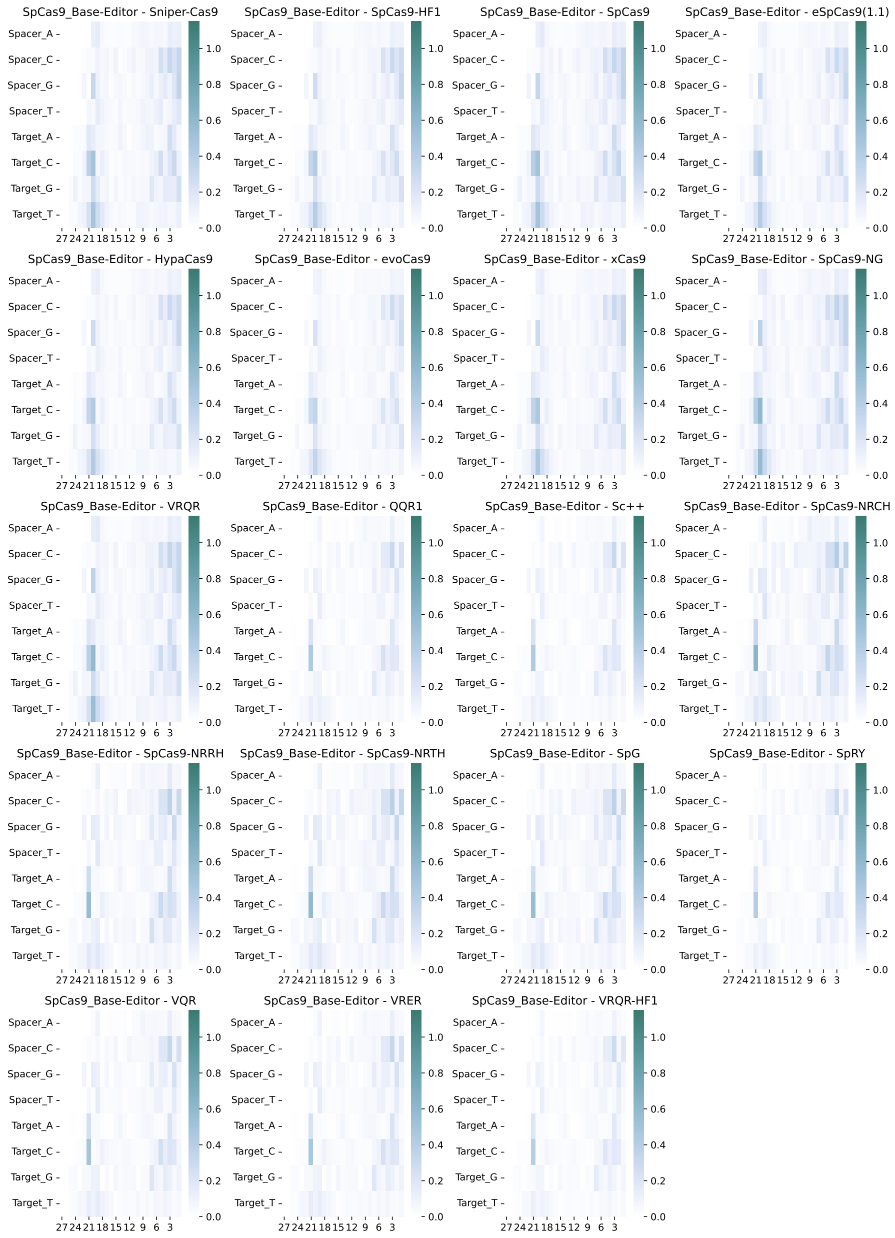


Figure S52: SHAP importance of spacer-target one-hot encoding features in driving DeepEmbCas9's change in predicted activity for Cas9 variants from Kim, Kim et al. [4] and Kim, Choi et al. [7].

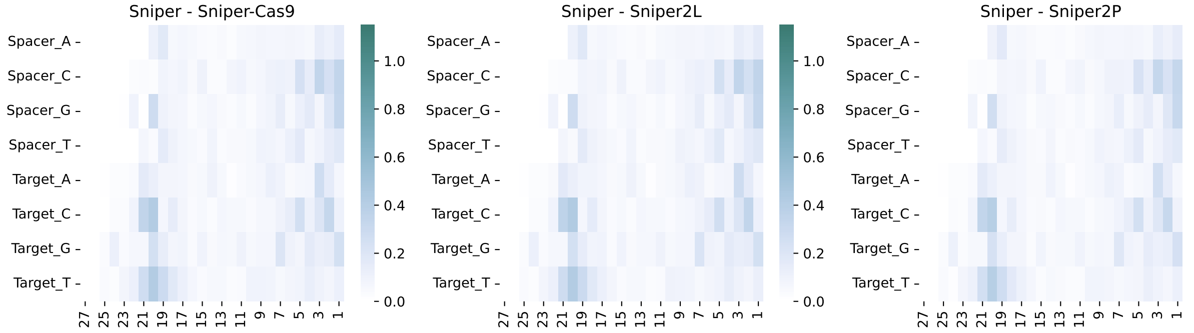


Figure S53: SHAP importance of spacer-target one-hot encoding features in driving DeepEmbCas9's change in predicted activity for Cas9 variants from Kim, Kim, Okafor et al. [7].

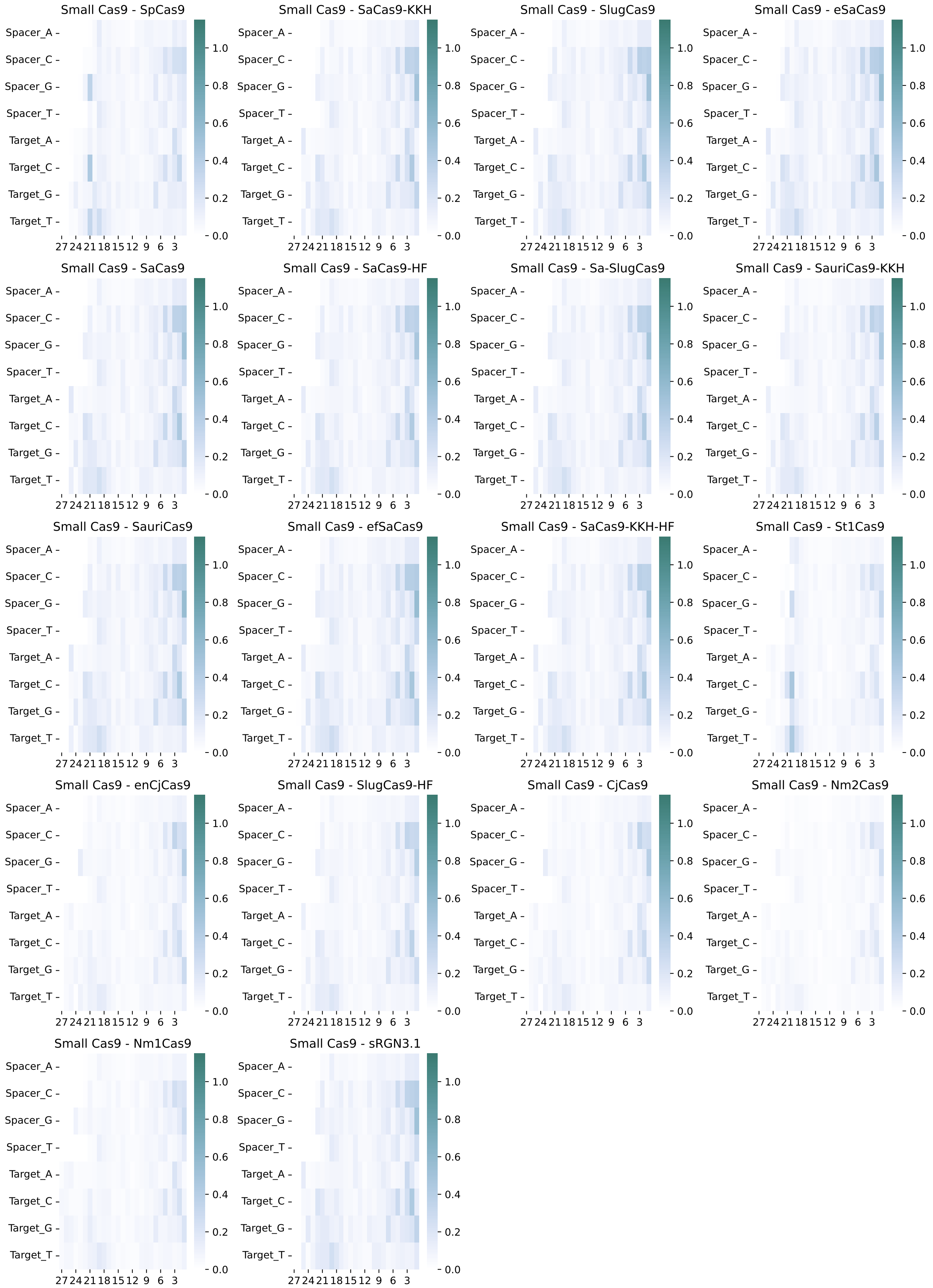


Figure S54: SHAP importance of spacer-target one-hot encoding features in driving DeepEmbCas9's change in predicted activity for Cas9 variants from Kim, Kim, Okafor et al. [7].
